# Continual integration of single-cell multimodal data with MIRACLE

**DOI:** 10.1101/2024.09.24.613833

**Authors:** Jiahao Zhou, Zhen He, Jing Wang, Shuofeng Hu, Tongtong Kan, Guohua Dong, Jinhui Shi, Runyan Liu, Le Ou-Yang, Xiaochen Bo, Xiaomin Ying

**Affiliations:** Center for Computational Biology, Beijing Institute of Basic Medical Sciences, Beijing, China; College of Electronics and Information Engineering, Shenzhen University, Shenzhen, China; Institute of Health Service and Transfusion Medicine, Beijing, China

## Abstract

Single-cell sequencing technologies have revolutionized our understanding of cellular heterogeneity and facilitated the construction of multi-omics cell atlases via data integration. However, updating these atlases with new data conventionally requires reintegration of all data and is computationally intensive, hindering timely updates and dynamic adjustments in biological and medical research. To address this challenge, we present Multimodal Integration with Continual Learning (MIRACLE), a novel online learning framework for the adaptive and efficient integration of single-cell multimodal data. MIRACLE employs dynamic architectures and data rehearsal strategies to support continual learning, allowing diverse data to be integrated while minimizing information loss over time. Our evaluations demonstrate that MIRACLE achieves accurate online integration with reduced computational requirements, effectively updating and expanding atlases with new cross-tissue and cross-modal data, and precisely identifying novel cell types and transferring labels across datasets. MIRACLE provides an efficient and flexible tool for single-cell community to integrate, share and explore biological knowledge from single-cell multimodal data.

## 1 Introduction

Single-cell sequencing technologies enable the in-depth detection of one or multiple omics modalities of functional molecules within individual cells, extending our exploration of biological mechanisms to a single-cell resolution. Integrating and analyzing single-cell sequencing data from different sources and omics can pool together the biological knowledge from heterogeneous data, thus enhancing our understanding of cellular functions [1–3]. Comprehensive single-cell atlases encompassing diverse cell states and types offer a landscape view and in-depth biological insights into cell heterogeneity, thereby accelerating the exploration of molecular mechanisms in biological processes, life development, disease onset, etc [4, 5].

However, single-cell sequencing data with heterogeneous cells are produced asynchronously and continuously in reality. These data often emphasize different biological issues, thus containing different biological knowledge. Incorporating the increasing data continuously will expand this knowledge and provide better biological insights. Therefore, developing methods capable of continuously updating and enhancing atlases with new data is essential.

To date, most existing integration methods are offline, which require all data to be available simultaneously [6–9]. Consequently, each atlas update necessitates the reintegration and reanalysis of all data, leading to a continuous increase in computational time and resources. To address this issue, several online integration methods [10–13] were developed, enabling atlas updates with only new data and improving efficiency to some extent.

Nevertheless, current online methods have significant limitations. Generalization-based methods [11, 13, 14], for instance, utilize models pretrained on large datasets to quickly map new data onto reference atlases, achieving rapid online integration. However, their performance relies heavily on the diversity and scale of the pretraining data, which makes them only suitable for well-developed fields with diverse, large-scale datasets. Furthermore, they struggle to integrate novel knowledge from new data that includes unknown biological or technical variations, leading to difficulties in data adaptation.

In contrast, fine-tuning-based methods [10, 12, 15] continuously adjust models by learning from new data, enabling adaptive online integration. Yet, these methods focus solely on new data during each integration, resulting in the gradual forgetting of previously learned knowledge. This characteristic of knowledge forgetting makes it challenging for these methods to accumulate knowledge during model updates, thus hindering the formation of comprehensive field knowledge. In other words, knowledge retention is a major challenge for fine-tuning-based methods.

Additionally, existing online methods are all designed for specific modalities and features. However, new data often arise from different sequencing technologies and likely possess varied modalities and features, forming ‘mosaic data’ [16] that renders these methods inapplicable.

In summary, there is an urgent need to develop new methods that address data adaptation and knowledge retention while allowing flexible online integration of mosaic data. To this end, we introduce MIRACLE, an online learning framework that effectively achieves online multimodal integration through continual learning. This approach enables models to incrementally learn from new data while retaining previously acquired knowledge [17, 18]. MIRACLE leverages the strategies including dynamic architecture and data rehearsal to adapt to new information while preventing forgetting. It employs MIDAS [19] as the base model to handle multimodal mosaic data, allowing online integration of trimodal mosaic data. Additionally, MIRACLE can precisely map query data onto a reference atlas via an innovative use of online integration, thereby enhancing label transfer. Systematic evaluation has shown that MIRACLE effectively performs online integration of multimodal data across batches, modalities, and cell types using limited memory, achieving accuracy comparable to offline integration with all data while consuming significantly less memory and time. Furthermore, by leveraging the constructed large-scale atlas, MIRACLE continually integrates new cross-tissue and cross-modal data, effectively enabling online updates and expansions of the atlas. The framework also uses the atlas to accurately transfer cell type labels and detect novel cell types in such new data. Users can continuously update MIRACLE with their own data and share it, fostering efficient collaboration and accelerating collective research efforts.

## 2 Results

### 2.1 The MIRACLE model

MIRACLE is a deep generative model [20, 21] combined with continual learning to incrementally integrate singlecell multimodal mosaic datasets of assays for transposase-accessible chromatin (ATAC), RNA and antibody-derived tags (ADT; Fig. 1a). The input data may be collected at different time points and exhibit variations in biological information, technical noise and modality combinations. Whenever new data arrives, MIRACLE can incorporate it to update the existing integration results, removing batch effects while preserving biological variation.

**Fig. 1:**
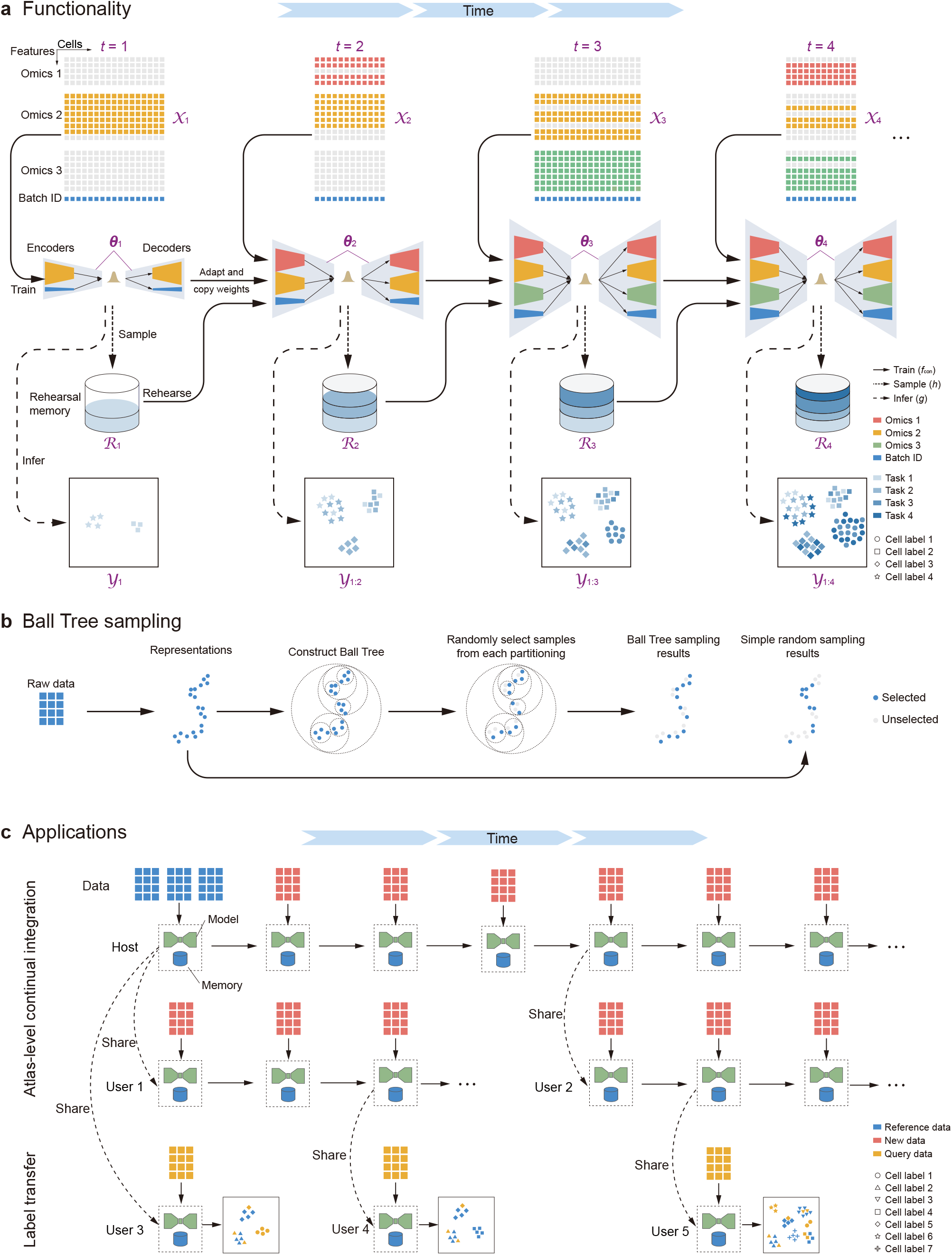
Overview of the MIRACLE framework. **a**, Initially, a model is constructed by training on the data 𝒳_1_ using the function *f*_con_ at time *t* = 1, and the integration results 𝒴_1_ are obtained through the inference function *g*. The rehearsal memory ℛ_1_ is then generated using the sampling function *h*. For subsequent times *t >* 1, new data 𝒳_*t*_ is integrated by updating the previous model ***θ***_*t*−1_ to obtain the new model ***θ***_*t*_ using the function *f*_con_. The integration results 𝒴_1:*t*_ for all data observed up to time *t* are generated by the inference function *g*. Following this, the rehearsal memory is updated from ℛ_*t−* 1_ to ℛ_*t*_. **b**, Based on the Ball Tree partition of representations, the Ball Tree sampling method ensures stability and uniformity compared to simple random sampling. **c**, To achieve atlas-level continual integration, the host can begin by building a model using reference data, and then continuously incorporate new data into this model. Additionally, users can continuously update the model with their own data and re-share it, or use the model to perform label transfer on their data.

MIRACLE uses MIDAS, a multimodal variational autoencoder (VAE) [22], as the base model to better handle mosaic data (Methods). A dynamic architecture approach is used so that the model can adapt to newly added modalities and features in the new data. To mitigate forgetting during the continual integration process, a data rehearsal strategy is employed. We also proposed a novel Ball Tree sampling (BTS) approach to limit the rehearsal memory while preserving the data distribution, enabling memory-efficient training of MIRACLE (Fig. 1b).

With MIRACLE, one can construct large-scale single-cell multimodal atlases and update the atlases by continuously integrating newly added mosaic data (Fig. 1c). The model updated at any time point can be shared with other users. These users can then use their own data to continuously update the model and re-share it, or use the model to transfer labels to their data. These unique features of MIRACLE enable efficient knowledge reuse in practical applications.

### 2.2 MIRACLE enables continual integration using limited memory

MIRACLE employs a rehearsal-based strategy for continual integration. To ensure computational efficiency, we also limit the memory capacity during integration. That is, we perform downsampling once the data exceeds the capacity. Here, we propose the distribution-preserving reservoir sampling (DPRS) approach (Methods), which can dynamically limit the memory capacity while effectively preserve the intrinsic distribution of the data, ultimately obtaining unbiased continual integration results. In this experiment, we collected a large-scale snRNA-seq dataset for human dilated and hypertrophic cardiomyopathy (DHCM) [23], comprising 42 batches and a total of 523,369 cells.

We initially assessed the distribution preservation capability of the proposed BTS algorithm, which is a core component of our DPRS approach. Since DPRS conducts sampling in the latent space, we used MIRACLE to embed the dataset into low-dimensional latent representations and randomly selected 10,000 of them as the overall samples. We then compared the performance of BTS with the classical simple random sampling (SRS) and the latest algorithms, Sketch [24] and scSampler [25], across different sampling ratios. The experiment was repeated 50 times. We used the Maximum Mean Discrepancy (MMD) to quantify the dissimilarity between the original and sampled data distributions. A lower MMD indicates better distribution preservation. The results showed that both BTS and SRS achieved much lower MMDs compared to Sketch and scSampler, with BTS having the lowest MMDs (Fig. 2a). Further comparisons demonstrated that BTS consistently exhibited smaller mean and variance of MMD than SRS across different sampling ratios (Fig. 2b), showcasing its accuracy and robustness in preserving data distribution. We also used UMAP to visualize the maximum MMD results at the lowest sampling ratio of 1/32 (Fig. 2c). The results showed that BTS still effectively preserved the data distribution and outperformed the other three methods. Additionally, we evaluated the sampling time (Fig. 2d). The time taken by BTS was comparable to that of SRS, with both methods being fast and robust to sampling ratios. Conversely, scSampler and Sketch required more time, and the sampling time of scSampler increased dramatically as the sampling ratio increased.

**Fig. 2:**
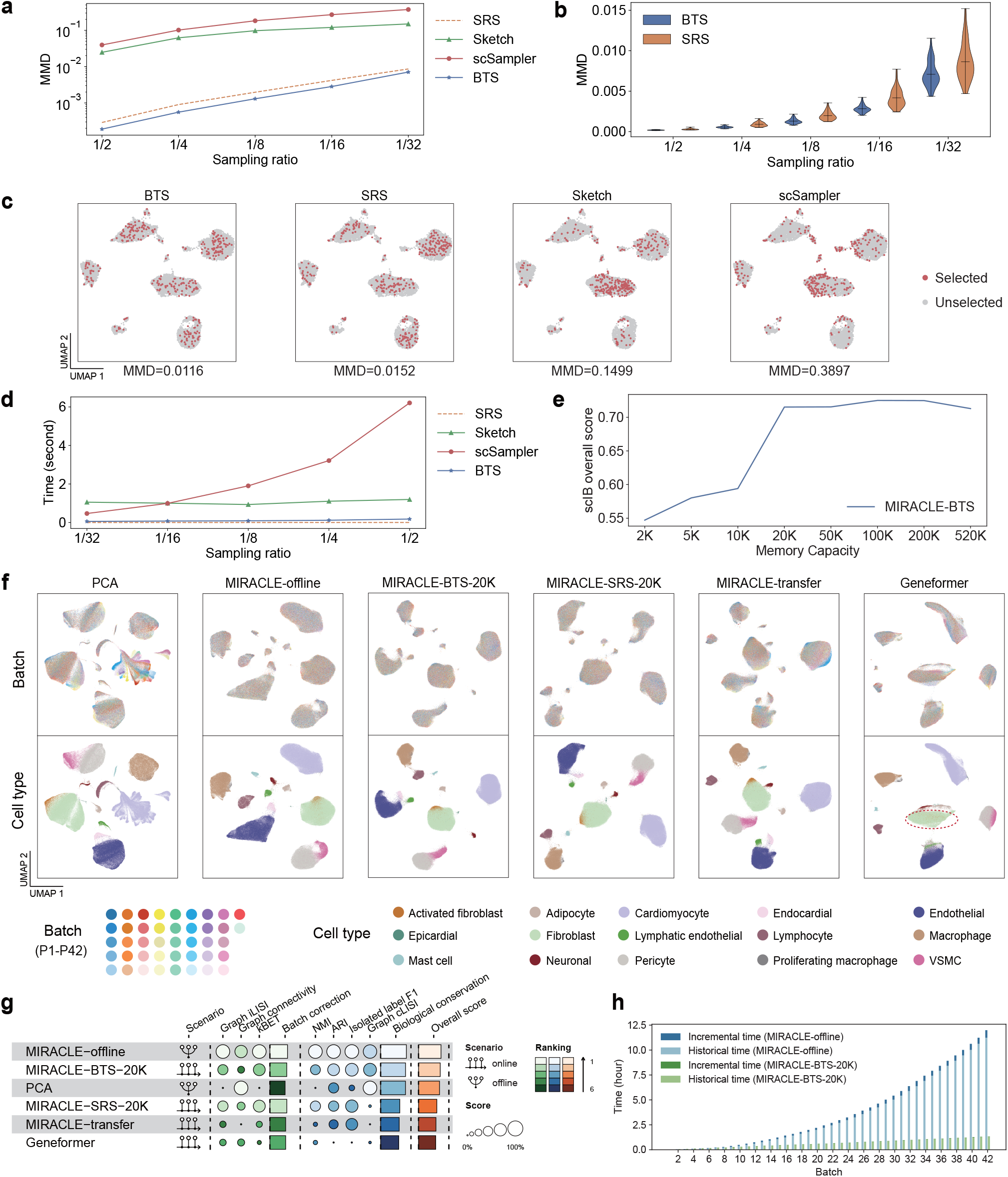
Evaluation of MIRACLE on the DHCM snRNA-seq dataset in memory-constrained continual integration tasks. **a**, Mean MMD for SRS, Sketch, scSampler and BTS at various sampling ratios. **b**, Violin plots comparing MMD values obtained with BTS and SRS across different sampling ratios. **c**, UMAP visualization of the maximum MMD results for each sampling method at a sampling ratio of 1/32. **d**, Runtime comparison for each sampling method across sampling ratios ranging from 1/32 to 1/2. **e**, scIB overall score of MIRACLE-BTS under different memory capacity constraints. **f**, UMAP visualization comparing the biological states inferred by MIRACLE-BTS-20K and other methods. In the Geneformer results, the region containing Fibroblast and Activated Fibroblast cells is highlighted by a red circle. **g**, scIB benchmarking comparing MIRACLE under various settings with other online and offline methods. **h**, Runtime comparison between MIRACLE-offline and MIRACLE-BTS-20K.

Having demonstrated the superiority of BTS, we then explored how different memory capacities impact the continual integration performance of the BTS-based MIRACLE (MIRACLE-BTS; Fig. 2e and Supplementary Fig. 1). The integration performance was quantitatively evaluated using single-cell integration benchmarking (scIB [26]; Methods). It was observed that MIRACLE’s performance improved as the memory capacity increased and reached a saturation point at 20K memory, which accounted for only 1/26 of the total sample size.

We additionally compared the integration performance of the above MIRACLE-BTS with 20K memory (MIRACLE-BTS-20K) against SRS-based MIRACLE with 20K memory (MIRACLE-SRS-20K), offline MIRACLE (MIRACLE-offline, which is trained on all data simultaneously and represents the upper-limit performance for MIRACLE), offline Principal Component Analysis (PCA), transfer-learning-based MIRACLE (MIRACLE-transfer, where no memory rehearsal is used) and the pretrained Geneformer [27]. The UMAP visualization of the integrated low-dimensional representations showed that MIRACLE-offline, MIRACLE-BTS-20K and MIRACLE-SRS-20K effectively removed batch effects and preserved biological differences (Fig. 2f). However, PCA and MIRACLE-transfer struggled to effectively mix different batches, while Geneformer failed to distinguish cell types such as Fibroblast and Activated fibroblast. The scIB results showed that the continual integration strategy MIRACLE-BTS-20K performed comparably to the optimal offline integration strategy MIRACLE-offline and outperformed offline PCA (Fig. 2g and Supplementary Table 1). However, when using SRS (MIRACLE-SRS-20K), not employing memory rehearsal (MIRACLE-transfer), or directly using a pretrained model (Geneformer), the integration performance significantly deteriorated.

To further validate the time efficiency of MIRACLE during the process of continually integrating new data, we compared the time consumption of MIRACLE-BTS-20K and MIRACLE-offline (Fig. 2h). It is worth noting that MIRACLE-offline needs to re-integrate all the batches whenever a new batch arrived. The results demonstrated that as the number of batches increased, the time taken for each integration by MIRACLE-offline also increased, leading to a dramatic rise in cumulative time. In contrast, the time taken for each integration by MIRACLE-BTS-20K remained unaffected by the number of batches, and its cumulative time was significantly lower than that of MIRACLE-offline, demonstrating its practicality.

### 2.3 MIRACLE enables continual integration across batches and cell types

The emerging multimodal single-cell datasets frequently arise from diverse platforms or subjects, necessitating the removal of batch effects and the preservation of biological variations, including cell type composition, during the continual integration. Therefore, in this experiment, we selected a bimodal dataset, WNN, derived from eight distinct batches of the published single-cell CITE-seq dataset of human peripheral blood mononuclear cells (PBMCs) [28]. Additionally, we removed specific cell types based on ground truth labels, ensuring that each of the eight batches possessed unique combinations of cell types (Supplementary Table 2).

We first continually integrated the WNN dataset with MIRACLE. The UMAP visualization of the low-dimensional cell embeddings showed that, MIRACLE robustly integrated new batches into the existing batches and updated all integration results successively, effectively removing batch effects while preserving differences in cell type composition across different batches (Fig. 3a and Supplementary Fig. 2a).

**Fig. 3:**
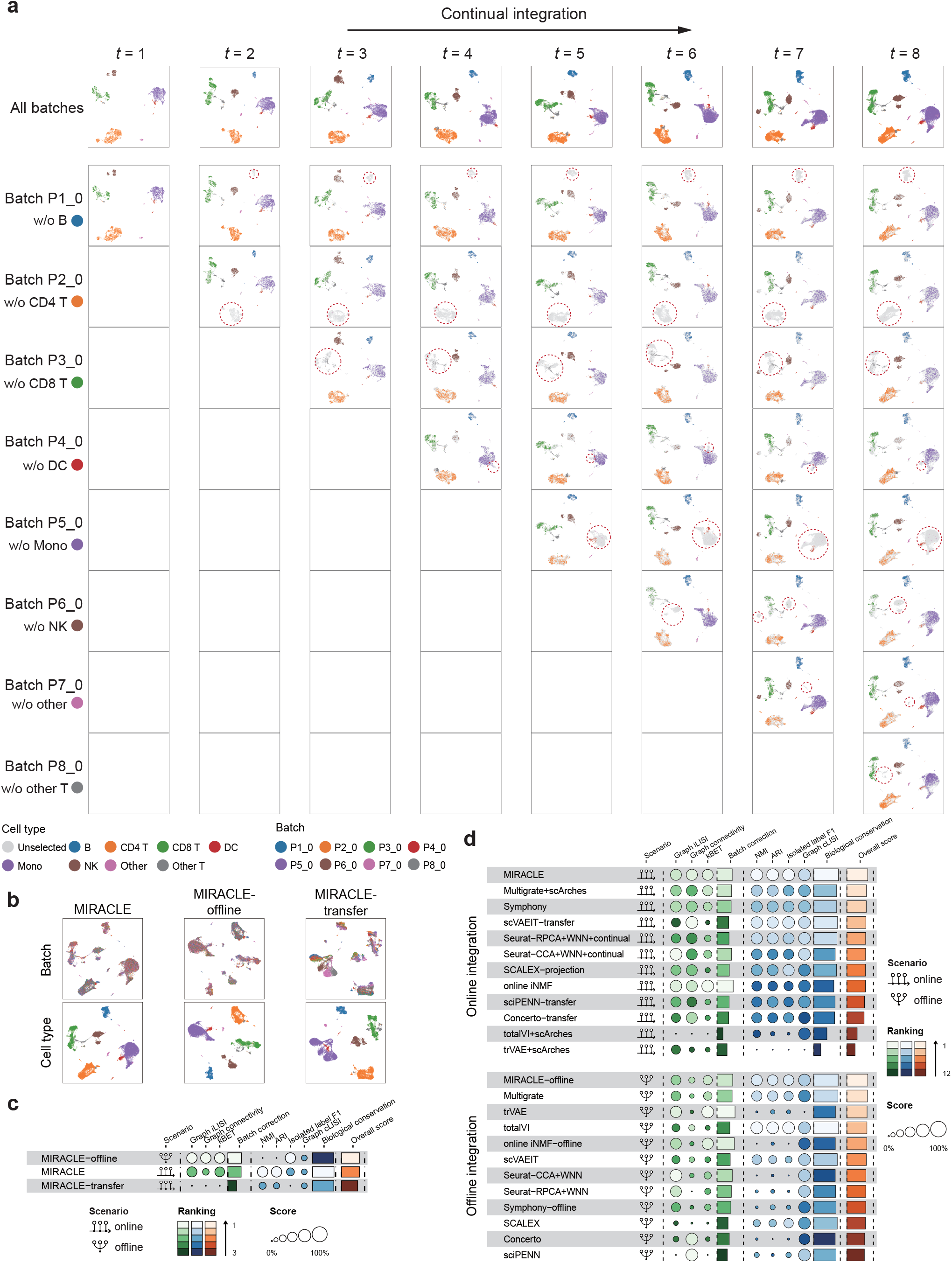
Evaluation of MIRACLE on the WNN bimodal dataset for continual integration across batches and cell types. **a**, UMAP visualization of the biological states inferred by MIRACLE at different time points, with each column corresponding to a specific time point. The first row shows the integration of all available data over time, with the dataset size increasing at each time point. Each row from the 2nd to the 9th displays the integration results for each batch at different time points, with missing cell type regions highlighted by red circles. Cells are colored by cell type. **b**, UMAP visualization of the biological states inferred by MIRACLE and its two variants. **c**, scIB benchmarking comparing MIRACLE with its two variants. **d**, scIB benchmarking comparing MIRACLE against 11 state-of-the-art single-cell integration methods, in both online (upper panel) and offline (lower panel) scenarios.

To validate the effectiveness of our memory rehearsal strategy, we compared MIRACLE with MIRACLE-offline (without knowledge forgetting, serving as the gold standard) and MIRACLE-transfer (without memory rehearsal, serving as the negative control). The UMAP results showed that both MIRACLE and MIRACLE-offline well-mixed the different batches and retained cell type information (Fig. 3b). However, MIRACLE-transfer failed to effectively mix different batches, as evidenced by the formation of multiple internal clusters within the distribution of Mono cells. In line with these findings, the scIB results illustrated that MIRACLE exhibited similar overall performance to MIRACLE-offline and surpassed MIRACLE-transfer, with the latter showing limited performance in batch correction (Fig. 3c). These results demonstrated the efficacy of our memory rehearsal strategy tackling the challenge of catastrophic forgetting.

Then, we compared MIRACLE against 11 state-of-the-art single-cell integration methods on the WNN bimodal dataset in both online and offline settings (Methods). For the methods lacking online integration capabilities, we employed fine-tuning based transfer learning to facilitate the comparison. The UMAP visualizations showed that, under both settings, MIRACLE consistently performed well in mixing different batches and clustering the same cell types (Supplementary Fig. 2b). While some methods showed promise in offline settings, they suffered from pronounced performance degradation in online settings with catastrophic forgetting. For instance, totalVI and trVAE removed batch effects well, but their online versions, totalVI-scArches and trVAE-scArches, failed to mix different batches. Furthermore, scIB results showed that MIRACLE and MIRACLE-offline achieved the highest overall scores in online and offline scenarios, respectively (Fig. 3d and Supplementary Tables 3 and 4). In comparison, while totalVI and trVAE ranked highly in offline settings, their online counterparts lagged behind, highlighting the significance of effective online integration strategies. These results collectively demonstrated MIRACLE’s superiority in continually integrating multi-omics datasets across different batches and cell types.

### 2.4 MIRACLE enables continual mosaic integration

To evaluate MIRACLE’s capability to continually integrate datasets with various modality combinations, we constructed the DOTEA dataset, which originates from two trimodal single-cell human PBMC datasets (DOGMA-seq [29] and TEA-seq [30]) concurrently profiling RNA, ADT and ATAC within individual cells. We selected four batches (LLL_Ctrl, LLL_Stim, DIG_Ctrl and DIG_Stim) from DOGMA-seq and four batches (W3, W4, W5 and W6) from TEA-seq and removed certain omics and cell types from these batches to diversify the dataset. Consequently, DOTEA encompasses four distinct omics combinations of ATAC+RNA, ATAC+ADT, RNA+ADT and ATAC+RNA+ADT, resulting in a mosaic dataset of eight batches spanning different modalities and cell types (Supplementary Table 5).

We first visualized the biological states of different batches and the technical noises inferred by MIRACLE at different integration stages (Supplementary Fig. 3a,b). UMAP results indicated that as the number of integrated batches increased, not only did the biological states of the newly integrated batch align well, but the information learned from prior batches was also retained without catastrophic forgetting, even though some batches had missing omics and cell types. After integrating all the batches, the distributions of both biological states and technical noises across different modalities were remarkably similar (Supplementary Fig. 4a,b), illustrating that MIRACLE is capable of aligning different modalities effectively.

Next, we benchmarked MIRACLE against leading mosaic integration methods in both online and offline scenarios. UMAP results revealed that MIRACLE’s integration performance was on par with that of MIRACLE-offline and far superior to MIRACLE-transfer, effectively preserving biological information and removing batch effects (Fig. 4a and Supplementary Fig. 5). For Multigrate+scArches and scVAEIT-transfer, as the number of batches increased, they were unable to effectively remove batch effects or retain cell type information, even with their offline counterparts. We further conducted a quantitative evaluation using single-cell mosaic integration benchmarking (scMIB [19]; Methods), which is specifically designed for mosaic integration. The results showed that MIRACLE achieved the highest overall score, considerably surpassing scVAEIT, Multigrate and especially their online versions (Fig. 4b and Supplementary Table 6). Remarkably, MIRACLE even outperformed MIRACLE-offline, both of which had significantly higher batch correction scores than the others. These findings demonstrated that MIRACLE could continually integrate datasets across different batches, modalities and cell types.

**Fig. 4:**
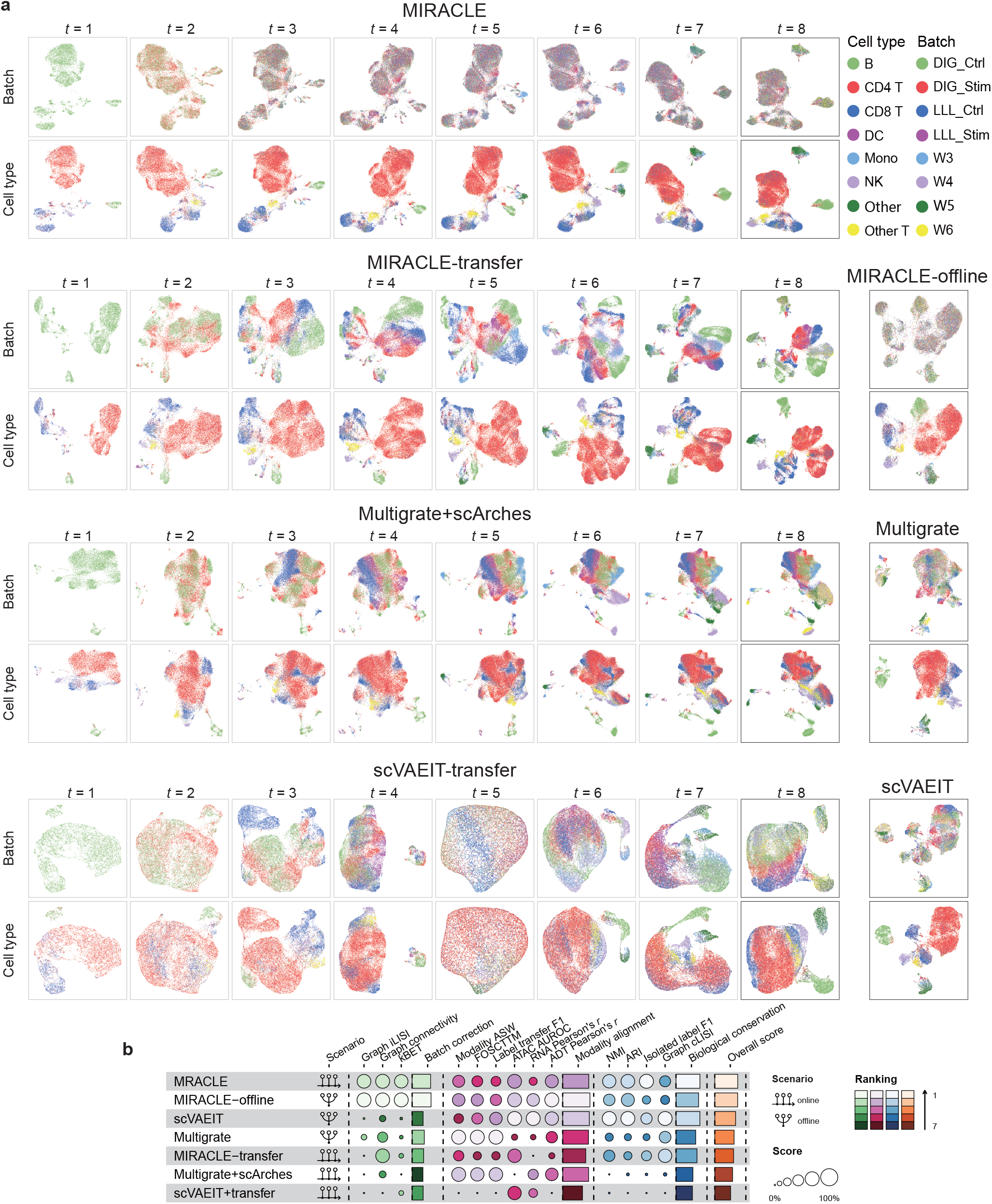
Evaluation of MIRACLE on the DOTEA trimodal dataset in continual mosaic integration tasks. **a**, For online methods, UMAP visualizations show the biological states of all cells observed up to each time point. For offline methods, UMAP visualizations of all cells are displayed. Cells are colored by batch and cell type. **b**, scMIB benchmarking comparing MIRACLE with Multigrate and scVAEIT in both online and offline settings.

### 2.5 Continual construction of cross-tissue multimodal atlas

Since single-cell multimodal atlases are often constructed from large-scale datasets with high-dimensional features, updating the atlas by reintegrating all the datasets every time new data is added requires significant computational time and resources, thus necessitating online integration. Here, we evaluate MIRACLE in atlas-level continual integration under both within- and cross-tissue scenarios. For within-tissue scenario, we collected 10 human PBMC datasets comprising 34 batches, where 28 batches were taken as the reference data for initial atlas construction and 6 batches including NEAT (ATAC+RNA) [31], ASAP (ATAC+ADT) [29] and ASAP-CITE (RNA+ADT) [29] were taken as the new data for continual atlas updating. We denote the first 28 batches as atlas-no-nac and the subsequent 6 batches as nac, respectively. For cross-tissue scenario that is crucial for understanding tissue-wise cell behavior, we further collected 3 batches related to human immunity, including data from the tonsil (RNA) [32], bone marrow (ATAC+ADT) [29] and spleen (RNA) [33], which are used to update the reference atlas constructed from the 34 PBMC batches (see Supplementary Table 7 for details).

In each scenario, we first used MIRACLE to integrate the reference batches offline, and then integrated the new batches online based on the pretrained model. Here, we compared three online integration strategies, including the default MIRACLE based on continual learning, MIRACLE-generalization which directly uses the pretrained model to infer latent variables [11, 13] and MIRACLE-transfer which fine-tunes the model on the new batches [12, 15]. Additionally, to evaluate the performance of online integration, we also used MIRACLE-offline to integrate the reference batches and new batches offline to obtain a gold standard for subsequent quantitative evaluation experiments (Supplementary Fig. 6a,b).

The UMAP visualizations demonstrated that MIRACLE effectively maintained cell type information while removing batch effects (Fig. 5a,b). Remarkably, new cell types in new cross-tissue datasets (Germinal center B and Cycling B cells in the tonsil and Progenitor cells in the bone marrow) formed distinct clusters (Fig. 5b). Additionally, MIRACLE achieved comparable performance to MIRACLE-offline, accomplishing the retention of previously learned knowledge (Supplementary Figs. 7 and 8). MIRACLE-generalization managed to preserve cell type information of the new PBMC batches to some extent but was unable to identify novel cells in new cross-tissue batches (Supplementary Fig. 9a,b), while MIRACLE-transfer failed to distinguish cell types in the new batches or to eliminate batch effects in both within- and cross-tissue scenarios (Supplementary Fig. 9c,d).

**Fig. 5:**
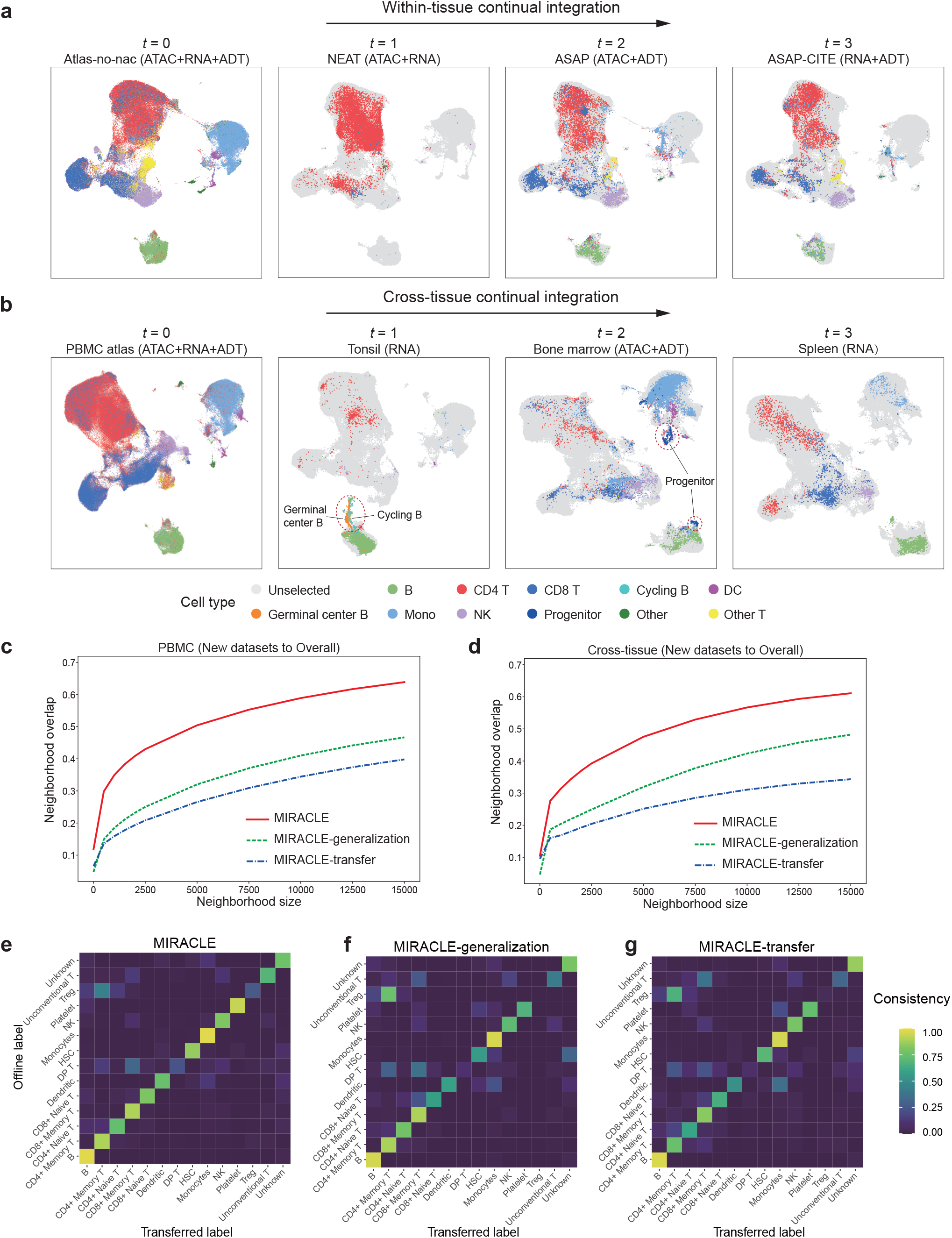
Evaluation of MIRACLE in continual multimodal atlas construction tasks. **a**,**b**, UMAP visualization of the biological states inferred by MIRACLE, showing the process of within-tissue (**a**) and cross-tissue (**b**) continual atlas construction. Novel cell types are highlighted by red circles. **c**,**d**, Neighborhood overlap analysis between the dimensionality reduction results of online strategies and MIRACLE-offline across different neighborhood sizes in within-tissue (**c**) and cross-tissue (**d**) continual integration tasks. **e-g**, Label confusion plots comparing transfer consistency for MIRACLE (**e**), MIRACLE-generalization (**f**), and MIRACLE-transfer (**g**) in the cross-tissue continual integration task.

Next, to quantitatively evaluate the cell neighborhood consistency between online and offline integration results, for each online integration strategy, we calculated the neighborhood overlap between its dimensionality reduction results and those of MIRACLE-offline. The results indicated that both MIRACLE and MIRACLE-generalization exhibited significantly higher neighborhood overlap compared to MIRACLE-transfer (Supplementary Fig. 10a,b). However, when examining the neighborhoods of the new data, the overlap achieved by MIRACLE was substantially higher than those of the other two strategies, indicating that MIRACLE is capable of integrating new data more appropriately (Fig. 5c,d). We also evaluated the cell type consistency between online and offline integration results through label transfer. Confusion plots revealed that the labels transferred by MIRACLE closely matched those of MIRACLE-offline and were significantly more accurate than those of the other two strategies (Fig. 5e-g and Supplementary Fig. 10c-e). These findings demonstrated that MIRACLE can effectively support the continual construction of cross-tissue single-cell multimodal atlases and yield results consistent with offline construction, approaching the gold standard.

### 2.6 Label transfer for cross-tissue mosaic data

In addition to continuous atlas construction, MIRACLE offers label transfer functionality. Once the reference atlas is constructed, labels can be transferred from the atlas to the query set, enabling rapid analysis of new query data. By innovatively applying continual integration, MIRACLE achieves the matching between the query set and the reference set, supporting subsequent label transfer. To prevent cross-contamination between different query sets, for both within- and cross-tissue scenarios, we conducted independent label transfers for each query set using the initial reference atlas. We also compared MIRACLE with its two variants, MIRACLE-generalization and MIRACLE-transfer, which use traditional matching strategies based on generalization [11, 13, 14, 34, 35] and fine-tuning-based transfer learning [12, 15], respectively. For each method, once the reference and query sets were matched, we used a kNN algorithm based on novelty detection to transfer labels. This kNN algorithm can identify new cell types in the query set and label them as ‘Unknown’.

UMAP results showed that the labels transferred from the reference atlas to the query set by MIRACLE exhibited high consistency with the query set’s ground truth labels (Fig. 6a and Supplementary Fig. 11). Furthermore, in the cross-tissue scenario, the cells identified as ‘Unknown’ corresponded precisely to the new cell types within the query set (Cycling B and Germinal center B cells in the tonsil, and Progenitor cells in the bone marrow; Fig. 6a). However, the mapping results of MIRACLE-generalization and MIRACLE-transfer displayed significant batch effects (Supplementary Figs. 12 and 13), and the transferred labels were inconsistent with the ground truth labels (Supplementary Figs. 12a and 13). Moreover, these two methods struggled to recognize new cell types in cross-tissue query sets (Supplementary Fig. 13a,b).

**Fig. 6:**
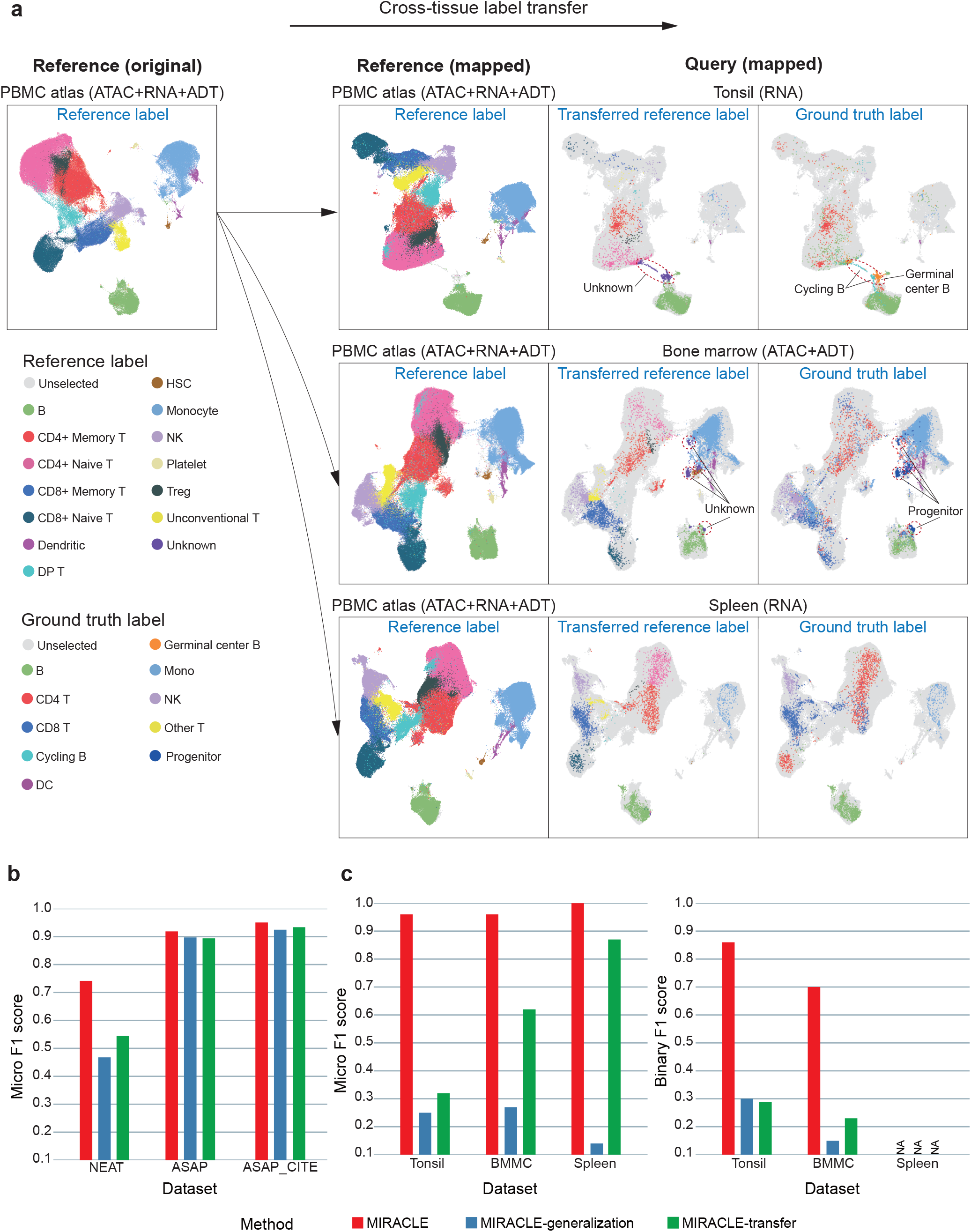
Evaluation of MIRACLE in label transfer tasks. **a**, UMAP visualization of the biological states inferred by MIRACLE for cross-tissue label transfer. Column 1 shows the original PBMC reference, while Column 2 shows the mapped PBMC references after separately incorporating different cross-tissue queries, both colored by reference label. Columns 3 and 4 show the three mapped cross-tissue queries, colored by transferred reference label (Column 3) and ground truth label (Column 4). **b**, Micro F1 scores for MIRACLE and two other strategies in within-tissue label transfer tasks. **c**, Micro F1 scores (left panel) and binary F1 scores (right panel) for MIRACLE and two other strategies in cross-tissue label transfer tasks.

Quantitative results showed that, in the within-tissue scenario, MIRACLE’s label transfer consistency (Supplementary Fig. 14) and micro F1 scores (Fig. 6b) both surpassed the other two strategies. This was particularly notable in the NEAT query set, which included only CD4 T cells and was harder to match with the reference set containing multiple cell types. In the cross-tissue scenario, MIRACLE’s label transfer consistency (Supplementary Fig. 15), as well as its micro and binary F1 scores for new cell type identification (Fig. 6c), were significantly higher than those of the other strategies. These findings suggested that MIRACLE could achieve accurate and robust label transfer and new cell type identification across batches, modalities, and tissues.

## 3 Discussion

The rapid generation of single-cell sequencing data from laboratories worldwide, often with various modalities from various platforms, presents an ongoing challenge for data integration. Different datasets often focus on distinct biological questions and introduce heterogeneous modalities, necessitating continual integration to construct an incrementally comprehensive single-cell atlas that can encapsulate progressively expanding biological knowledge. To address this pressing need, we propose MIRACLE—a framework designed to continuously integrate new data across varying modalities, batches, and cell types, while preserving previously acquired knowledge. By combining continual learning strategies with multimodal deep generative models, MIRACLE achieves efficient and precise online multimodal integration, delivering performance close to its offline counterpart, which represents its upper limit, while significantly reducing memory and time consumption. Our evaluations demonstrate that MIRACLE surpasses existing online methods, especially in its ability to assimilate novel information while preserving prior knowledge. For example, based on a pre-constructed large-scale PBMC trimodal atlas, MIRACLE effectively updates the atlas by integrating data from different tissues (e.g., tonsil, bone marrow, spleen), platforms, and modalities. Moreover, MIRACLE facilitates precise cross-tissue label transfer and identifies new cell types in the new data.

The importance of continual integration is underscored by the ongoing accumulation of new single-cell data, which provides an invaluable opportunity to expand and refine biological knowledge. Traditional methods, however, often require computationally expensive retraining on the entire dataset for each update, which becomes especially prohibitive for smaller laboratories with limited resources. MIRACLE addresses this challenge by offering a memory-efficient and computationally lightweight solution that minimizes both memory usage and computational time. This makes continual integration feasible even for resource-constrained laboratories. MIRACLE provides an opportunity for a broader range of research groups to integrate, share, and benefit from a continuously updated body of knowledge, hence promoting more efficient and comprehensive understanding of biological mechanisms.

In this work, MIRACLE leverages the MIDAS model to effectively integrate mosaic data, demonstrating impressive robustness and adaptability. Notably, MIRACLE’s continual learning strategy is versatile and can be seamlessly incorporated into most existing deep learning-based integration frameworks [7, 9, 12, 13, 15, 35], enabling their application in dynamic, online scenarios. Looking ahead, a promising direction involves exploring more advanced dynamic architecture strategies [36, 37], which could enable the model to automatically optimize its structure based on the complexity of the incoming data, ensuring scalability and adaptability across a wide range of applications. Another potential avenue is to facilitate more collaborative model updates across distributed environments. By incorporating federated learning techniques [38, 39], MIRACLE could enhance knowledge sharing among multiple users while simultaneously preserving data privacy.

In conclusion, MIRACLE not only addresses the critical challenges of data adaptation and knowledge retention in the continuously evolving field of single-cell multimodal sequencing, but also lays the foundation for future advancements in the seamless integration and sharing of biological knowledge.

## 4 Methods

### 4.1 Single-cell continual integration with continual learning

#### The scenario of single-cell continual integration

Single-cell continual integration aims to continuously incorporate new single-cell data into the atlas, ensuring up-to-date knowledge. Let 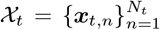 be the new data batch of *N*_*t*_ cells obtained at the current time *t*, where the vector ***x***_*t,n*_ is the observation of the *n*-th cell. Additionally, let 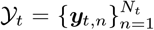 be the corresponding integration results of 𝒳_*t*_, where the vector ***y***_*t,n*_ can include the low-dimensional representation or batch-corrected data of cell *n*. The goal of continual integration is to update all integration results 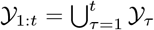 whenever new data 𝒳_*t*_ is available.

#### Offline strategy

To achieve continual integration, the common approach is to use an offline strategy [6–9]:

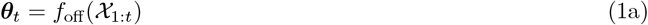

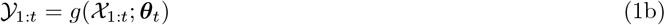

Upon obtaining new data 𝒳_*t*_, all data 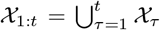 are used to train the model parameters ***θ***_*t*_ via the offline learning function *f*_off_. The inference function *g* then uses ***θ***_*t*_ to predict the integration results 𝒴 _1:*t*_ for 𝒳_1:*t*_ based on each cell. While effective, this strategy requires retraining on all data 𝒳_1:*t*_ each time, leading to decreasing efficiency.

#### Online strategy 1: generalization

To avoid repeated retraining during continual integration, online strategies can be adopted. One efficient method is generalization [11, 13, 14]:

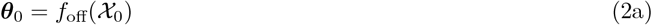

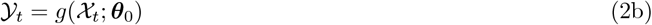

Here, the model is first pre-trained offline on large external datasets 𝒳_0_ through *f*_off_ to obtain parameters ***θ***_0_. Then, ***θ***_0_ is fixed and generalized to new data 𝒳_*t*_, that is, inferring 𝒴_*t*_ for each 𝒳_*t*_ using *g*. Since ***θ***_0_ is fixed, there is no need to update past results 𝒴_1:*t*− 1_. Although this strategy avoids training during continual integration, it relies on external datasets, which often fail to capture unknown biological and technical variations in new data, resulting in poor adaptability.

#### Online strategy 2: fine-tuning

To continuously adapt to unknown changes in new data, online integration strategies based on fine-tuning have emerged [10, 12, 15]:

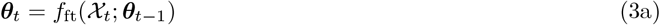

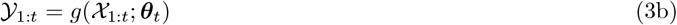

In the *t*-th integration, the fine-tuning function *f*_ft_ fine-tunes the previous model parameters ***θ***_*t* 1_ on new data 𝒳_*t*_ to obtain updated parameters ***θ***_*t*_, which are then used to predict 𝒴_1:*t*_ through *g*. Note that ***θ***_0_ can be obtained via pre-training or random initialization. However, as the number of integrations *t* increases, ***θ***_*t*_ will gradually forget past information during updates, affecting the integration results, especially those of the past data.

#### Online strategy 3 (ours): continual learning

To enable the model to adapt to new data while preventing forgetting, we propose a novel online integration strategy based on continual learning [17, 18]:

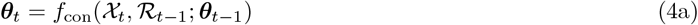

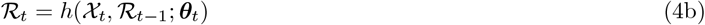

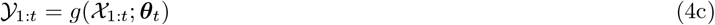

Unlike traditional fine-tuning, continual learning prevents forgetting during continual integration by introducing rehearsal memory ℛ_*t*_ ⊆ 𝒳_1:*t*_ to retain key data. By combining new data 𝒳_*t*_ with previous memory ℛ_*t*−1_, the continual learning function *f*_con_ updates the parameters from ***θ***_*t*−1_ to ***θ***_*t*_. The updated memory ℛ_*t*_ is then obtained through the sampling function *h*, and predictions 𝒴_1:*t*_ are made using the inference function *g*. Initially, the memory ℛ_0_ is set to Ø, and the parameters ***θ***_0_ can be either pre-trained or randomly initialized.

### 4.2 Dynamic architecture

In continual integration, new data 𝒳_*t*_ often originates from various sequencing technologies and may introduce new modalities or features. To seamlessly incorporate these, we propose dynamically expanding the model architecture in the parameter update equation (Eq. (4a)). Let 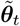 be the model parameters associated with the new modalities or features. We randomly initialize 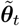 and combine it with the previous parameters ***θ***_*t*−1_ to get the initial value 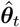:

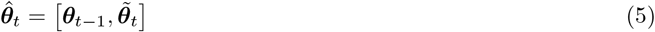

Based on 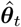, we can then obtain ***θ***_*t*_ by training on 𝒳_*t*_ and ℛ_*t*−1_.

### 4.3 Distribution-preserving reservoir sampling

In the memory update equation Eq. (4b), the capacity of rehearsal memory ℛ_*t*_ increases with more integration iterations, affecting computational efficiency and knowledge sharing. To address this, we propose a batched reservoir sampling algorithm that limits the capacity of ℛ_*t*_ when updating with a new batch 𝒳_*t*_. Additionally, to achieve unbiased integration results, we combine stratified sampling to preserve inter-batch distribution and introduce a new Ball Tree sampling method to maintain intra-batch distribution. This combined sampling method is called DPRS.

#### Handling batch data via batched reservoir sampling

During the continual addition of new batches, to ensure each cell’s data has the same probability of being retained in the rehearsal memory with limited capacity, the reservoir sampling algorithm can be adopted. However, the classical reservoir sampling algorithm only handles single new cells. To efficiently handle new batches with multiple cells, we propose batched reservoir sampling.

Assume the rehearsal memory has a capacity limit of *M* cells. After each integration, we add new data 𝒳_*t*_ of size *N*_*t*_ to the previous rehearsal memory ℛ_*t*−1_ of size *M*_*t*−1_, resulting in ℛ_*t*_. However, if *N*_*t*_ + *M*_*t*−1_ *> M*, we first sample ℛ_*t*−1_ and 𝒳_*t*_ with sampling ratios *α* and *β*, respectively, where *α* and *β* satisfy:

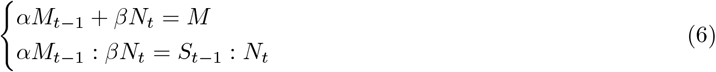

where *S*_*τ*_ = *N*_1_ +*N*_2_ + · · · +*N*_*τ*_ is the total number of cells in the first *τ* batches. Eq. (6) ensures ℛ_*t*_ has size *M*, with new and old data proportions consistent with their original batches. From Eq. (6), we get *α* = *S*_*t* − 1_*M/*(*S*_*t*_*M*_*t*− 1_) and *β* = *M/S*_*t*_. Note that when *N*_*t*_ = 1, batched reservoir sampling reduces to classical reservoir sampling.

#### Preserving inter-batch distribution via stratified sampling

To maintain cell distribution across batches, in addition to batched reservoir sampling, we further introduce batch-based stratified sampling on past data. Specifically, in memory ℛ_*t*−1_, assuming 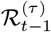 is the data from the *τ* -th batch and is of size 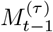, we sample each 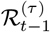 with a ratio of *α* for *τ* = 1, 2, …, *t* − 1. This ensures that the cell proportions in ℛ_*t*_ match those in the original batches.

#### Preserving intra-batch distribution via Ball Tree sampling

When sampling data within a batch, a common method is SRS. However, SRS struggles to maintain data distribution, especially at low sampling ratios [40]. To better preserve the data distribution within each batch at various sampling ratios, we propose a novel Ball Tree sampling algorithm.

Assume the data before sampling is generated from the underlying distribution *p*(***x***). We divide the data space 𝒟 into infinitesimal intervals _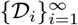_, each with a Lebesgue measure *λ*(𝒟_*i*_) close to 0. Then, for any ***x***, assuming it lies in the *j*(***x***)-th interval, *p*(***x***) can be approximated as:

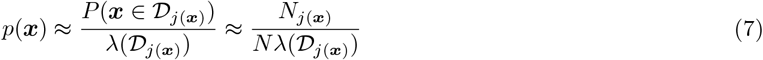

where *N* is the total number of cells to be sampled, and *N*_*j*(***x***)_ is the number of cells in 𝒟_*j*(***x***)_. Similarly, after sampling with ratio *γ*, the underlying distribution 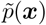 is approximated as:

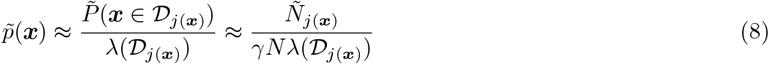

where *γN* is the total number of cells after sampling, and *Ñ*_*j*(***x***)_ is the number of cells in 𝒟_*j*(***x***)_ after sampling. To ensure the sampled distribution matches the original, 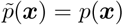 is required. Combining Eq. (7) and Eq. (8), we get *Ñ*_*j*(***x***)_ *γN*_*j*(***x***)_, which means that for any interval 𝒟_*i*_, we should sample at a ratio of *γ*.

Before sampling, we need to partition the space 𝒟. A common method is equal-interval partitioning, where each interval has the same Lebesgue measure *λ*(𝒟 _*i*_). However, this method struggles with non-uniform data, lacking detail in dense regions and missing data in sparse regions. Additionally, the required number of intervals grows exponentially with the space’s dimensionality, leading to the ‘curse of dimensionality’. To address this, we use equal-frequency partitioning, where each interval has the same number of cells *N*_*i*_. This approach automatically adjusts the interval sizes based on data density, handling non-uniform data more effectively. Moreover, the number of intervals only grows linearly with the total sample size *N*, mitigating the impact of data dimensionality.

To improve the accuracy of distribution preservation, we can use fine-grained equal-frequency partitioning, setting the number of samples per interval *N*_*i*_ to 1*/γ*. For example, when *γ* = 1*/*3, we have *N*_*i*_ = 3. Thus, we can divide 𝒟 into intervals with 3 samples each and randomly select 1 sample from each interval to complete sampling. However, because 1*/γ* may not always be an integer, we propose a multi-resolution space partitioning method. By selecting samples from intervals at different resolutions, we can achieve fine-grained sampling at any ratio. Here, we use the Ball Tree method [41] for multi-resolution partitioning, which is better suited for non-uniform and high-dimensional data compared to the traditional KD Tree. We refer to this intra-batch distribution-preserving sampling algorithm based on Ball Tree partitioning as BTS, detailed in Supplementary Note 1.

### 4.4 Handling mosaic data

During the process of continuous integration, different batches of single-cell data may contain different combinations of modalities, which is known as mosaic data. To handle mosaic data, we use MIDAS as the base model. Suppose at time *t*, the set of modalities present in the observed data 𝒳_1:*t*_ is ℳ, for example, ℳ = {ATAC, RNA, ADT}. For the *n*-th cell observation ***x***_*τ,n*_ in batch *τ* ∈ {1, …, *t*}, we denote it as:

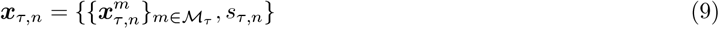

where ℳ_*τ*_ ⊆ ℳ is the set of modalities in batch 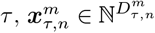 is the observation vector of modality with size 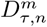, and *s*_*τ,n*_ = *τ* is the batch ID.

Using MIDAS, we obtain the integration result ***y***_*τ,n*_ for each cell observation ***x***_*τ,n*_:

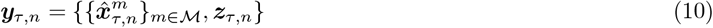

where 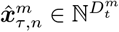, with size 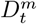, is the imputed and batch-corrected count vector for modality *m*, and 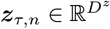 is the low-dimensional cell embedding representing the biological state of the cell.

### 4.5 Training MIRACLE

MIRACLE uses MIDAS as the base model to handle single-cell multimodal data with diverse modality combinations. For the *t*-th integration, the model parameters ***θ***_*t*_ consist of 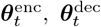 and 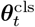 for the encoders, decoders and classifiers, respectively. These parameters are optimized by iteratively minimizing the encoder-decoder loss function 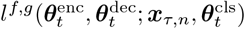 and the classifier loss function 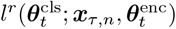 over new data 𝒳_*t*_ and rehersal data ℛ_*t*−1_, where ***x***_*τ,n*_ ∈ 𝒳_*t*_ for *τ* = *t* and 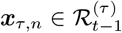 for 1 ≤ *τ < t*. The detailed MIRACLE training process corresponding to Eq. (4a) is illustrated in Algorithm 1.

### 4.6 Integrating multiple new batches at once

In practical applications, new data often arrives in multiple batches, denoted as {𝒳_*t*_, *𝒳*_*t*+1_, … }, which may also consist of mosaic data. In this case, integrating these data batches simultaneously, rather than sequentially, can enhance the efficiency of continuous integration. Leveraging MIDAS’s capability to support mosaic data, MIRACLE can be seamlessly extended to accommodate this scenario.

### 4.7 Datasets

All datasets used in our experiments were publicly available (Supplementary Table 8). For data analysis, we downloaded count matrices for gene unique molecular identifiers (UMIs), ATAC fragments, and ADTs.

#### Algorithm 1

The MIRACLE training algorithm.

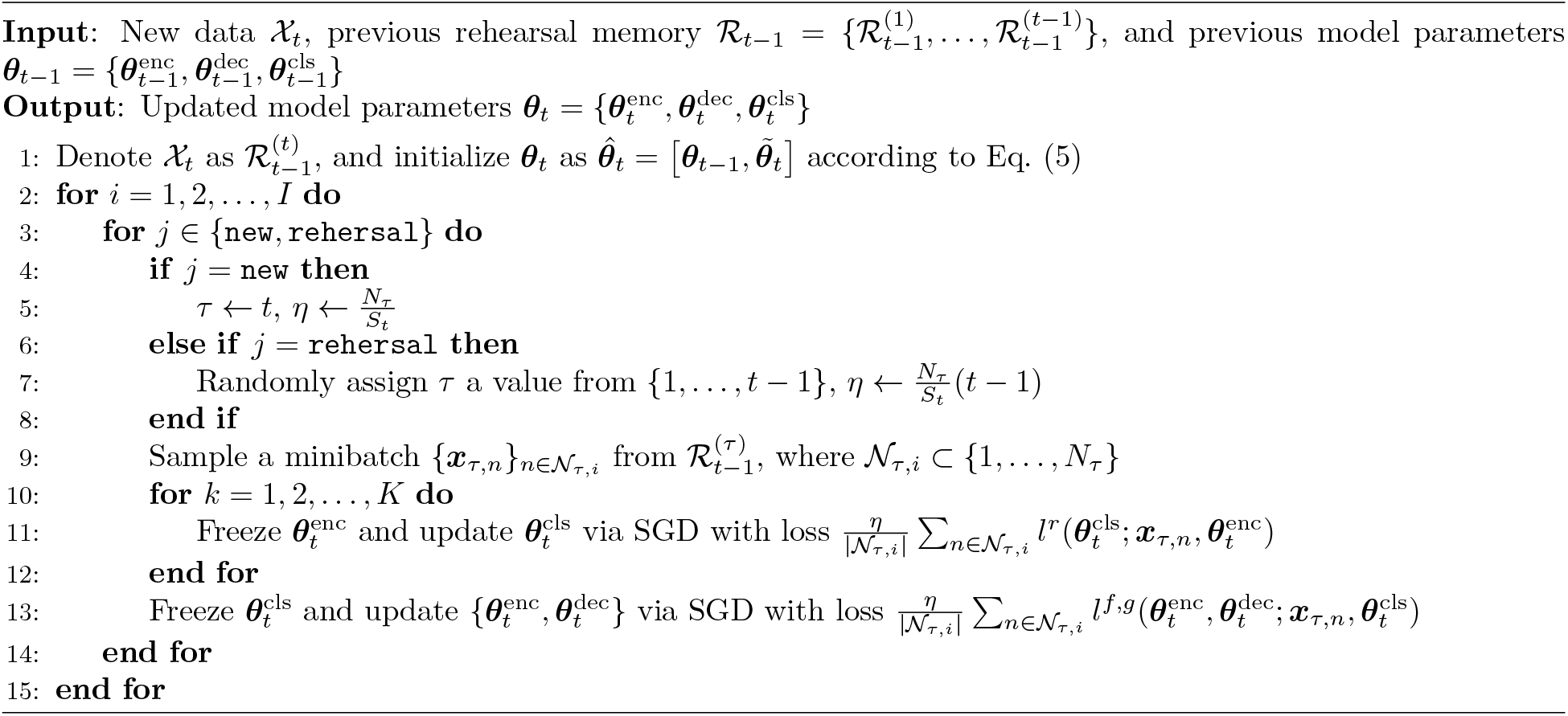

#### DHCM dataset

The DHCM dataset [23] an snRNA-seq dataset spanning 42 batches. It can be downloaded from https://singlecell.broadinstitute.org/single_cell/study/SCP1303/ single-nuclei-profiling-of-human-dilated-and-hypertrophic-cardiomyopathy.

#### WNN dataset

The WNN dataset [28] was profiled by CITE-seq and measures paired RNA and ADT data. It can be acquired from https://atlas.fredhutch.org/nygc/multimodal-pbmc.

#### DOGMA dataset

The DOGMA dataset [42] contains RNA, ATAC, and ADT data across four batches. They were obtained from Gene Expression Omnibus (GEO) under accession ID GSE166188.

#### TEA dataset

The TEA dataset [30] contains RNA, ATAC, and ADT data across four batches. These batches were obtained from GEO under accession ID GSE158013.

#### TEA Multiome dataset

The TEA Multiome dataset [30] is profiled by 10x Chromium Single Cell Multiome ATAC + Gene Expression, which contains ATAC and RNA data across two batches. These batches were obtained from GEO under accession ID GSE158013.

#### 10X Multiome dataset

The 10X Multiome dataset [43–46] contains ATAC and RNA data across four batches. These batches were obtained from 10x Genomics https://www.10xgenomics.com/resources/datasets/.

#### DOGMA-CITE dataset

The DOGMA-CITE dataset [47] consists of one batch generated by CITE-seq measuring RNA and ADT data and three batches generated by DOGMA-seq measuring ATAC, RNA, and ADT data. These batches were obtained from GEO under accession ID GSE200417.

#### ISSAAC dataset

The ISSAAC dataset [30] is generated by ISSAAC-seq, which contains only one batch that measures ATAC and RNA data. The raw sequencing data was obtained from ArrayExpress under accession number E-MTAB-11264.

#### NEAT dataset

The NEAT dataset [31] is generated by NEAT-seq, which contains ATAC and RNA data across two batches. These batches were obtained from GEO under accession ID GSE178707.

#### ASAP dataset

The ASAP dataset [29] is generated by ASAP-seq, which contains ATAC and ADT data across two batches. These batches were obtained from GEO under accession ID GSE156473.

#### ASAP-CITE dataset

The ASAP-CITE dataset [29] is generated by CITE-seq, which contains RNA and ADT data across two batches. These batches were obtained from GEO under accession ID GSE156473.

#### TONSIL dataset

The TONSIL dataset [32] contains only one batch measuring RNA data, which was obtained from GEO under accession ID GSE165860.

#### BMMC dataset

The BMMC dataset [29] contains only one batch measuring ATAC and ADT data, which was obtained from GEO under accession ID GSE156477.

#### SPLEEN dataset

The SPLEEN dataset [33] contains only one batch measuring RNA data, which was obtained from GEO under accession ID GSE159929.

### 4.8 Data preprocessing

When handling single-cell multimodal data that may encompass multiple batches, we begin with quality control and feature selection for each individual batch. Subsequently, we unify the features across different batches. We used the Seurat package (v4.1.0) [28] to process the count matrices of RNA and ADT.

For RNA data, we assessed metrics such as the number of detected genes per cell, total UMI count, and the percentage of mitochondrial RNA reads to filter out low-quality cells. We then utilized the NormalizeData function in Seurat to normalize and log-transform the UMI count data. Next, we removed genes with low frequency and used the FindVariableFeatures function to select the top 4000 highly variable genes for each batch. Finally, we employed the SelectionIntegrationFeatures function to identify their union genes, resulting in a set of 4000 final features.

For ADT data, we evaluated the total protein tag number to exclude low-quality cells. Following this, we applied centered log-ratio normalization using the NormalizeData function in Seurat. In our experiments, we retained all ADT features without performing further feature selection.

To process ATAC fragment files, we used the Signac package (v1.6.0) [48] and preprocessed the peaks via MACS2 [49]. We evaluated the transcription start site enrichment score and nucleosome signal for each batch for quality control. Afterward, we merged the peaks using the reduce function in Signac. Following the peak merging step, we recalculated the fragment counts within the merged peaks. The UMI counts for RNA data, tag counts for ADT data, and binarized fragment counts for ATAC data were used as inputs.

When processing newly arrived single-cell multimodal data, we followed the same procedure. Notably, to prevent excessive expansion of ATAC features during continual integration, we aligned the peaks with those from previous batches.

### 4.9 Generating third-party cell-type labels

To achieve a comprehensive qualitative and quantitative evaluation of PBMC datasets, we used the third-party tool Seurat for cell type annotation via label transfer. The reference dataset selected for this task was the CITE-seq PBMC atlas from [28]. Label transfer was conducted using Seurat’s FindTransferAnchors and TransferData functions, utilizing the ‘cca’ reduction method for reference mapping. In cases where raw RNA expression data was not available, we first used the GeneActivity function in Signac to generate a gene activity matrix based on ATAC data. This gene activity matrix was then employed for label transfer. However, due to the predominance of CD4 T cells in the NEAT dataset, there may be a limitation in finding suitable reference datasets for accurate annotation. Therefore, after performing the initial annotation of the NEAT dataset, we applied Seurat’s FindClusters function to cluster the data and refine the annotation results.

For the TONSIL dataset, we uploaded its gene expression data to Azimuth, a reference-based single-cell analysis tool available at https://azimuth.hubmapconsortium.org, to perform automated cell annotation. We then manually adjusted the cell annotation results by integrating Seurat’s clustering outcomes. For the BMMC dataset, Azimuth’s ATAC application was used to annotate the ATAC data at the cellular level. For the SPLEEN dataset, we employed SingleR [50], a tool for unbiased cell type identification from single-cell RNA sequencing data, to annotate its RNA data, followed by manual refinement of the annotations using Seurat clustering results.

### 4.10 Comparison of sampling methods for distribution preservation

To evaluate the distribution preservation capability of our BTS algorithm, we compared its performance with SRS, Sketch, and scSampler.

#### SRS

We used the subsample function in the Scanpy package [51] to perform SRS. To ensure diverse sampling results, we set different random seeds for each run of the experiment.

#### Sketch

The Sketch method [24] is designed to preserve rare populations during sampling. We utilized the SketchData function in the Seurat package (v5.0.0) for sample selection [52].

#### scSampler

The python package scSampler [25] is also designed to preserve rare populations and can be downloaded from https://github.com/SONGDONGYUAN1994/scsampler. Since our input data were already low-dimensional embeddings, we skipped the default reduction process.

#### Evaluation of distribution preservation

We used the MMD metric to evaluate the consistency of data distributions before and after sampling. A smaller MMD value indicates better sampling performance. Given two sets of samples 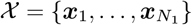 and 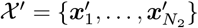, MMD is defined as:

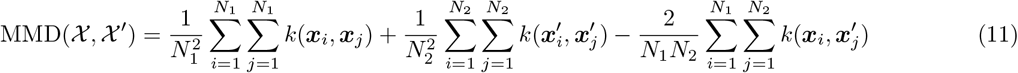

where *k*(·, ·) is the Gaussian kernel function.

### 4.11 Implementation of comparative integration methods

We compared the horizontal, rectangular, and mosaic integration methods (Supplementary Table 9). Their effectiveness was evaluated in both offline and online integration scenarios. In the offline scenario, all data batches were available from the start, whereas in the online scenario, data batches were observed incrementally. To facilitate this evaluation, we adapted offline integration methods that originally lacked online integration support by incorporating continual learning or fine-tuning based transfer learning techniques.

#### 4.11.1 Horizontal integration methods

##### Geneformer

Geneformer [27] is a large-scale pre-trained model for single-cell RNA sequencing (scRNA-seq) data. The Geneformer Python package is available at https://huggingface.co/ctheodoris/Geneformer. We used a model pre-trained on 30 million cells to predict gene embeddings for each cell and generated cell embeddings by averaging these gene embeddings.

##### PCA

PCA was applied to the dataset after selecting highly variable genes. We then selected the top 32 principal components as the embeddings for further analysis.

#### 4.11.2 Rectangular integration methods

##### SCALEX

SCALEX [13] is an online integration method for scRNA-seq or scATAC-seq data, available at https://github.com/jsxlei/SCALEX. Because it does not support ADT data integration natively, we concatenated RNA and ADT data as input.

##### SCALEX-projection

SCALEX allows the generation of embeddings on new data without training. We trained a SCALEX model with the first data batch and projected the remaining batches using this model, setting the embedding dimension to 32 for consistency.

##### trVAE

trVAE [53] is an RNA data integration method available at https://github.com/theislab/trVAE. Because it supports only single modality input, we concatenated RNA and ADT data for input.

##### trVAE+scArches

Using scArches for transfer learning, we concatenated RNA and ADT profiles for input. We trained a reference model with the first data batch and then trained adaptors with the remaining batches.

##### Concerto

Concerto [12] integrates RNA and ADT data. The code is available at https://github.com/melobio/Concerto-reproducibility.

##### Concerto-transfer

Since Concerto supports transfer learning, we used the first data batch to pretrain the model and then fine-tuned it sequentially on the remaining batches.

##### Symphony

Symphony [14] efficiently handles new data queries. The code is at https://github.com/immunogenomics/symphony. We concatenated RNA and ADT data for input, using the first batch to build a reference model and querying the remaining batches.

##### Symphony-offline

We trained all data batches collectively with Symphony in this experiment.

##### online iNMF

Online iNMF [10] iteratively updates models for single-cell RNA data integration. The code is available at https://github.com/welch-lab/liger. While it does not natively support ADT data, we concatenated RNA and ADT data for input. We integrated data by batch and used the final model to generate embeddings for all data.

##### online iNMF-offline

We used online iNMF to train all data batches together for offline integration.

##### CCA+WNN

Seurat’s CCA [6] is used to remove batch effects in feature space, and the weighted nearest neighbors (WNN) is used for fusing omics. We first used the FindIntegrationAnchors function (reduction=‘cca’) in the Seurat package to obtain anchors of batches for each modality. Then we used the IntegrateData function to correct batch effects. After that, we calculated PCA on the integrated data and used the WNN strategy to gain the graph output.

##### CCA+WNN-continual

We integrated new data batches with previously corrected counts, continually updating the dataset. However, efficiency decreases as integrations accumulate, resembling an online integration method based on continual learning with unbounded rehearsal memory.

##### RPCA+WNN

This method is similar to CCA+WNN, except the parameter reduction was set to ‘rpca’ in the FindIntegrationAnchors function.

##### RPCA+WNN-continual

This method is similar to CCA+WNN-continual, except the parameter reduction was set to ‘rpca’ in the FindIntegrationAnchors function.

##### sciPENN

sciPENN [34] integrates RNA and ADT data and is available at https://github.com/jlakkis/sciPENN. Since sciPENN does not handle missing genes, we used the intersection of RNA features and the union of ADT features across batches.

##### sciPENN-transfer

Since sciPENN does not support direct transfer learning, we trained the model on the initial batch, fine-tuned it on subsequent batches, and used the final model to predict latent variables for the entire dataset.

#### 4.11.3 Mosaic integration methods

##### MIRACLE

MIRACLE uses MIDAS [19] as the base model, adopting the same model architecture and training hyperparameters as MIDAS. We trained MIRACLE batch by batch and used the final trained model to generate embeddings for all data.

##### MIRACLE-offline

MIRACLE-offline uses the same base model as MIRACLE. For this experiment, all data batches were collectively input for training.

##### MIRACLE-transfer

MIRACLE-transfer uses the same base model as MIRACLE. The model was fine-tuned using new data.

##### Multigrate

Multigrate [54] is a multimodal mosaic integration method for RNA, ADT, and ATAC data. The code is available at https://github.com/theislab/multigrate. Hyperparameters KL and integ were set to 0.1 and 3000, respectively.

##### Multigrate+scArches

Multigrate+scArches [11] was implemented with scArches https://github.com/theislab/scarches for transfer learning. For WNN rectangular dataset, the first data batch was used as reference data to construct the initial model, and subsequent batches trained adaptors. For DOTEA mosaic dataset, because the model cannot dynamically adapt to new modalities, the first two batches were used as reference data, encompassing all three modalities.

##### scVAEIT

scVAEIT [35] is a multimodal mosaic integration method for RNA, ADT, and ATAC data. The code is available at https://github.com/jaydu1/scVAEIT.

##### scVAEIT-transfer

Since scVAEIT lacks transfer learning support, we extended it for this purpose. The model was initially trained on the first batch and then fine-tuned on subsequent batches.

##### totalVI

totalVI [7] integrates RNA and ADT data. The Python package is at https://github.com/YosefLab/totalVI_reproducibility and is incorporated into scArches.

##### totalVI+scArches

totalVI is implemented with scArches for transfer learning as well. The first data batch built the initial model and subsequent batches trained adaptors.

### 4.12 Evaluation metrics for integration

To comprehensively evaluate the integration performance of MIRACLE and various state-of-the-art methods, we used scIB and scMIB metrics. For horizontal and rectangular integration tasks, we used the scIB metric to assess batch correction and biological conservation performance. For mosaic integration tasks, we used the scMIB metric to evaluate batch correction, modality alignment, and biological conservation performance. Modality alignment is crucial for mosaic integration as it reflects the model’s capabilities in applications such as modality imputation and cross-modal label transfer. Note that for other mosaic integration methods compared, batch correction can only be performed in the embedding space rather than the feature space, so we excluded feature space-based metrics for batch correction and biological conservation in scMIB.

Specifically, the batch correction metrics include graph integration local inverse Simpson’s index (iLISI) *y*^iLISI^, graph connectivity *y*^gc^, and kNN batch effect test (kBET) *y*^kBET^. The modality alignment metrics include modality averaged silhouette width (ASW) *y*^ASW^, fraction of samples closer than the true match (FOSCTTM) *y*^FOSCTTM^, label transfer F1 *y*^ltF1^, ATAC area under the receiver operating characteristic (AUROC) *y*^AUROC^, RNA Pearson’s *r y*^RNAr^ and ADT Pearson’s *r y*^ADTr^. The biological conservation metrics include normalized MI (NMI) *y*^NMI^, adjusted Rand index (ARI) *y*^ARI^, isolated label F1 *y*^ilF1^ and graph cell-type LISI (cLISI) *y*^cLISI^. The definitions of these metrics are detailed in Supplementary Note 2.

We average each type of metrics to obtain its comprehensive score, including the batch correction score *y*^batch^, modality alignment score *y*^mod^, and biological conservation score *y*^bio^:

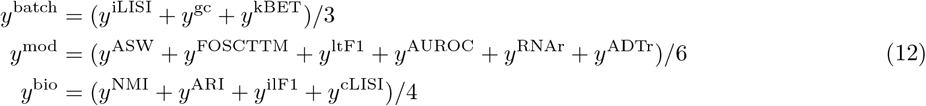

Following Luecken et al. [26], the scIB overall score *y*^scIB^ is the sum of *y*^batch^ weighted by 0.4 and *y*^bio^ weighted by 0.6:

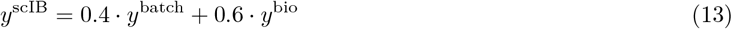

As an extension of scIB, following He et al. [19], the scMIB overall score *y*^scMIB^ is the sum of *y*^batch^ weighted by 0.3, *y*^mod^ weighted by 0.3, and *y*^bio^ weighted by 0.4:

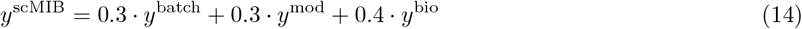

## Supplementary Notes

### Supplementary Note 1: Details of Ball Tree sampling

Due to the high dimensionality and noise in the observed single-cell data 𝒳 = {***x***_1_, …, ***x***_*N*_ }, we first obtain their low-dimensional cell embeddings Ƶ = {***z***_1_, …, ***z***_*N*_ }. We then use the Ball Tree algorithm from the scikit-learn [55] Python package to perform a multi-resolution partitioning of Ƶ, resulting in an *L*-layer binary tree 𝒯. The *i*-th tree at layer *l*, denoted as 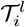, can be represented as:

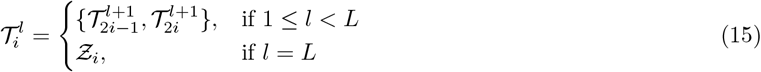

where at the top layer, 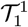 represents the entire tree, while at the bottom layer, 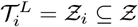 represents the subtree containing the smallest subset of embeddings.

Based on the Ball Tree partitioning, we can sample at each layer. For layer *l*, we only need to randomly select one cell from each subtree 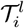 to achieve distribution-preserving sampling at that layer’s resolution. Therefore, the larger the *l*, the higher the resolution and the more subtrees there are, resulting in more cells being collected. For a given sampling ratio *γ*, we conduct sampling from the bottom layer to the top layer, with resolution decreasing from high to low, until the proportion of sampled data reaches *γ*. See Algorithm 2 for details.

#### Algorithm 2

The Ball Tree sampling algorithm.

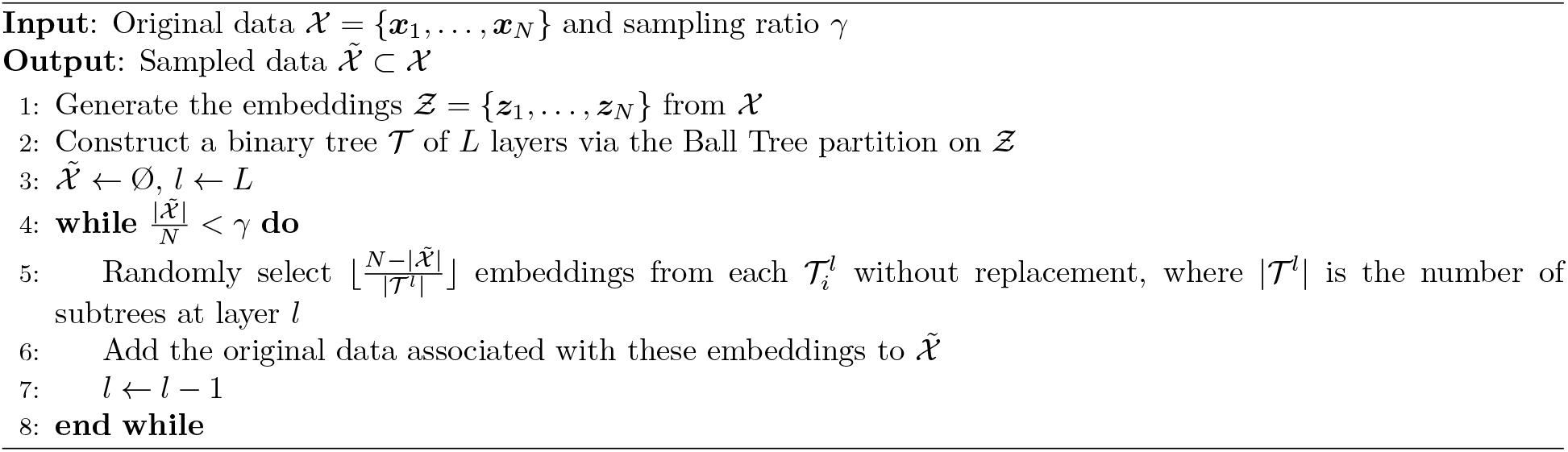

### Supplementary Note 2: Detailed definitions of integration metrics

We used the scIB and scMIB metrics to assess the integration performance of different methods. The detailed definitions of these metrics for batch correction, modality alignment, and biological conservation performance are as follows.

#### Batch correction metrics

The batch correction metrics include graph iLISI *y*^iLISI^, graph connectivity *y*^gc^, and kBET *y*^kBET^.

#### Graph iLISI

The graph iLISI metric, derived from iLISI [56], measures batch mixing by computing the inverse Simpson’s index for diversity in kNN graphs. It estimates the effective number of batches in the neighborhood, ranging from 1 to *N*, where *N* is the number of batches. Scores close to the actual batch number indicate good mixing. Unlike the typical iLISI, which is unsuitable for graph-based outputs, scIB’s graph iLISI uses a graph-based distance metric to determine the nearest neighbors, avoiding skews in graph-based integration. The graph iLISI scores range from 0 to 1, with 0 indicating strong separation and 1 indicating perfect mixing.

#### Graph connectivity

Graph connectivity, proposed by scIB, checks if cells with the same label are connected in the kNN graph. For each label *c*, it calculates the largest connected component size of *c*-labeled cells divided by the total number of *c*-labeled cells. The average of these values across all labels represents the total graph connectivity, ranging from 0 to 1. A score of 1 means all same-label cells from different batches are connected in the integrated kNN graph, indicating perfect batch mixing.

#### kBET

The kBET [57] assesses local batch mixing in the kNN graph by repeatedly testing random cell fractions to see if their local label distributions match the global distribution (null hypothesis). The kBET value is the rejection rate of this hypothesis across neighborhoods, with values near 0 indicating well-mixed batches. scIB modifies kBET with a diffusion-based correction for unbiased comparison of graph- and non-graph-based integration results. kBET values are computed for each label, averaged, and subtracted from 1 to obtain the final kBET score.

#### Modality alignment metrics

The modality alignment metrics include modality ASW, FOSCTTM, label transfer F1, ATAC AUROC, RNA Pearson’s *r*, and ADT Pearson’s *r*.

#### Modality ASW

The modality ASW measures the alignment of different modality embeddings’ distributions. Originally, ASW [58] assessed cluster separation. In scIB, ASW is adapted to evaluate batch effect removal, yielding a batch ASW score from 0 to 1, where 0 indicates strong batch separation and 1 indicates perfect batch mixing. Similarly, replacing batch embeddings with modality embeddings allows us to define a modality ASW, where 0 denotes strong modality separation and 1 denotes perfect modality alignment. In our comparison methods, modality embeddings are generated by processing each modality individually through the trained model.

#### FOSCTTM

The FOSCTTM [59] measures the alignment between different modality embeddings. For a modality pair {*m*_1_, *m*_2_}, the FOSCTTM is calculated as follows:

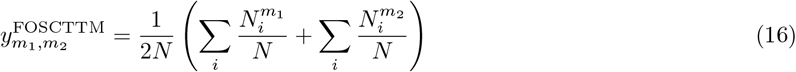

where:

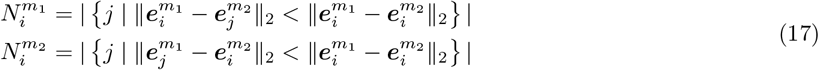

Here, *N* is the number of cells, *i* and *j* are cell indices, and 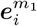 and 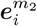 are the embeddings of cell *i* in modalities *m*_1_ and *m*_2_, respectively. 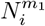 counts how many cells in modality *m*_2_ that are closer to 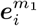 than 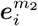 is, and similarly for 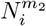. To compute the final FOSCTTM score, we first obtain the embeddings for each modality, calculate the FOSCTTM values for every modality pair and then average these values and subtract the result from 1. A higher FOSCTTM score indicates better modality alignment.

#### Label transfer F1

Label transfer F1 measures the alignment of cell types between different modality embeddings. It evaluates if cell-type labels can be transferred between modalities without bias. For each modality pair, we construct a kNN graph using their embeddings and transfer labels from one modality to the other based on nearest neighbors. The transferred labels are then compared to the original labels using the micro F1 score, termed the label transfer F1. The final score is the average F1 score across all modality pairs.

#### ATAC AUROC

The ATAC AUROC measures the alignment of different modalities in the ATAC feature space, previously used to evaluate ATAC prediction quality [60]. For each method, we convert different non-ATAC modality combinations into ATAC features, calculate the AUROC using true ATAC features as the ground truth, and average these AUROCs for the final score. For example, with ATAC, RNA, and ADT data, we evaluate combinations {RNA}, {ADT} and {RNA, ADT}. Each combination is processed by the model to generate imputed data of all modalities {ATAC, RNA, ADT}, with the generated ATAC features used for AUROC calculation.

#### RNA Pearson’s *r*

The RNA Pearson’s *r* value measures the alignment of different modalities in the RNA feature space. For each method, we convert different non-RNA modality combinations into RNA features, calculate the Pearson’s *r* value between each converted result and the true RNA features, and average these values for the final score.

#### ADT Pearson’s *r*

The ADT Pearson’s *r* value measures the alignment of different modalities in the ADT feature space, calculated similarly to the RNA Pearson’s *r* value.

#### Biological conservation metrics

The biological conservation metrics include NMI, ARI, isolated label F1 and graph cLISI.

#### NMI

The NMI measures the similarity between predefined cell-type labels and clustering results from embeddings or graphs. Optimized Louvain clustering is used as per scIB. NMI scores range from 0 to 1, with 0 indicating no correlation and 1 indicating a perfect match.

#### ARI

The ARI measures the overlap between two clustering results. The Rand Index (RI [61]) considers both cell pairs in the same clusters and those in different clusters in the predicted (Louvain clustering) and true (cell-type) clusters. The ARI corrects the RI for randomly correct labels, with 0 indicating random labeling and 1 representing a perfect match.

#### Isolated label F1

scIB proposes the isolated label F1 score to evaluate integration performance, focusing on cells with labels shared by few batches. Labels present in the fewest batches are identified as isolated labels.The isolated label F1 score measures clustering performance on these labels, ranging from 0 to 1, with 1 indicating perfect separation of isolated label cells into one cluster.

#### Graph cLISI

The graph cLISI is similar to the graph iLISI but focuses on cell-type labels instead of batch labels. While iLISI highlights group mixing, cLISI values group separation [56]. The graph-adjusted cLISI ranges from 0 to 1, with 0 indicating low cell-type separation and 1 indicating strong cell-type separation.

**Supplementary Fig. 1:**
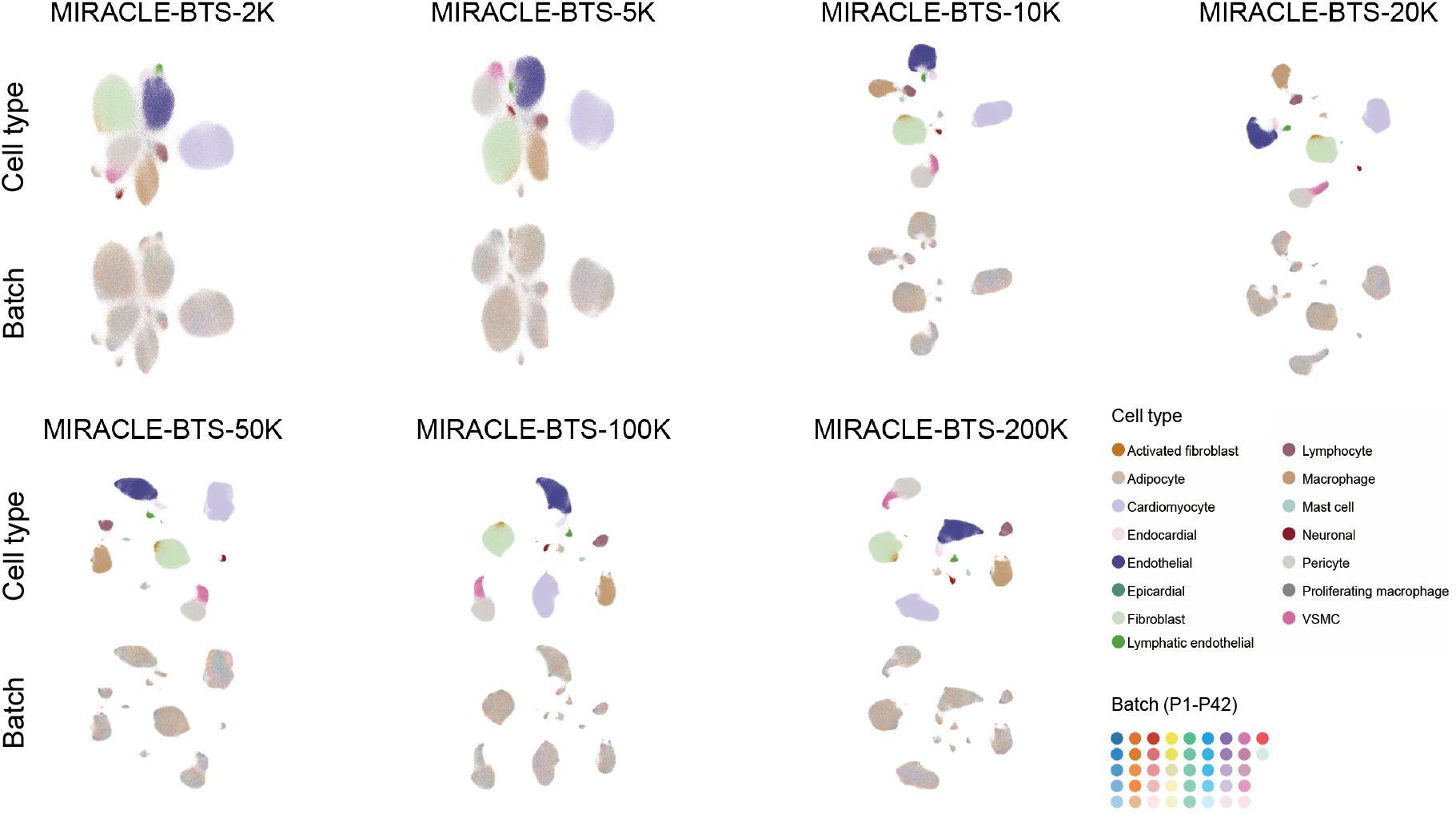
UMAP visualization of the biological states inferred by MIRACLE-BTS under various memory capacity conditions.

**Supplementary Fig. 2:**
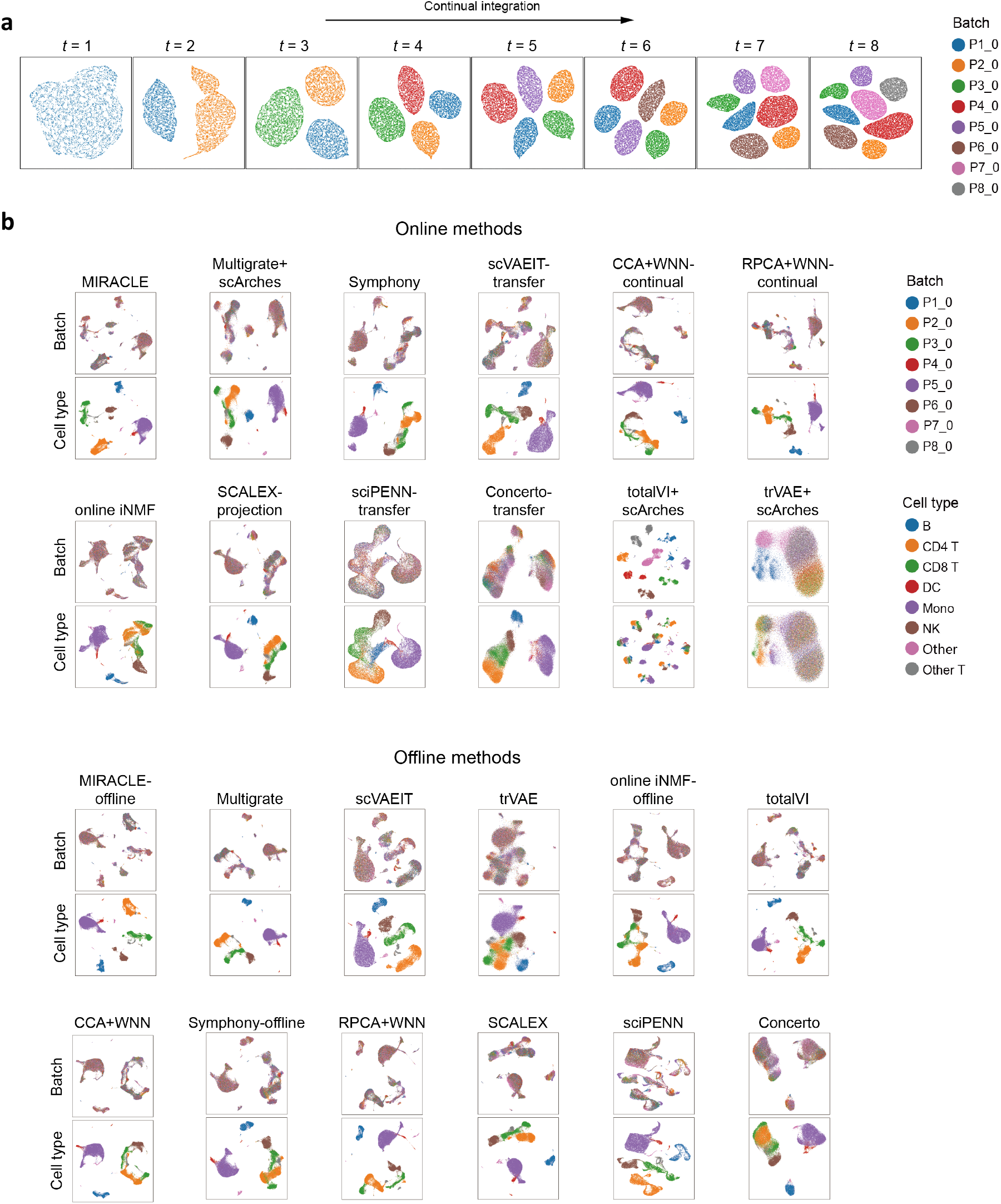
UMAP visualization results of MIRACLE and other methods on the WNN bimodal dataset for integration across batches and cell types. **a**, UMAP visualization of the technical noise inferred by MIRACLE at different time points. **b**, UMAP visualization comparing the biological states generated by MIRACLE and other state-of-the-art single-cell integration methods, in both online (upper panel) and offline (lower panel) scenarios.

**Supplementary Fig. 3:**
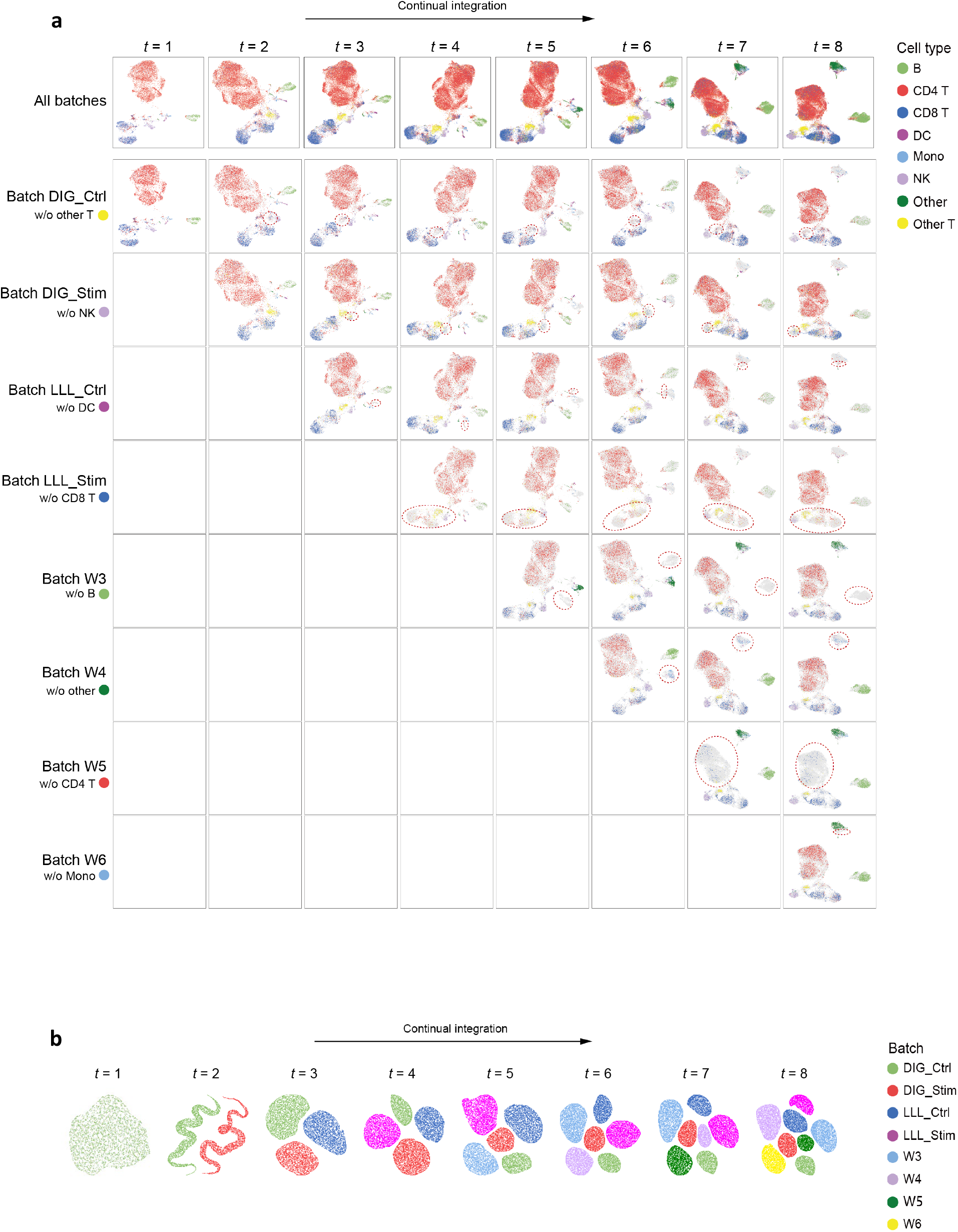
UMAP visualization of the cell embeddings inferred by MIRACLE on the DOTEA trimodal dataset during continual mosaic integration. **a**, UMAP visualization of the biological states inferred by MIRACLE at different time points, with each column corresponding to a specific time point. The first row shows the integration of all available data over time, with the dataset size increasing at each time point. Each row from the 2nd to the 9th displays the integration results for each batch at different time points, with missing cell type regions highlighted by red circles. Cells are colored by cell type. **b**, UMAP visualization of the technical noise inferred by MIRACLE at different time points.

**Supplementary Fig. 4:**
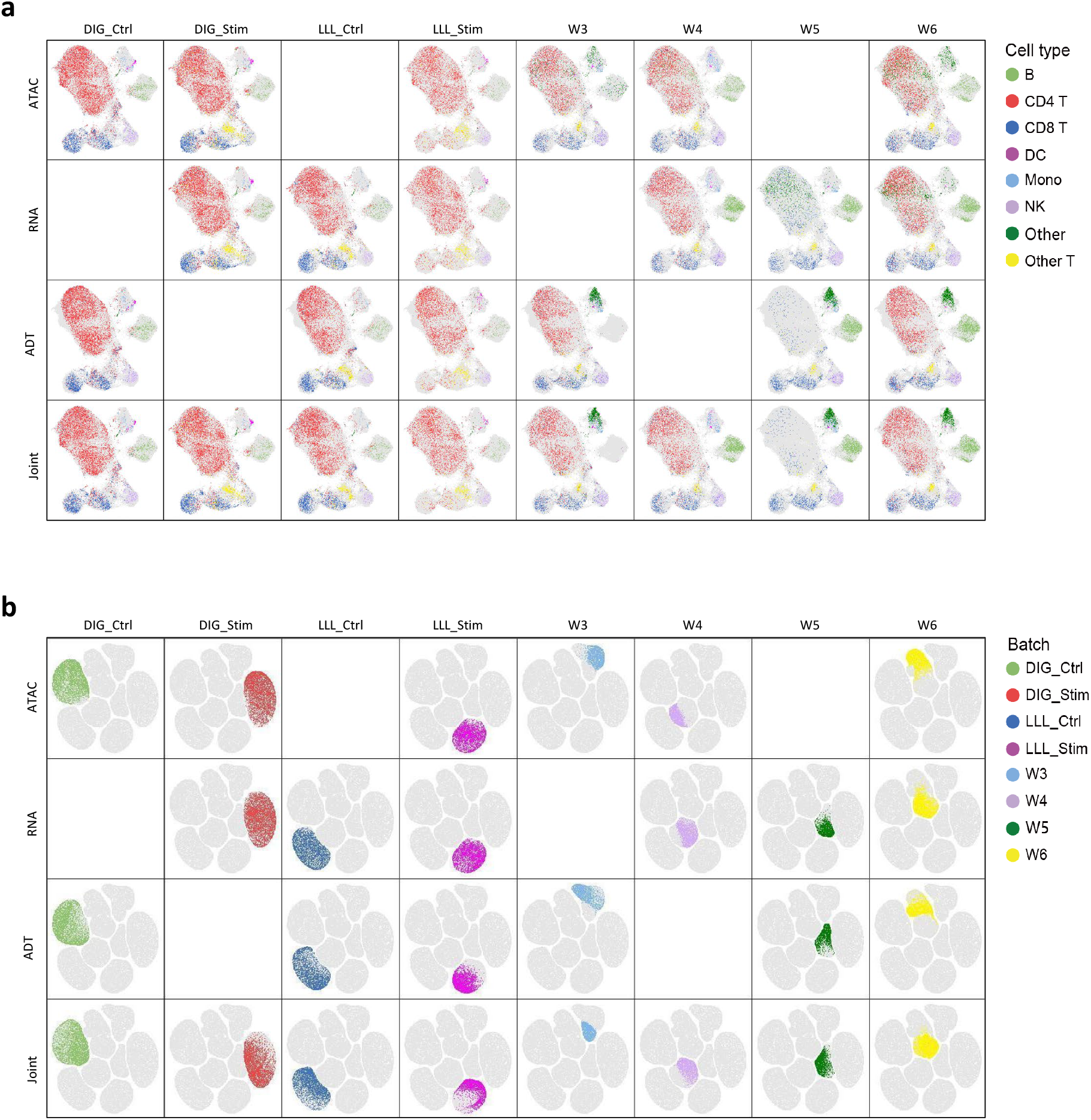
UMAP visualization of the biological states (a) and technical noise (b) inferred by MIRACLE on the DOTEA trimodal dataset in continual mosaic integration tasks. In each panel, the UMAPs are vertically split by modality and horizontally split by batch.

**Supplementary Fig. 5:**
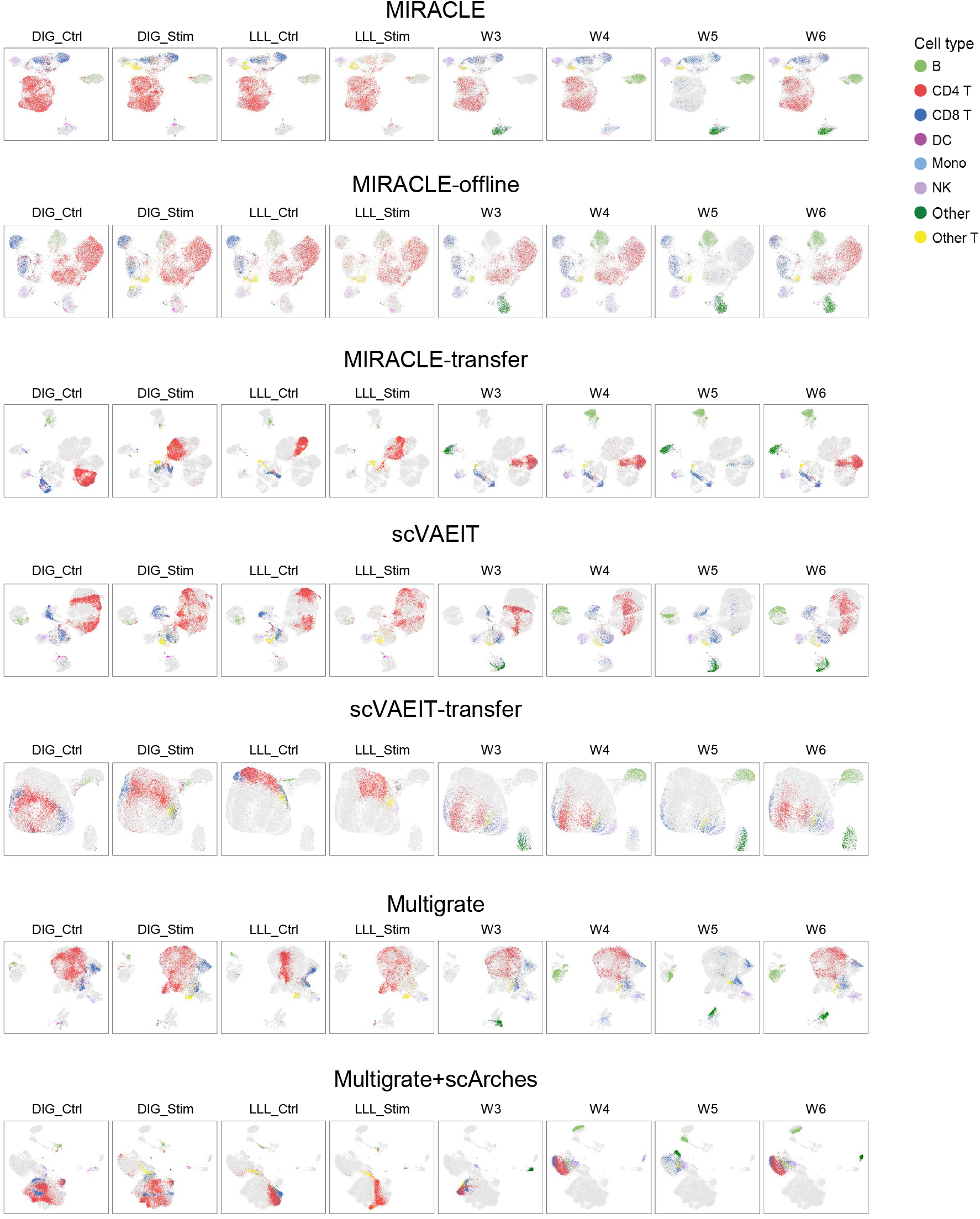
UMAP visualization results of MIRACLE and other methods on the DOTEA trimodal dataset in continual mosaic integration tasks. In each panel, the UMAPs are split by batch.

**Supplementary Fig. 6:**
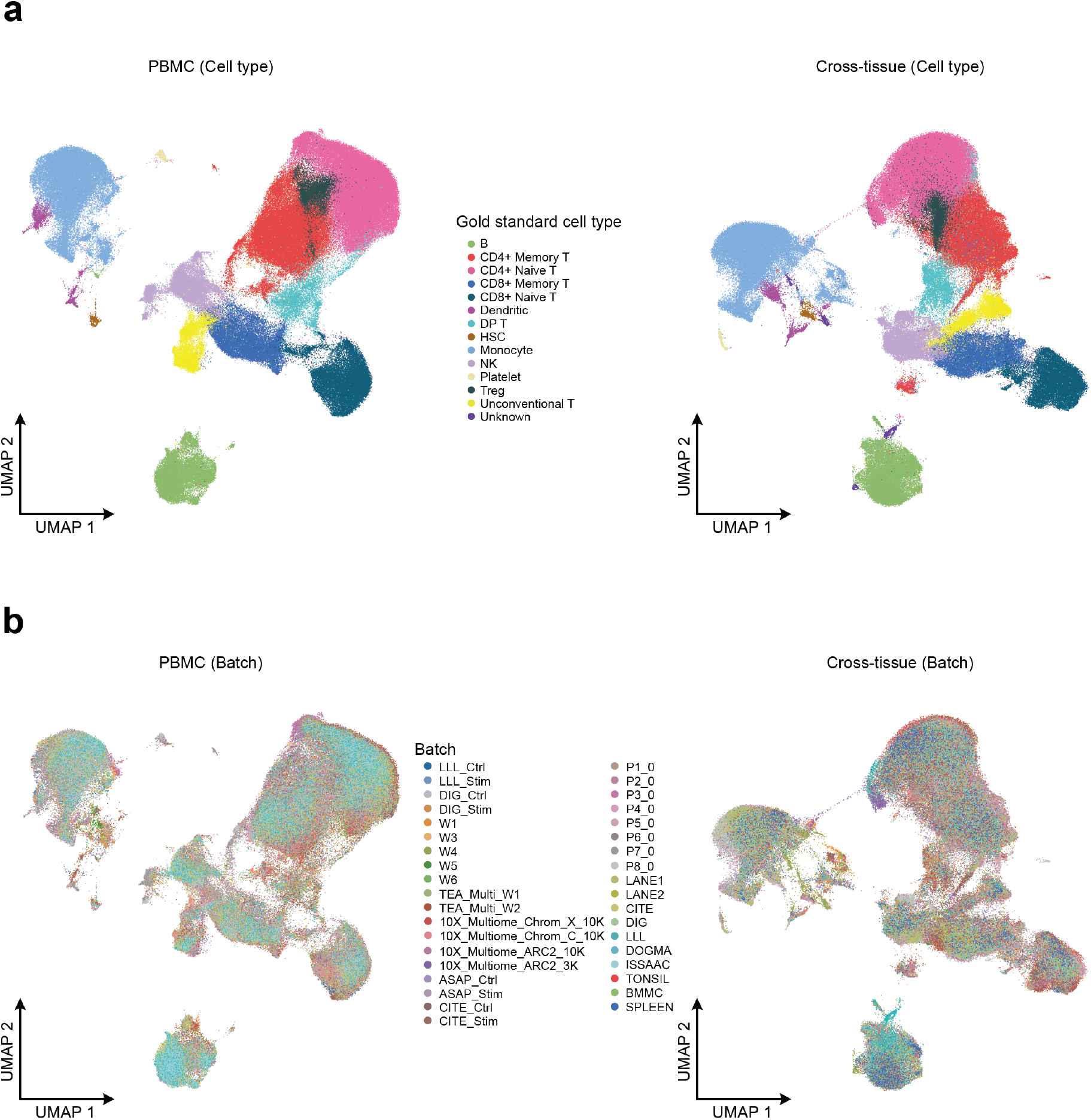
UMAP visualization (merged view) of the biological states inferred by MIRACLE-offline, with the PBMC mosaic datasets containing 34 batches (left panel) and cross-tissue mosaic datasets containing 37 batches (right panel). Cells are colored by cell type (a) and batch (b), where the cell types are derived from MIRACLE-offline.

**Supplementary Fig. 7:**
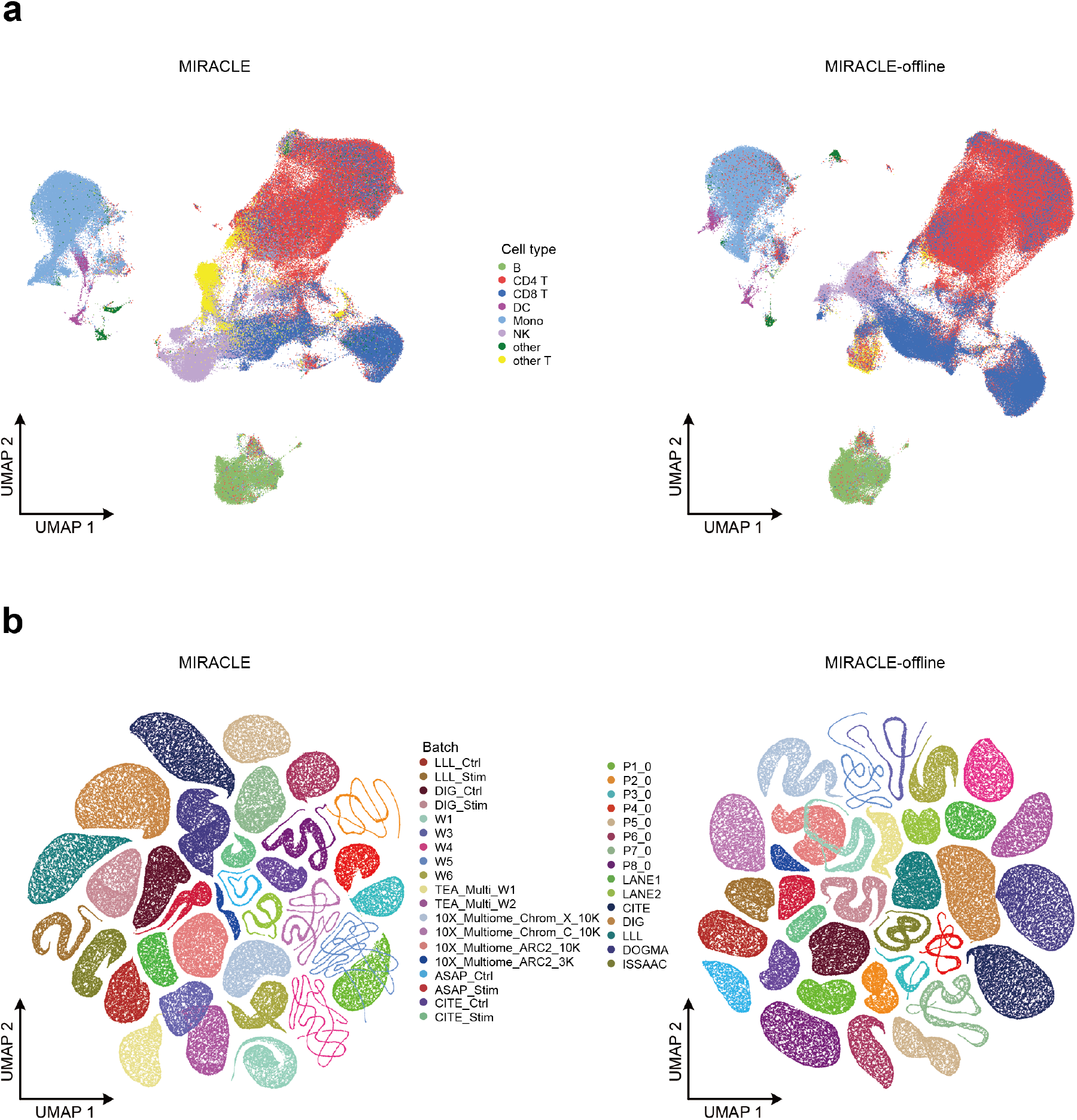
UMAP visualization (merged view) of the biological states (a) and technical noise (b) inferred by MIRACLE (left panels) and MIRACLE-offline (right panels) on the PBMC trimodal mosaic dataset containing 34 batches. Cells are colored by cell type (a) and batch (b), where the cell types are derived from Seurat labeling.

**Supplementary Fig. 8:**
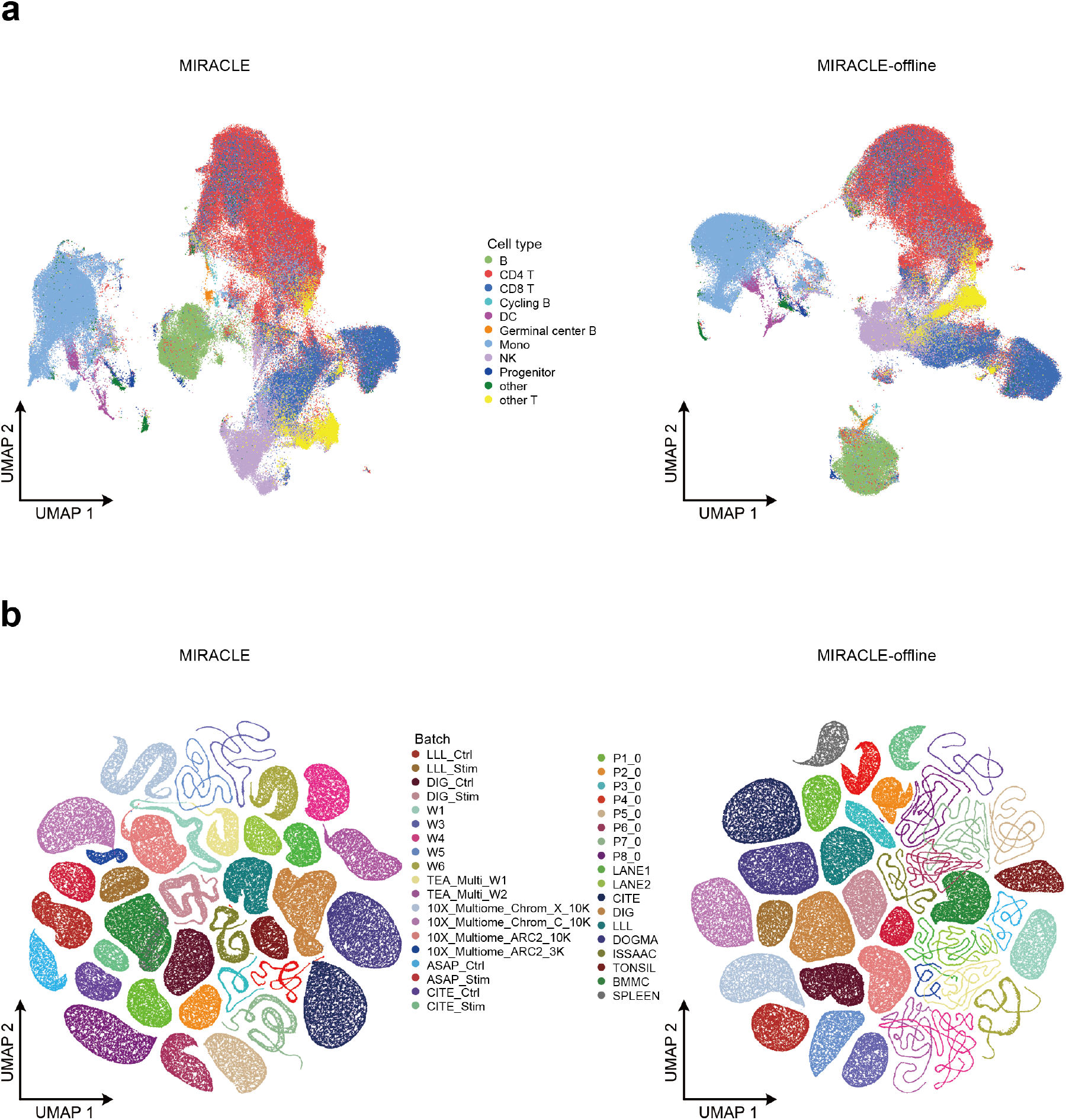
UMAP visualization (merged view) of the biological states (a) and technical noise (b) inferred by MIRACLE (left panel) and MIRACLE-offline (right panel) on the cross-tissue (PBMC, tonsil, BMMC and spleen) trimodal mosaic dataset containing 37 batches. Cells are colored by cell type (a) and batch (b), where the cell types are derived from Seurat and SingleR labeling.

**Supplementary Fig. 9:**
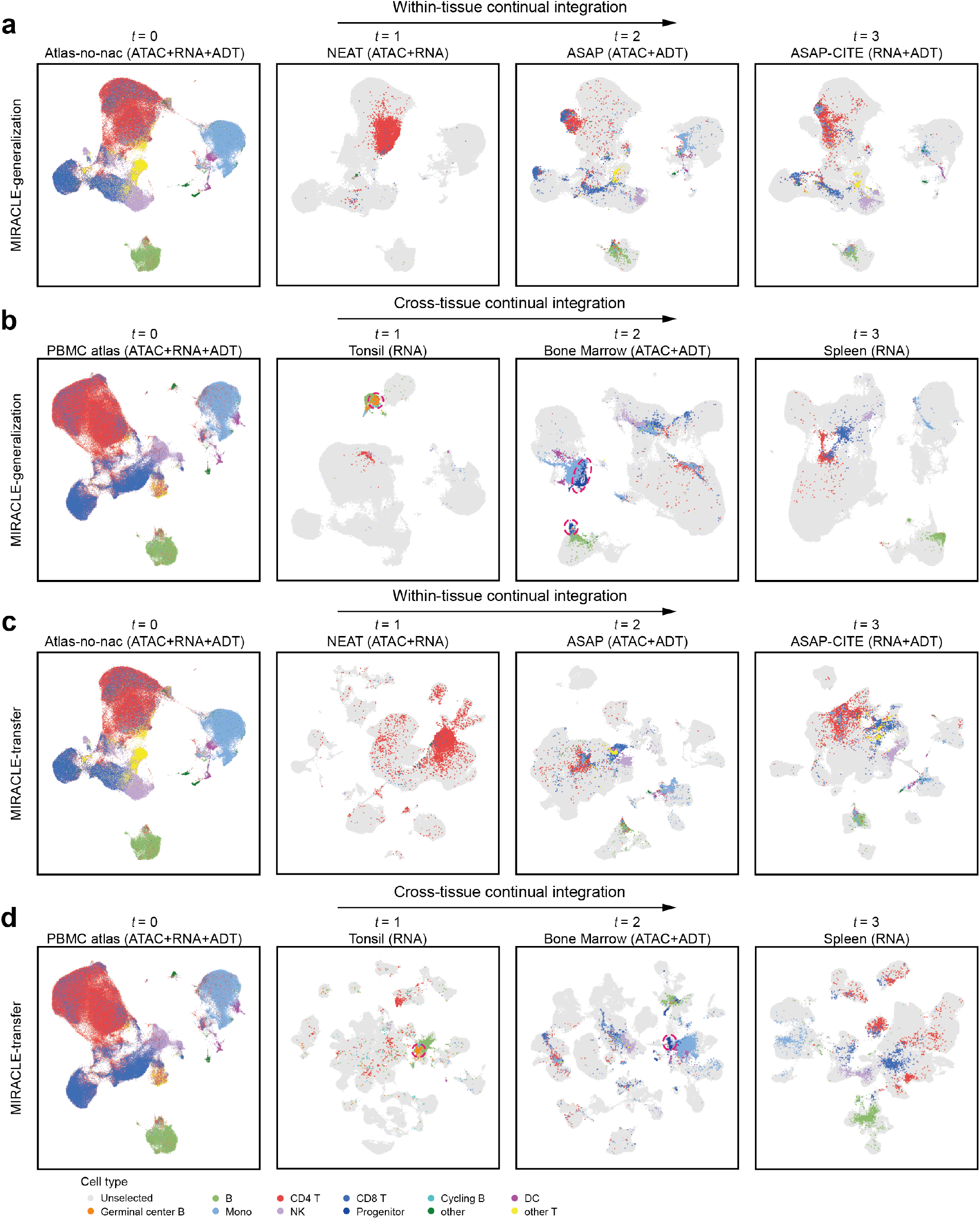
**UMAP visualization of the biological states inferred by MIRACLE-generalization (a,b) and MIRACLE-transfer (c,d), showing the process of within-tissue (a,c) and cross-tissue (b,d) continual atlas construction. Novel cell types are highlighted by red circles.**

**Supplementary Fig. 10:**
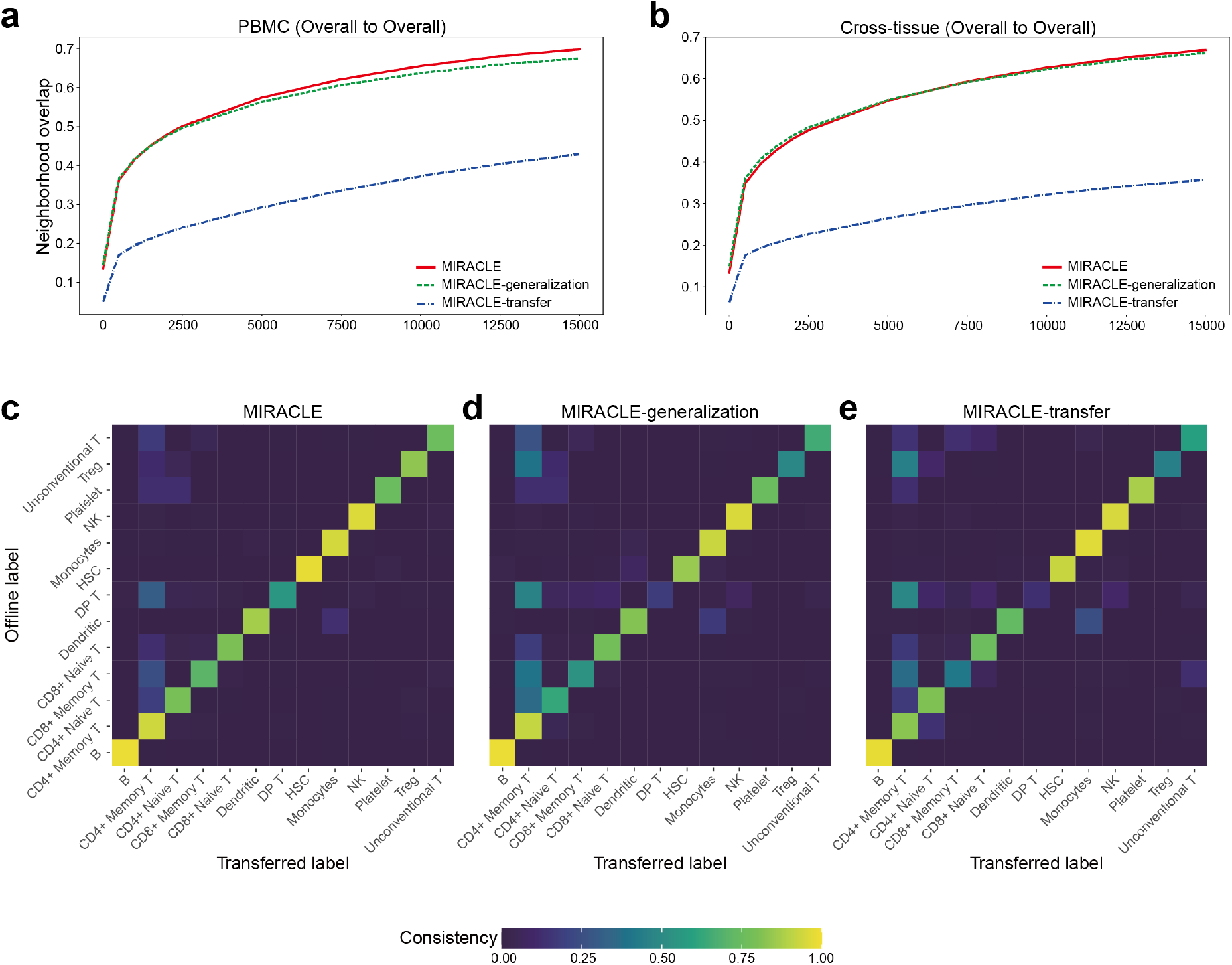
Neighborhood overlap analysis and label transfer confusion plots showing the consistency between MIRACLE’s online and offline integration results. **a, b**, Neighborhood overlap analysis between the dimensionality reduction results of online strategies and MIRACLE-offline across different neighborhood sizes in within-tissue (a) and cross-tissue (b) continual integration tasks, measured by the identification overlap of overall cells’ nearest neighbors. **c-e**, Label confusion plots comparing transfer consistency for MIRACLE (c), MIRACLE-generalization (d), and MIRACLE-transfer (e) in the within-tissue continual integration task.

**Supplementary Fig. 11:**
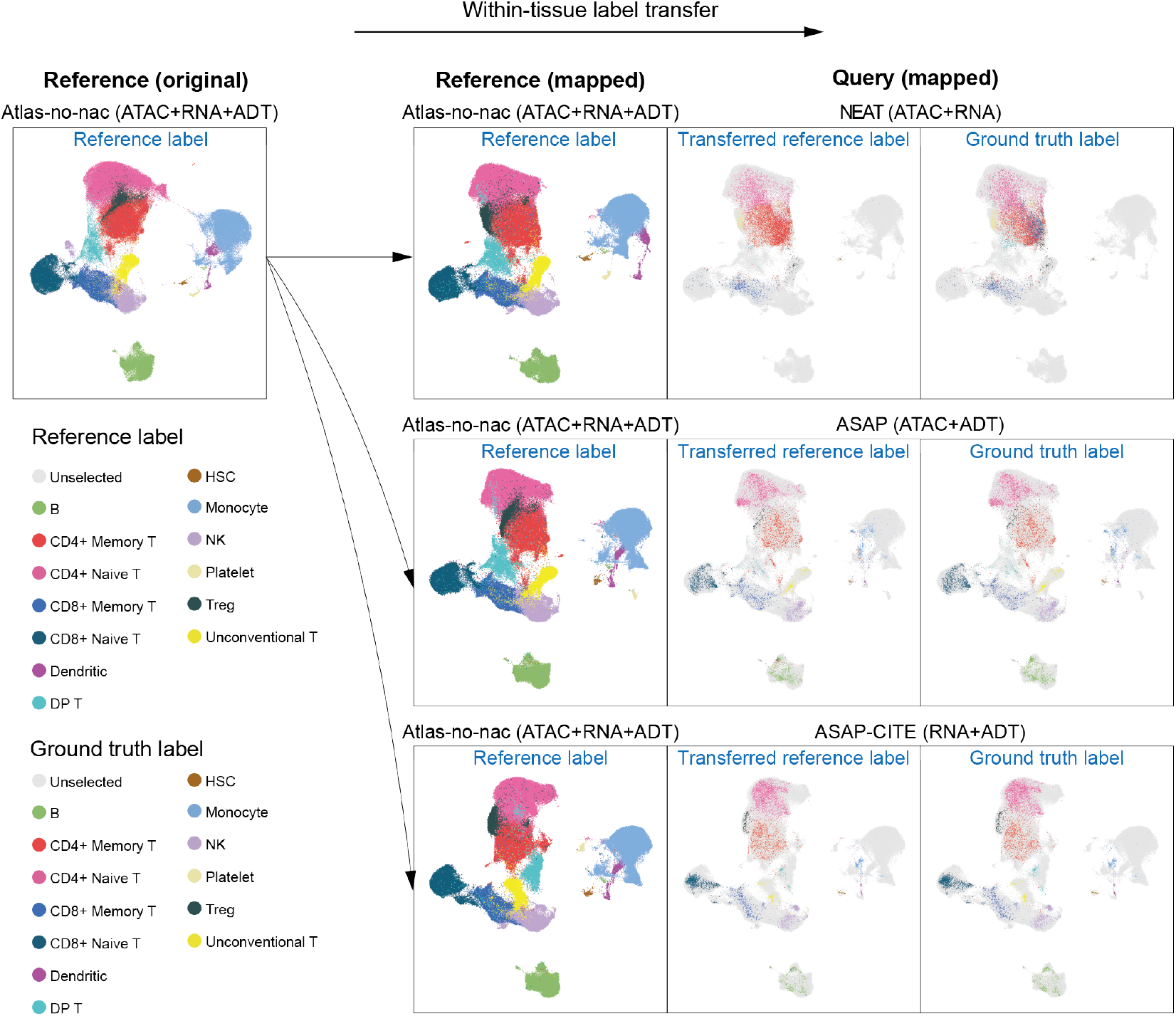
UMAP visualization of the biological states inferred by MIRACLE for within-tissue label transfer. Column 1 shows the original PBMC reference, while Column 2 shows the mapped PBMC references after separately incorporating different PBMC queries, both colored by reference label. Columns 3 and 4 show the three mapped PBMC queries, colored by transferred reference label (Column 3) and ground truth label (Column 4).

**Supplementary Fig. 12:**
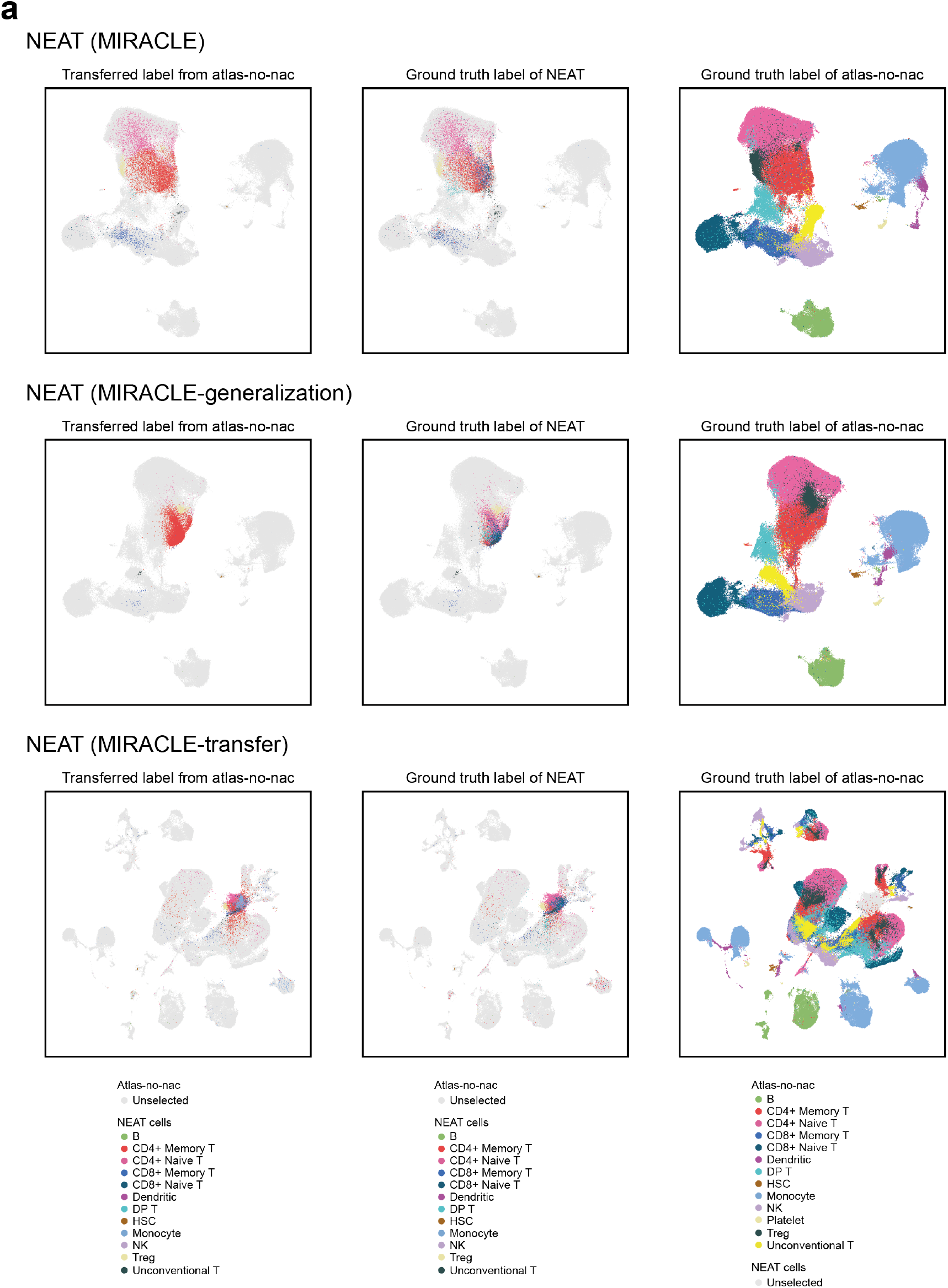
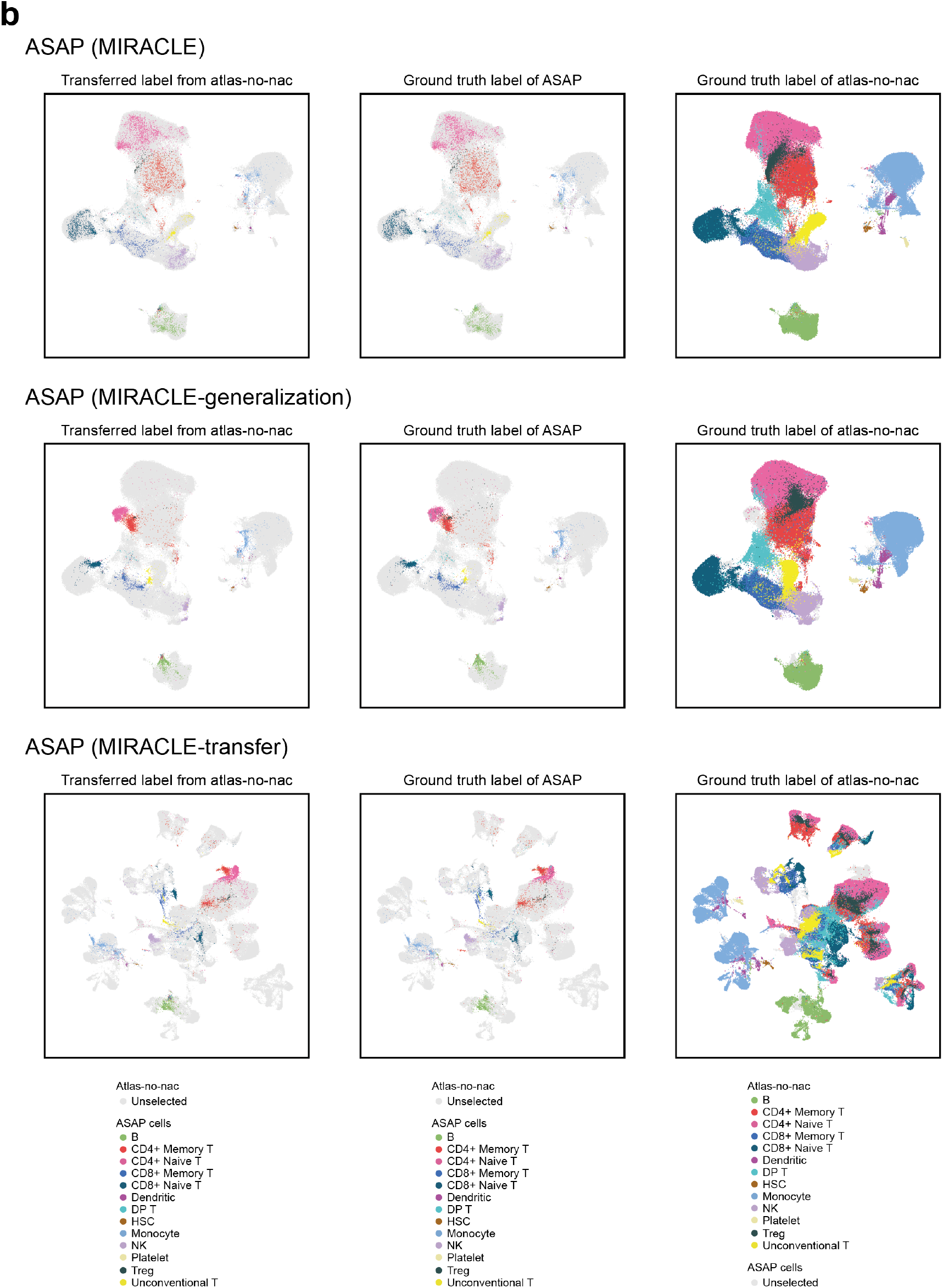
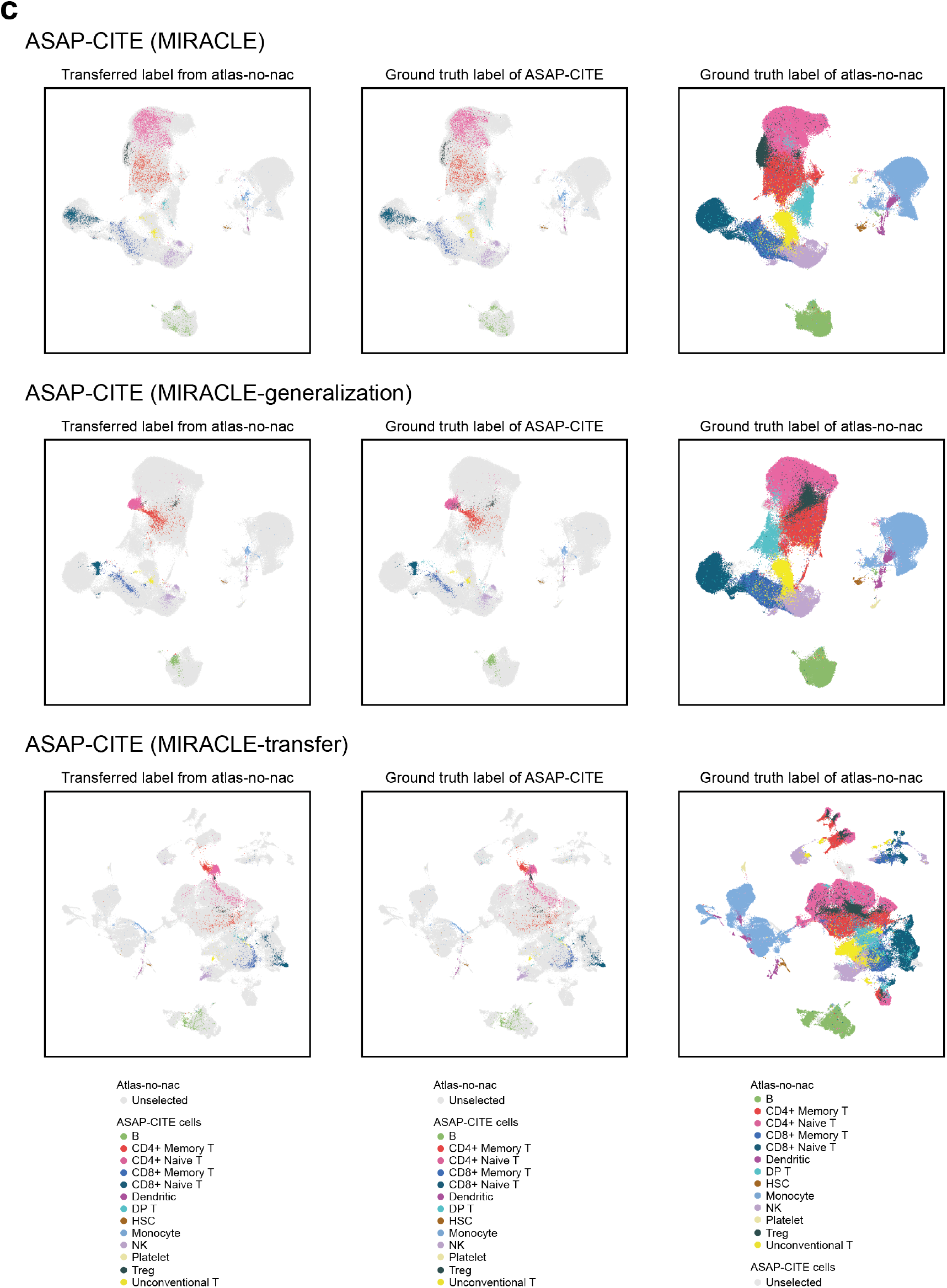
UMAP visualization of the biological states inferred by different online strategies of MIRACLE for within-tissue label transfer. Columns 1 and 2 show the mapped PBMC queries, colored by transferred reference label (Column 1) and ground truth label (Column 2). Column 3 shows the mapped PBMC references after incorporating the PBMC queries, colored by reference label.

**Supplementary Fig. 13:**
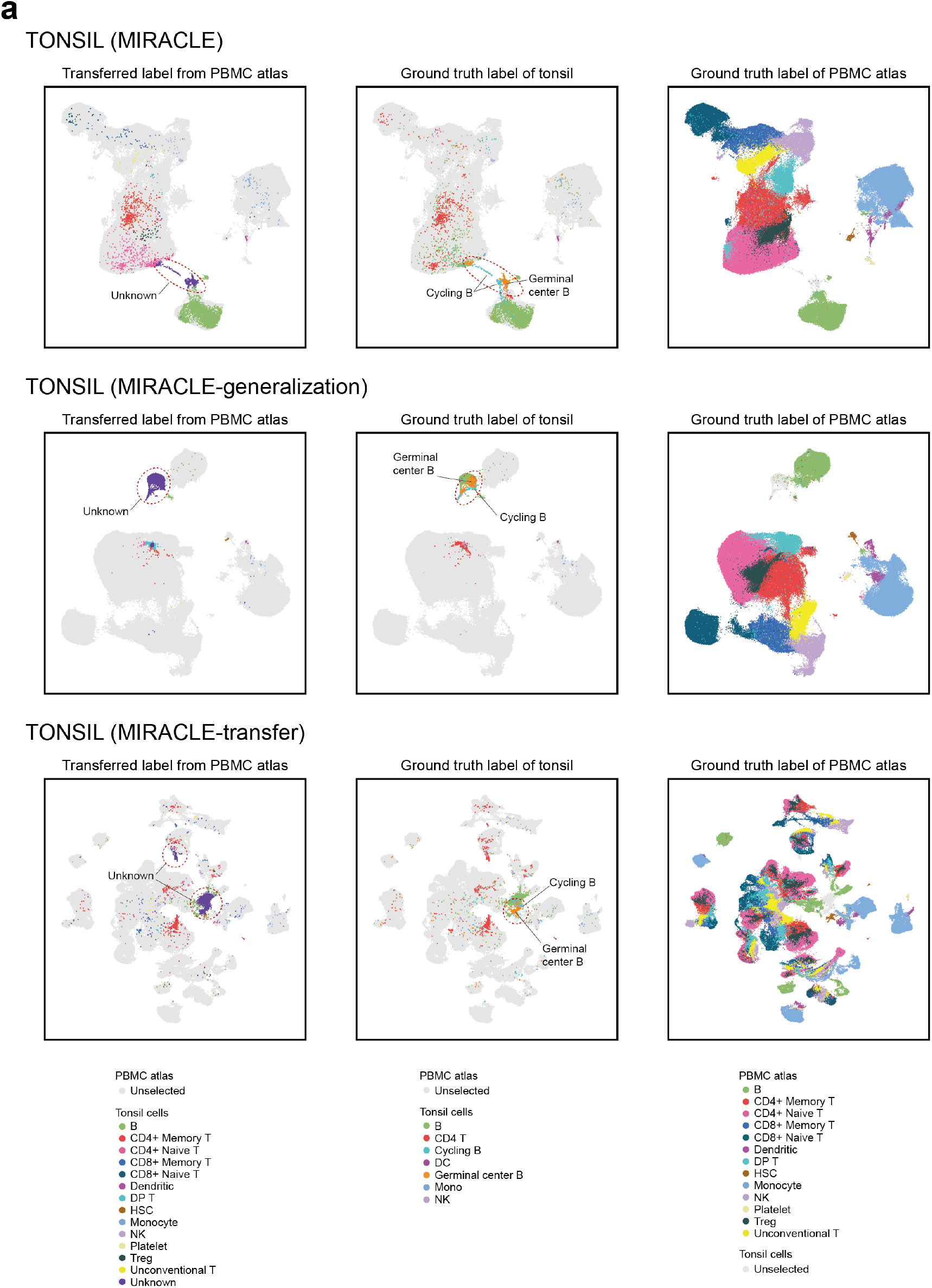
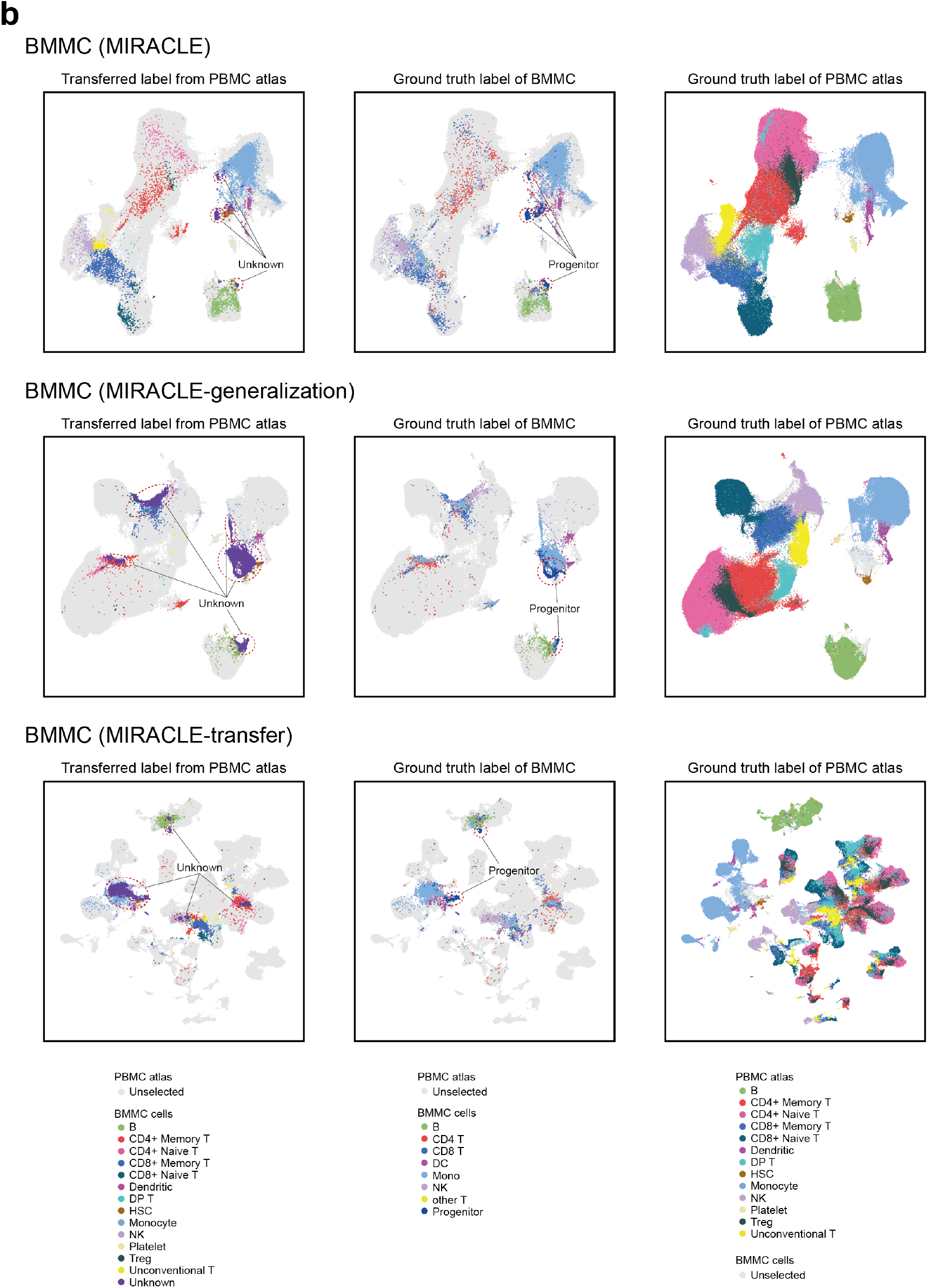
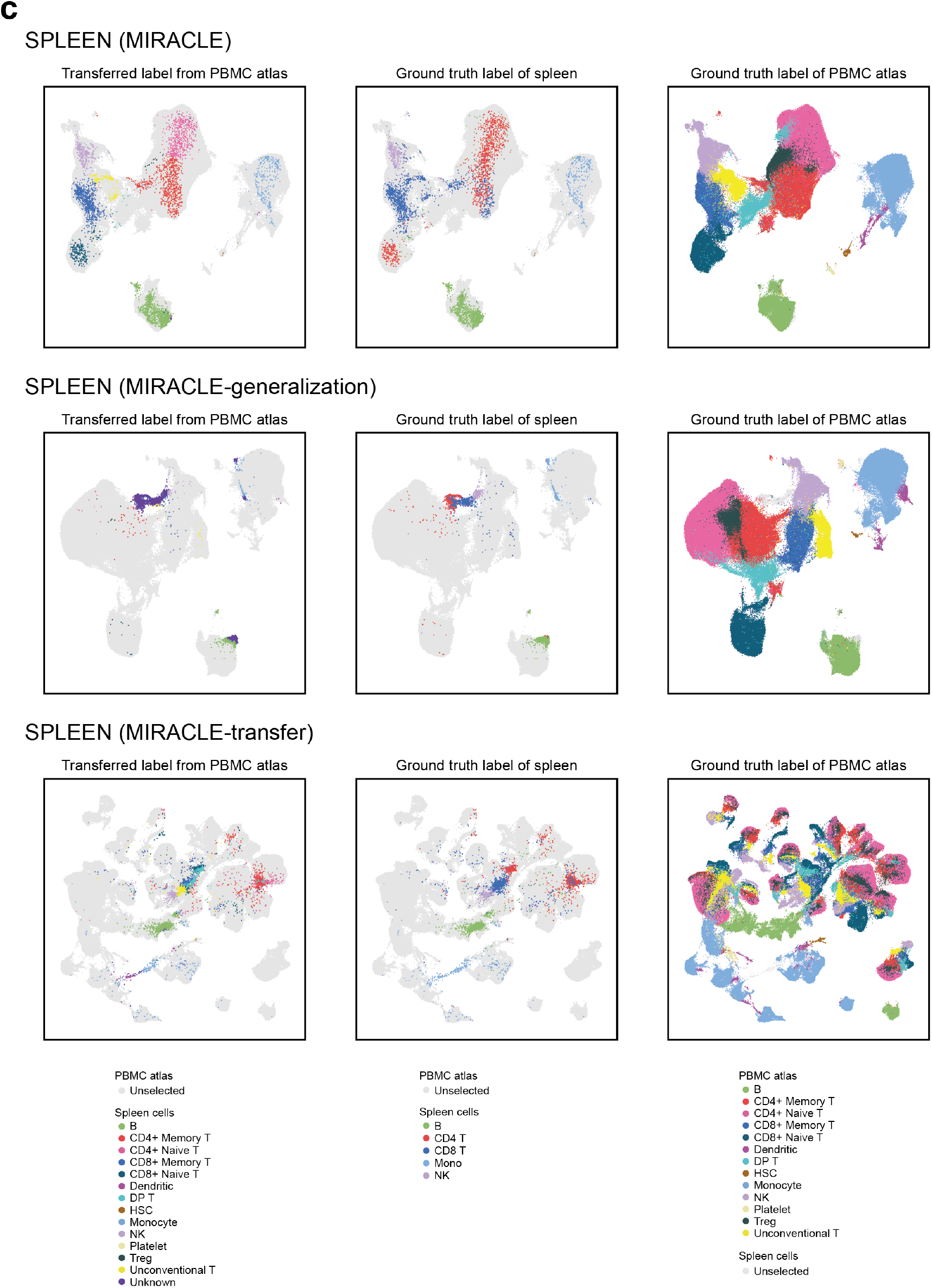
UMAP visualization of the biological states inferred by different online strategies of MIRACLE for cross-tissue label transfer. Columns 1 and 2 show the mapped cross-tissue queries, colored by transferred reference label (Column 1) and ground truth label (Column 2). Column 3 shows the mapped PBMC references after incorporating the cross-tissue queries, colored by reference label.

**Supplementary Fig. 14:**
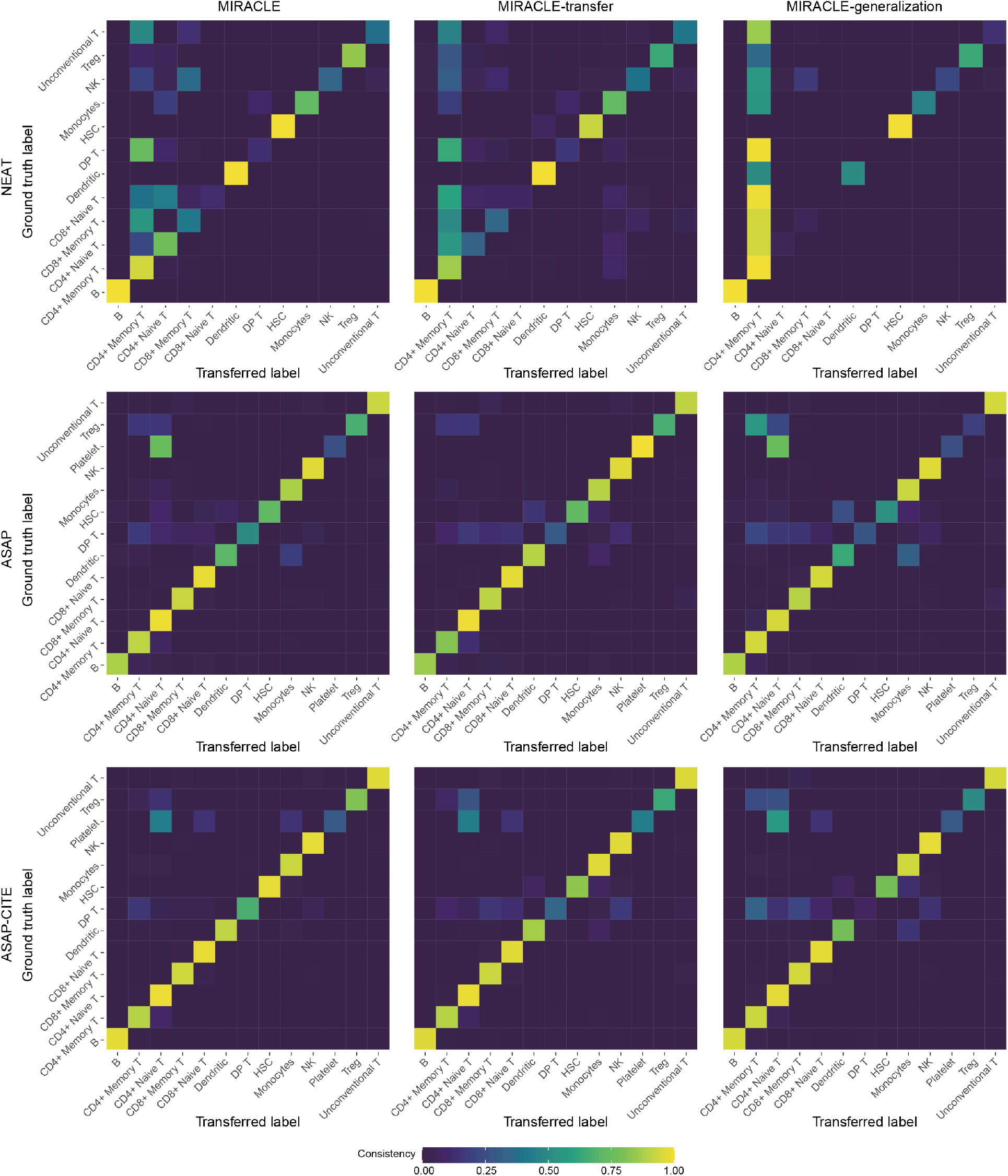
Confusion plots corresponding to the label transfer micro F1-scores for MIRACLE and two other strategies in within-tissue label transfer tasks.

**Supplementary Fig. 15:**
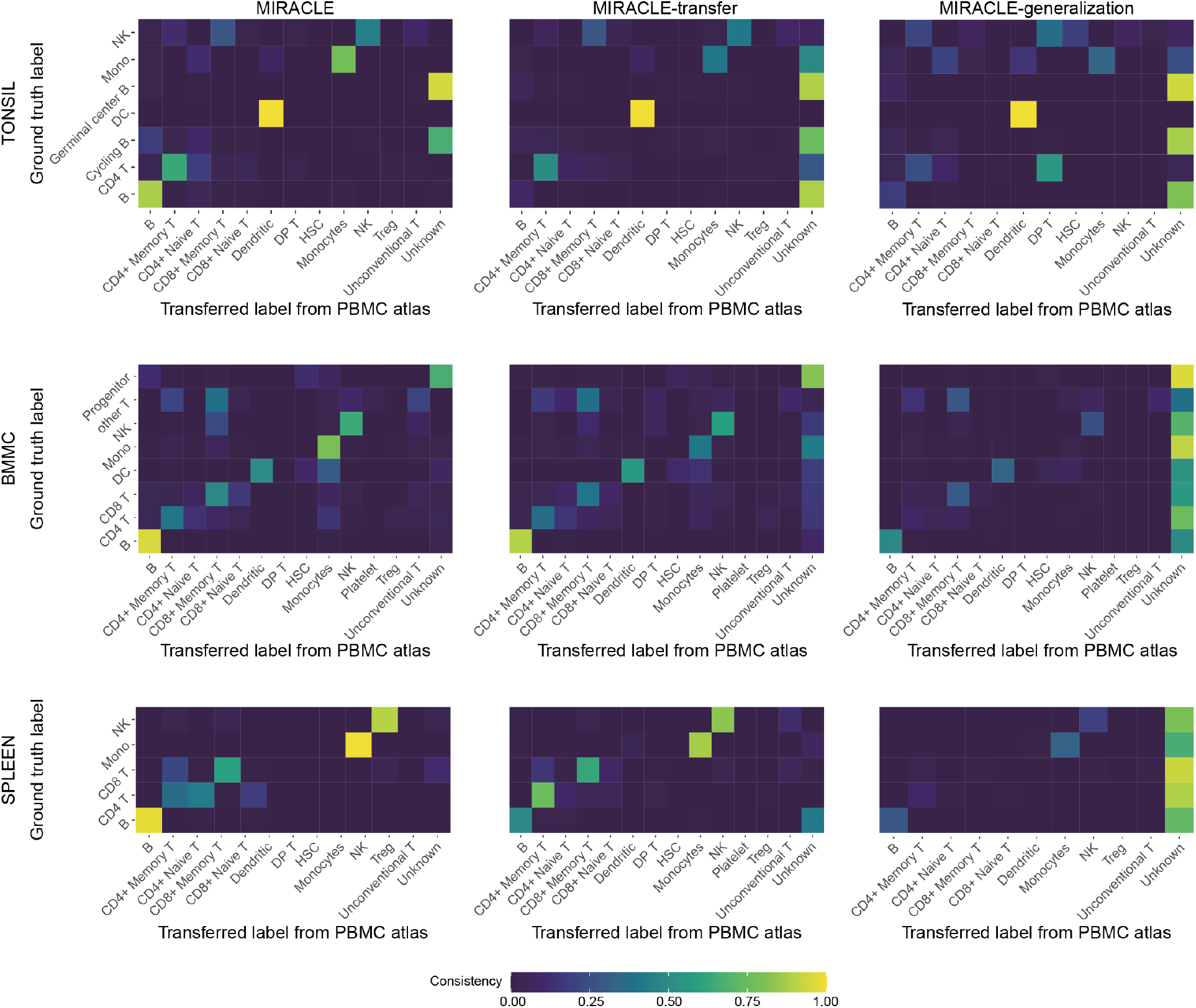
Confusion plots corresponding to the label transfer micro F1-scores for MIRACLE and two other strategies in cross-tissue label transfer tasks.

**Supplementary Table 1.** scIB benchmarking on the DHCM dataset.

| Method | Graph<br>iLISI | Graph<br>connectivity | kBET | Batch<br>correction | NMI | ARI | Isolated<br>label F1 | Graph<br>cLISI | Biological<br>conservation | Overall<br>score |
| --- | --- | --- | --- | --- | --- | --- | --- | --- | --- | --- |
| **MIRACLE-offline** | 0.3156 | 0.9718 | 0.4546 | 0.5807 | 0.9506 | 0.9677 | 0.8966 | 1.0000 | 0.9537 | 0.8045 |
| **MIRACLE-BTS-20K** | 0.2744 | 0.9559 | 0.4376 | 0.5560 | 0.9419 | 0.9628 | 0.8263 | 0.9998 | 0.9327 | 0.7820 |
| **PCA** | 0.1824 | 0.9757 | 0.3459 | 0.5013 | 0.9343 | 0.9229 | 0.9183 | 1.0000 | 0.9439 | 0.7669 |
| **MIRACLE-SRS-20K** | 0.2763 | 0.9591 | 0.4384 | 0.5579 | 0.9349 | 0.9604 | 0.0795 | 0.9997 | 0.7436 | 0.6693 |
| **MIRACLE-transfer** | 0.2330 | 0.9396 | 0.3860 | 0.5195 | 0.9295 | 0.9567 | 0.0414 | 0.9996 | 0.7318 | 0.6469 |
| **Geneformer** | 0.2348 | 0.9564 | 0.3690 | 0.5200 | 0.8938 | 0.8990 | 0.0786 | 0.9998 | 0.7178 | 0.6387 |
| **MIRACLE-520K** | 0.2723 | 0.9772 | 0.3160 | 0.5218 | 0.9516 | 0.9686 | 0.9191 | 1.0000 | 0.9598 | 0.7846 |
| **MIRACLE-BTS-200K** | 0.2918 | 0.9746 | 0.3673 | 0.5445 | 0.9537 | 0.9686 | 0.9132 | 1.0000 | 0.9589 | 0.7931 |
| **MIRACLE-BTS-100K** | 0.2852 | 0.9780 | 0.3754 | 0.5462 | 0.9529 | 0.9680 | 0.9098 | 1.0000 | 0.9577 | 0.7931 |
| **MIRACLE-BTS-50K** | 0.2669 | 0.9681 | 0.3732 | 0.5361 | 0.9449 | 0.9645 | 0.8931 | 1.0000 | 0.9506 | 0.7848 |
| **MIRACLE-BTS-20K** | 0.2744 | 0.9559 | 0.4376 | 0.5560 | 0.9419 | 0.9628 | 0.8263 | 0.9998 | 0.9327 | 0.7820 |
| **MIRACLE-BTS-10K** | 0.2812 | 0.9343 | 0.4915 | 0.5690 | 0.8952 | 0.9271 | 0.0518 | 0.9988 | 0.7182 | 0.6585 |
| **MIRACLE-BTS-5K** | 0.3000 | 0.8979 | 0.5958 | 0.5979 | 0.7849 | 0.8473 | 0.0430 | 0.9897 | 0.6662 | 0.6389 |
| **MIRACLE-BTS-2K** | 0.3007 | 0.8628 | 0.5257 | 0.5631 | 0.7518 | 0.8152 | 0.0159 | 0.9878 | 0.6427 | 0.6108 |

**Supplementary Table 2.** Construction of the cell type missing WNN dataset.

| Data batch |  | Cell type |  |  |  |  |  |  |  |
| --- | --- | --- | --- | --- | --- | --- | --- | --- | --- |
|  |  | B | CD4 T | CD8 T | DC | Mono | NK | other | other T |
| Original data | P1_0 | 236 | 2198 | 1154 | 128 | 1890 | 494 | 184 | 94 |
|  | P2_0 | 455 | 1921 | 1140 | 93 | 1284 | 554 | 75 | 377 |
|  | P3_0 | 363 | 1422 | 525 | 72 | 1159 | 890 | 40 | 157 |
|  | P4_0 | 410 | 1524 | 913 | 113 | 1740 | 119 | 46 | 420 |
|  | P5_0 | 543 | 896 | 1157 | 216 | 2747 | 863 | 190 | 340 |
|  | P6_0 | 729 | 1062 | 356 | 95 | 2755 | 795 | 151 | 117 |
|  | P7_0 | 987 | 1258 | 1223 | 383 | 3748 | 997 | 84 | 174 |
|  | P8_0 | 394 | 2052 | 1646 | 123 | 3188 | 1075 | 100 | 330 |
| Constructed data | P1_0 | 0 | 2198 | 1154 | 128 | 1890 | 494 | 184 | 94 |
|  | P2_0 | 455 | 0 | 1140 | 93 | 1284 | 554 | 75 | 377 |
|  | P3_0 | 363 | 1422 | 0 | 72 | 1159 | 890 | 40 | 157 |
|  | P4_0 | 410 | 1524 | 913 | 0 | 1740 | 119 | 46 | 420 |
|  | P5_0 | 543 | 896 | 1157 | 216 | 0 | 863 | 190 | 340 |
|  | P6_0 | 729 | 1062 | 356 | 95 | 2755 | 0 | 151 | 117 |
|  | P7_0 | 987 | 1258 | 1223 | 383 | 3748 | 997 | 0 | 174 |
|  | P8_0 | 394 | 2052 | 1646 | 123 | 3188 | 1075 | 100 | 0 |

\* Cell numbers of each batch belonging different cell types are presented.

**Supplementary Table 3.** scIB benchmarking of online methods on the WNN dataset.

| Method | Graph<br>iLISI | Graph<br>connectivity | kBET | Batch<br>correction | NMI | ARI | Isolated<br>label F1 | Graph<br>cLISI | Biological<br>conservation | Overall<br>score |
| --- | --- | --- | --- | --- | --- | --- | --- | --- | --- | --- |
| **MIRACLE** | 0.4485 | 0.9569 | 0.3138 | 0.5731 | 0.9079 | 0.9167 | 0.8958 | 1.0000 | 0.9301 | 0.7873 |
| **Multigrade+scArches** | 0.4248 | 0.9498 | 0.2776 | 0.5507 | 0.8248 | 0.8174 | 0.8118 | 0.9983 | 0.8631 | 0.7381 |
| **Symphony** | 0.4452 | 0.9525 | 0.2021 | 0.5333 | 0.8219 | 0.8108 | 0.8221 | 0.9996 | 0.8636 | 0.7315 |
| **scVAEIT-transfer** | 0.2887 | 0.9806 | 0.0493 | 0.4396 | 0.8676 | 0.8350 | 0.8957 | 1.0000 | 0.8996 | 0.7156 |
| **CCA+WNN+continual** | 0.5291 | 0.9483 | 0.1725 | 0.5500 | 0.7566 | 0.7074 | 0.7728 | 1.0000 | 0.8092 | 0.7055 |
| **RPCA+WNN+continual** | 0.3109 | 0.9389 | 0.1105 | 0.4534 | 0.8393 | 0.8255 | 0.8349 | 1.0000 | 0.8749 | 0.7063 |
| **MIRACLE-transfer** | 0.1543 | 0.9568 | 0.0182 | 0.3764 | 0.9039 | 0.9026 | 0.8803 | 1.0000 | 0.9217 | 0.7036 |
| **online iNMF** | 0.4983 | 0.9549 | 0.3790 | 0.6107 | 0.7017 | 0.6276 | 0.7066 | 0.9933 | 0.7573 | 0.6987 |
| **SCALEX-projection** | 0.3736 | 0.9513 | 0.0673 | 0.4641 | 0.8100 | 0.7839 | 0.8521 | 0.9975 | 0.8609 | 0.7022 |
| **sciPENN-transfer** | 0.3736 | 0.9155 | 0.0808 | 0.4566 | 0.7189 | 0.7621 | 0.7319 | 0.9975 | 0.8026 | 0.6642 |
| **Concerto-transfer** | 0.2980 | 0.9534 | 0.0694 | 0.4403 | 0.7246 | 0.6681 | 0.7755 | 0.9873 | 0.7889 | 0.6494 |
| **totalVI+scArches** | 0.0000 | 0.6722 | 0.0022 | 0.2248 | 0.4821 | 0.1710 | 0.3402 | 0.9918 | 0.4963 | 0.3877 |
| **trVAE+scArches** | 0.2949 | 0.7845 | 0.0576 | 0.3790 | 0.0939 | 0.0276 | 0.2643 | 0.6744 | 0.2651 | 0.3106 |

**Supplementary Table 4.** scIB benchmarking of offline methods on the WNN dataset.

| Method | Graph<br>iLISI | Graph<br>connectivity | kBET | Batch<br>correction | NMI | ARI | Isolated<br>label F1 | Graph<br>cLISI | Biological<br>conservation | Overall<br>score |
| --- | --- | --- | --- | --- | --- | --- | --- | --- | --- | --- |
| **MIRACLE-offline** | 0.4622 | 0.9570 | 0.3536 | 0.5909 | 0.8966 | 0.8856 | 0.9017 | 1.0000 | 0.9210 | 0.7890 |
| **Multigrade** | 0.4543 | 0.9518 | 0.3045 | 0.5702 | 0.8841 | 0.8803 | 0.8855 | 0.9995 | 0.9124 | 0.7755 |
| **scVAEIT** | 0.3907 | 0.9551 | 0.1523 | 0.4994 | 0.8757 | 0.8272 | 0.8498 | 1.0000 | 0.8882 | 0.7327 |
| **trVAE** | 0.5226 | 0.9489 | 0.5681 | 0.6799 | 0.7666 | 0.7442 | 0.8141 | 0.9413 | 0.8166 | 0.7619 |
| **online iNMF-offline** | 0.5073 | 0.9508 | 0.5131 | 0.6571 | 0.7538 | 0.7230 | 0.7759 | 0.9978 | 0.8126 | 0.7504 |
| **totalVI** | 0.3850 | 0.9570 | 0.1131 | 0.4850 | 0.9089 | 0.9190 | 0.9201 | 1.0000 | 0.9370 | 0.7562 |
| **CCA+WNN** | 0.5511 | 0.9515 | 0.2264 | 0.5763 | 0.7563 | 0.7093 | 0.7767 | 1.0000 | 0.8106 | 0.7169 |
| **Symphony-offline** | 0.4377 | 0.9496 | 0.1796 | 0.5223 | 0.7771 | 0.7617 | 0.8165 | 0.9982 | 0.8384 | 0.7119 |
| **RPCA+WNN** | 0.5136 | 0.9408 | 0.1688 | 0.5411 | 0.7812 | 0.7340 | 0.7920 | 1.0000 | 0.8268 | 0.7125 |
| **SCALEX** | 0.2749 | 0.9438 | 0.0309 | 0.4165 | 0.8384 | 0.8150 | 0.8874 | 0.9996 | 0.8851 | 0.6977 |
| **sciPENN** | 0.1732 | 0.9755 | 0.0129 | 0.3872 | 0.8091 | 0.7182 | 0.8316 | 1.0000 | 0.8397 | 0.6587 |
| **Concerto** | 0.3113 | 0.9753 | 0.1019 | 0.4628 | 0.7725 | 0.6772 | 0.7894 | 0.9945 | 0.8084 | 0.6702 |

**Supplementary Table 5.** Construction of the cell type missing mosaic DOTEA dataset.

| Data batch |  | Cell type |  |  |  |  |  |  |  | Modality |  |  |
| --- | --- | --- | --- | --- | --- | --- | --- | --- | --- | --- | --- | --- |
|  |  | B | CD4 T | CD8 T | DC | Mono | NK | other | other T | ATAC | RNA | ADT |
| Original data | DIG Ctrl | 463 | 6913 | 1713 | 57 | 112 | 298 | 23 | 611 | ✓ | ✓ | ✓ |
|  | DIG Stim | 332 | 6377 | 1608 | 62 | 64 | 431 | 33 | 620 | ✓ | ✓ | ✓ |
|  | LLL Ctrl | 296 | 4983 | 1435 | 43 | 53 | 166 | 21 | 364 | ✓ | ✓ | ✓ |
|  | LLL Stim | 159 | 3974 | 1080 | 45 | 11 | 213 | 27 | 388 | ✓ | ✓ | ✓ |
|  | W3 | 1245 | 2859 | 1066 | 78 | 216 | 309 | 652 | 162 | ✓ | ✓ | ✓ |
|  | W4 | 1390 | 2984 | 1088 | 39 | 206 | 312 | 716 | 162 | ✓ | ✓ | ✓ |
|  | W5 | 1280 | 2747 | 1283 | 64 | 275 | 343 | 711 | 207 | ✓ | ✓ | ✓ |
|  | W6 | 1326 | 3128 | 1119 | 13 | 277 | 328 | 742 | 204 | ✓ | ✓ | ✓ |
| Constructed data | DIG Ctrl | 463 | 6913 | 1713 | 57 | 112 | 298 | 23 | 0 | ✓ |  | ✓ |
|  | DIG Stim | 332 | 6377 | 1608 | 62 | 64 | 0 | 33 | 620 | ✓ | ✓ |  |
|  | LLL Ctrl | 296 | 4983 | 1435 | 0 | 53 | 166 | 21 | 364 |  | ✓ | ✓ |
|  | LLL Stim | 159 | 3974 | 0 | 45 | 11 | 213 | 27 | 388 | ✓ | ✓ | ✓ |
|  | W3 | 0 | 2859 | 1066 | 78 | 216 | 309 | 652 | 162 | ✓ |  | ✓ |
|  | W4 | 1390 | 2984 | 1088 | 39 | 206 | 312 | 0 | 162 | ✓ | ✓ |  |
|  | W5 | 1280 | 0 | 1283 | 64 | 275 | 343 | 711 | 207 |  | ✓ | ✓ |
|  | W6 | 1326 | 3128 | 1119 | 13 | 0 | 328 | 742 | 204 | ✓ | ✓ | ✓ |

\* Cell numbers across different cell types and modalities of each batch are presented.
\* To address the large difference in the number of various cell types in the DOTEa dataset, we adopted the strategy that minimizing the variance of the number of deleted cells.

**Supplementary Table 6.** scMIB benchmarking of online and offline methods on the DOTEA dataset.

| Method | Graph<br>iLISI | Graph<br>connect<br>ivity | kBET | Batch<br>correct<br>ion | Modali<br>ty<br>ASW | FOSC<br>TTM | Label<br>Transf<br>er F1 | ATAC<br>AURO<br>C | RNA<br>Pearso<br>n's R | ADT<br>Pearso<br>n's R | Modali<br>ty<br>alignm<br>ent | NMI | ARI | Isolate<br>d label<br>F1 | Graph<br>cLISI | Biologi<br>cal<br>conserv<br>ation | Overall<br>score |
| --- | --- | --- | --- | --- | --- | --- | --- | --- | --- | --- | --- | --- | --- | --- | --- | --- | --- |
| **MRACLE** | 0.4819 | 0.9389 | 0.3649 | 0.5952 | 0.9040 | 0.9030 | 0.8413 | 0.8171 | 0.7209 | 0.8539 | 0.8400 | 0.6433 | 0.7028 | 0.7591 | 0.9834 | 0.7721 | 0.7394 |
| **MIRACLE-**<br>**offline** | 0.4929 | 0.9390 | 0.3962 | 0.6093 | 0.9291 | 0.9371 | 0.8722 | 0.8178 | 0.8201 | 0.8928 | 0.8782 | 0.6031 | 0.6668 | 0.6675 | 0.9788 | 0.7291 | 0.7379 |
| **scVAEIT** | 0.1345 | 0.9114 | 0.0163 | 0.3541 | 0.8052 | 0.9094 | 0.8959 | 0.8195 | 0.8523 | 0.8661 | 0.8581 | 0.6581 | 0.7202 | 0.7135 | 0.9900 | 0.7705 | 0.6718 |
| **MIRACLE-**<br>**transfer** | 0.1258 | 0.9381 | 0.0464 | 0.3701 | 0.8333 | 0.8347 | 0.8126 | 0.8070 | 0.5865 | 0.5838 | 0.7430 | 0.5972 | 0.4305 | 0.6937 | 0.9886 | 0.6775 | 0.6049 |
| **Multigrade** | 0.2226 | 0.9253 | 0.0301 | 0.3927 | 0.9920 | 0.9873 | 0.9264 | 0.6303 | 0.6687 | 0.7860 | 0.8318 | 0.5290 | 0.3830 | 0.6458 | 0.9849 | 0.6357 | 0.6216 |
| **Multigrade+s**<br>**cArches** | 0.1243 | 0.9135 | 0.0050 | 0.3476 | 0.9916 | 0.9706 | 0.9165 | 0.5263 | 0.7684 | 0.8349 | 0.8347 | 0.3986 | 0.2332 | 0.5810 | 0.9665 | 0.5448 | 0.5726 |
| **scVAEIT+tr**<br>**ansfer** | 0.1237 | 0.8938 | 0.0486 | 0.3554 | 0.5021 | 0.6727 | 0.6837 | 0.8032 | 0.7355 | 0.4310 | 0.6380 | 0.4058 | 0.1862 | 0.5638 | 0.9619 | 0.5294 | 0.5098 |

**Supplementary Table 7.** Datasets used in atlas construction and knowledge transfer.

| Dataset | Batch | Modality |  |  | Cell number | Knowledge transfer |  | Knowledge transfer |  |
| --- | --- | --- | --- | --- | --- | --- | --- | --- | --- |
|  |  | ATAC | RNA | ADT |  | Referenc<br>e (atlas-no-neap,<br>PBMC) | Query<br>(neap,<br>PBMC) | Referenc<br>e (atlas,<br>PBMC) | Query<br>(tonsil, bm,<br>spleen) |
| TONSIL | TONSIL |  | ✓ |  | 5,503 |  |  |  | ✓ |
| BMMC dataset | BMMC | ✓ |  | ✓ | 10,671 |  |  |  | ✓ |
| SPLEEN dataset | SPLEEN |  | ✓ |  | 4,287 |  |  |  | ✓ |
| NEAT dataset | lane1 | ✓ | ✓ |  | 5,568 |  | ✓ | ✓ |  |
|  | lane2 | ✓ | ✓ |  | 4,956 |  | ✓ | ✓ |  |
| ASAP dataset | Ctrl | ✓ |  | ✓ | 4,255 |  | ✓ | ✓ |  |
|  | Stim | ✓ |  | ✓ | 5,241 |  | ✓ | ✓ |  |
| ASAP CITE dataset | Ctrl |  | ✓ | ✓ | 5,086 |  | ✓ | ✓ |  |
|  | Stim |  | ✓ | ✓ | 3,629 |  | ✓ | ✓ |  |
| DOGMA dataset | LLL Ctrl | ✓ | ✓ | ✓ | 7,361 | ✓ |  | ✓ |  |
|  | LLL Stim | ✓ | ✓ | ✓ | 5,897 | ✓ |  | ✓ |  |
|  | DIG Ctrl | ✓ | ✓ | ✓ | 10,190 | ✓ |  | ✓ |  |
|  | DIG Stim | ✓ | ✓ | ✓ | 9,527 | ✓ |  | ✓ |  |
| TEA dataset | W1 | ✓ | ✓ | ✓ | 7,325 | ✓ |  | ✓ |  |
|  | W3 | ✓ | ✓ | ✓ | 6,587 | ✓ |  | ✓ |  |
|  | W4 | ✓ | ✓ | ✓ | 6,897 | ✓ |  | ✓ |  |
|  | W5 | ✓ | ✓ | ✓ | 6,910 | ✓ |  | ✓ |  |
|  | W6 | ✓ | ✓ | ✓ | 7,137 | ✓ |  | ✓ |  |
| TEA Multiome dataset | W1 | ✓ | ✓ |  | 6,096 | ✓ |  | ✓ |  |
|  | W2 | ✓ | ✓ |  | 7,284 | ✓ |  | ✓ |  |
| 10X Multiome dataset | ChrX | ✓ | ✓ |  | 9,868 | ✓ |  | ✓ |  |
|  | ChrC | ✓ | ✓ |  | 9,582 | ✓ |  | ✓ |  |
|  | ARC2 10K | ✓ | ✓ |  | 11,116 | ✓ |  | ✓ |  |
|  | ARC2 3K | ✓ | ✓ |  | 2,566 | ✓ |  | ✓ |  |
| WNN CITE dataset | P1_0 |  | ✓ | ✓ | 6,378 | ✓ |  | ✓ |  |
|  | P2_0 |  | ✓ | ✓ | 5,899 | ✓ |  | ✓ |  |
|  | P3_0 |  | ✓ | ✓ | 4,628 | ✓ |  | ✓ |  |
|  | P4_0 |  | ✓ | ✓ | 5,285 | ✓ |  | ✓ |  |
|  | P5_0 |  | ✓ | ✓ | 6,952 | ✓ |  | ✓ |  |
|  | P6_0 |  | ✓ | ✓ | 6,060 | ✓ |  | ✓ |  |
|  | P7_0 |  | ✓ | ✓ | 8,854 | ✓ |  | ✓ |  |
|  | P8_0 |  | ✓ | ✓ | 8,908 | ✓ |  | ✓ |  |
| DOGMA-CITE dataset | CITE |  | ✓ | ✓ | 11,967 | ✓ |  | ✓ |  |
|  | LLL | ✓ | ✓ | ✓ | 9,794 | ✓ |  | ✓ |  |
|  | DIG | ✓ | ✓ | ✓ | 14,838 | ✓ |  | ✓ |  |
|  | DOGMA | ✓ | ✓ | ✓ | 13,095 | ✓ |  | ✓ |  |
| ISSAAC dataset | ISSAAC | ✓ | ✓ |  | 7,053 | ✓ |  | ✓ |  |
| Total number of cells |  | 189,814 | 253,083 | 199,371 | 273,250 | 224,054 | 28,735 | 252,789 | 20,461 |

**Supplementary Table 8.**
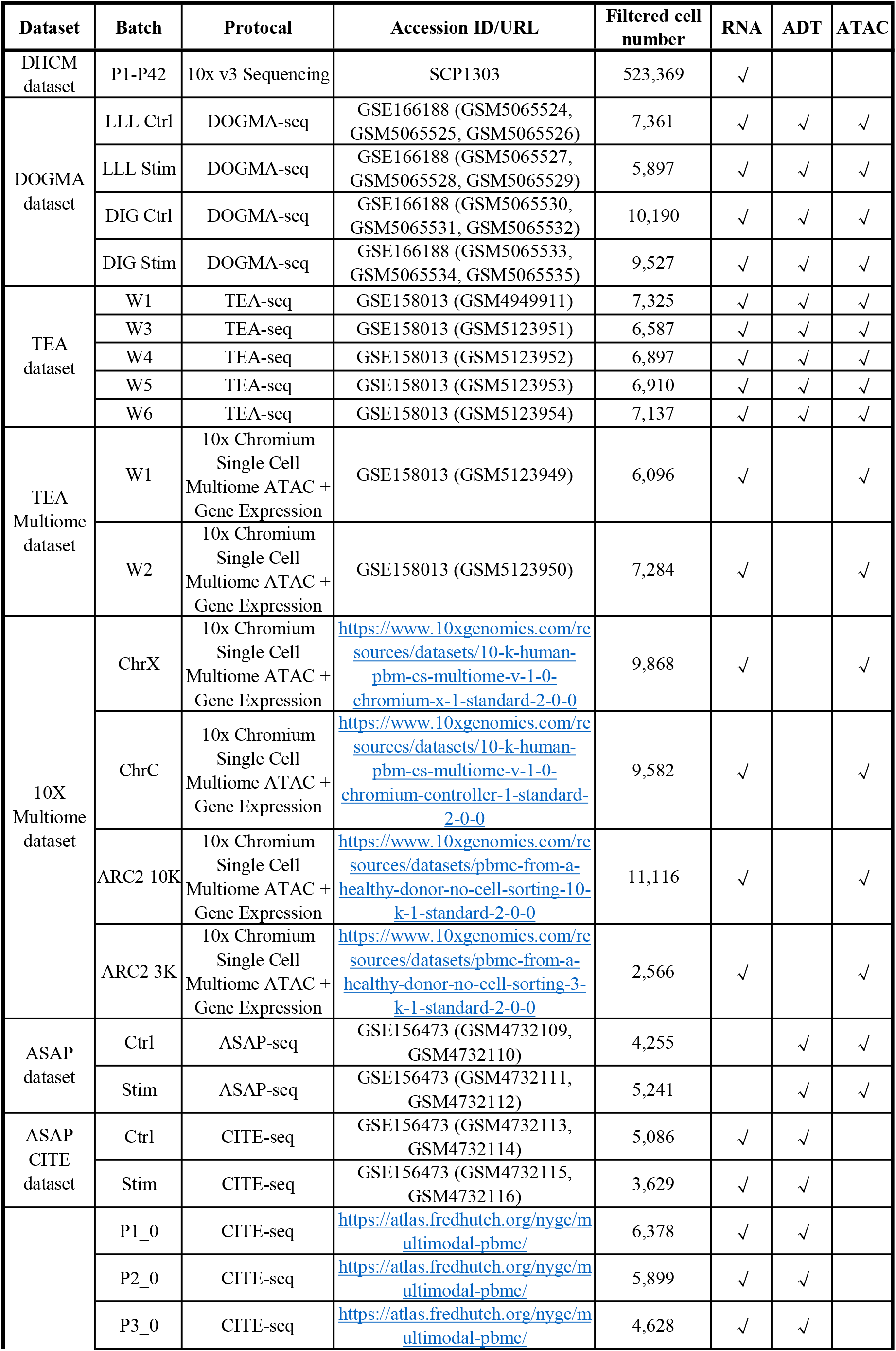
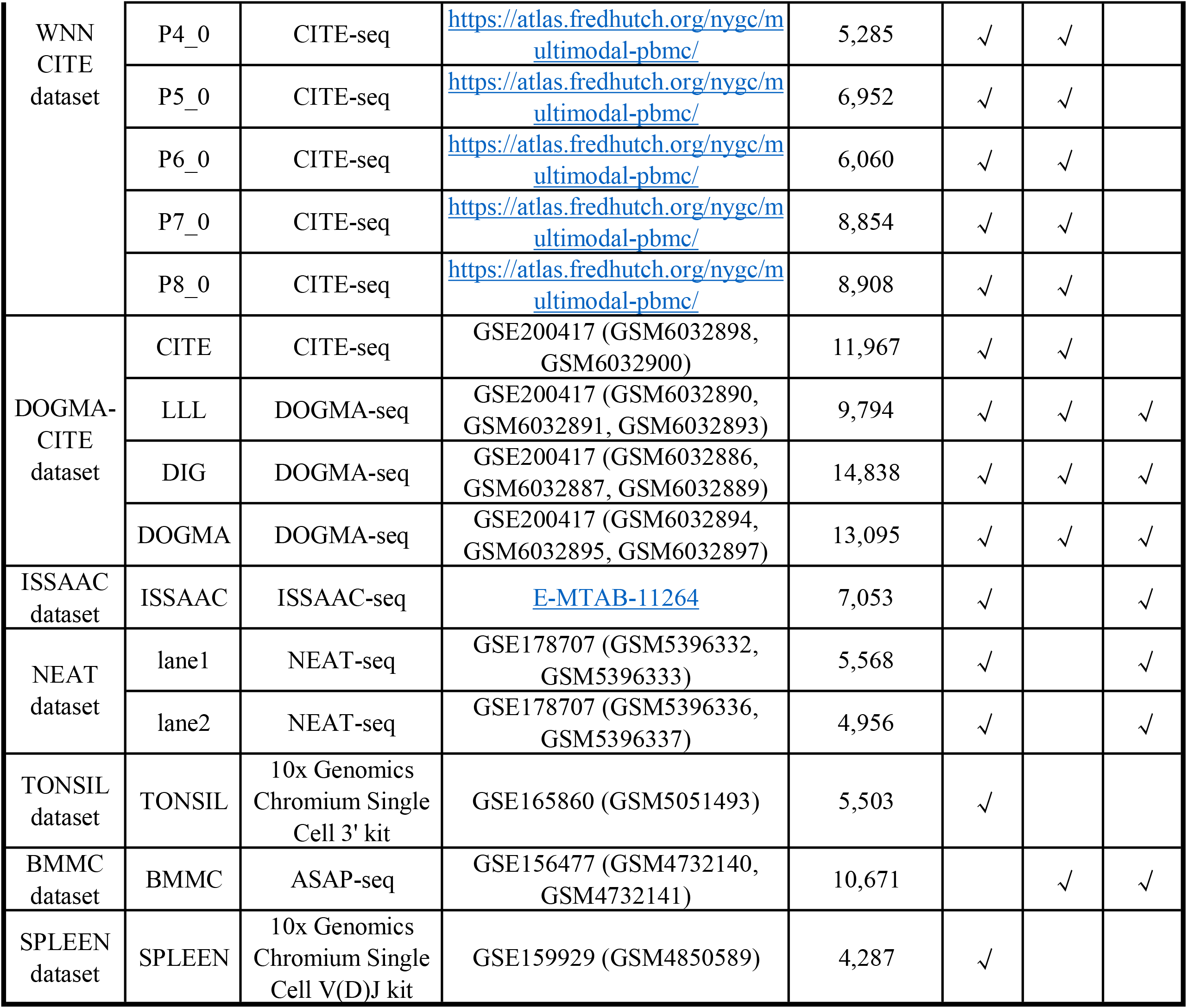
Public datasets used in this study.

**Supplementary Table 9.** Overview of compared methods for single-cell multimodal integration.

| Integration type | Method | Scenario |  |  |  | Task |  |  |
| --- | --- | --- | --- | --- | --- | --- | --- | --- |
|  |  | offline | online |  |  | Unimodal task<br>(RNA,<br>horizontal) | Bimodal task<br>(RNA+ADT,<br>rectangular) | Trimodal task<br>(ATAC+RNA+ADT,<br>mosaic) |
|  |  |  | generalization | fine-tuning | continual |  |  |  |
| Mosaic | MIRACLE-offline | √ |  |  |  | √ | √ | √ |
|  | MIRACLE |  |  |  | √ | √ | √ | √ |
|  | MIRACLE-transfer |  |  | √ |  | √ | √ | √ |
|  | Multigrade | √ |  |  |  |  | √ | √ |
|  | Multigrade+scArches |  | √ |  |  |  | √ | √ |
|  | scVAEIT | √ |  |  |  |  | √ | √ |
|  | scVAEIT-transfer* |  |  | √ |  |  | √ | √ |
|  | totalVI | √ |  |  |  |  | √ |  |
|  | totalVI+scArches |  | √ |  |  |  | √ |  |
| Rectangular | SCALEX | √ |  |  |  |  | √ |  |
|  | SCALEX-projection |  | √ |  |  |  | √ |  |
|  | trVAE | √ |  |  |  |  | √ |  |
|  | trVAE+scArches |  | √ |  |  |  | √ |  |
|  | Concerto | √ |  |  |  |  | √ |  |
|  | Concerto-transfer |  |  | √ |  |  | √ |  |
|  | Symphony-offline | √ |  |  |  |  | √ |  |
|  | Symphony |  | √ |  |  |  | √ |  |
|  | online iNMF-offline | √ |  |  |  |  | √ |  |
|  | online iNMF |  |  | √ |  |  | √ |  |
|  | CCA+WNN | √ |  |  |  |  | √ |  |
|  | CCA+WNN-continual |  |  |  | √ |  | √ |  |
|  | RPCA+WNN | √ |  |  |  |  | √ |  |
|  | RPCA+WNN-continual |  |  |  | √ |  | √ |  |
|  | sciPENN | √ |  |  |  |  | √ |  |
|  | sciPENN-transfer* |  |  | √ |  |  | √ |  |
| Horizontal | Geneformer |  | √ |  |  | √ |  |  |
|  | PCA | √ |  |  |  | √ |  |  |

The asterisk (\*) indicates that this method does not support online scenario, thus we adopted fine-tuning-based transfer learning strategies.

## Notes

### Competing Interest Statement

The authors have declared no competing interest.

### Summary of Updates

Some metadata of the paper has been corrected.

## References

[1] Tim Stuart and Rahul Satija. Integrative single-cell analysis. Nature Reviews Genetics, 20(5):257–272, May 2019. ISSN 1471-0064. doi: 10.1038/s41576-019-0093-7.

[2] Katy Vandereyken, Alejandro Sifrim, Bernard Thienpont, and Thierry Voet. Methods and applications for single-cell and spatial multi-omics. Nature Reviews Genetics, pages 1–22, March 2023. ISSN 1471-0064. doi: 10.1038/s41576-023-00580-2.

[3] Alev Baysoy, Zhiliang Bai, Rahul Satija, and Rong Fan. The technological landscape and applications of single-cell multi-omics. Nature Reviews Molecular Cell Biology, pages 1–19, June 2023. ISSN 1471-0080. doi: 10.1038/s41580-023-00615-w.

[4] Lisa Sikkema, Ciro Ramírez-Suástegui, Daniel C. Strobl, Tessa E. Gillett, Luke Zappia, Elo Madissoon, Nikolay S. Markov, Laure-Emmanuelle Zaragosi, Yuge Ji, Meshal Ansari, Marie-Jeanne Arguel, Leonie Apperloo, Martin Banchero, Christophe Bécavin, Marijn Berg, Evgeny Chichelnitskiy, Mei-i Chung, Antoine Collin, Aurore C. A. Gay, Janine Gote-Schniering, Baharak Hooshiar Kashani, Kemal Inecik, Manu Jain, Theodore S. Kapellos, Tessa M. Kole, Sylvie Leroy, Christoph H. Mayr, Amanda J. Oliver, Michael von Papen, Lance Peter, Chase J. Taylor, Thomas Walzthoeni, Chuan Xu, Linh T. Bui, Carlo De Donno, Leander Dony, Alen Faiz, Minzhe Guo, Austin J. Gutierrez, Lukas Heumos, Ni Huang, Ignacio L. Ibarra, Nathan D. Jackson, Preetish Kadur Lakshminarasimha Murthy, Mohammad Lotfollahi, Tracy Tabib, Carlos Talavera-López, Kyle J. Travaglini, Anna Wilbrey-Clark, Kaylee B. Worlock, Masahiro Yoshida, Maarten van den Berge, Yohan Bossé, Tushar J. Desai, Oliver Eickelberg, Naftali Kaminski, Mark A. Krasnow, Robert Lafyatis, Marko Z. Nikolic, Joseph E. Powell, Jayaraj Rajagopal, Mauricio Rojas, Orit Rozenblatt-Rosen, Max A. Seibold, Dean Sheppard, Douglas P. Shepherd, Don D. Sin, Wim Timens, Alexander M. Tsankov, Jeffirey Whitsett, Yan Xu, Nicholas E. Banovich, Pascal Barbry, Thu Elizabeth Duong, Christine S. Falk, Kerstin B. Meyer, Jonathan A. Kropski, Dana Pe’er, Herbert B. Schiller, Purushothama Rao Tata, Joachim L. Schultze, Sara A. Teichmann, Alexander V. Misharin, Martijn C. Nawijn, Malte D. Luecken, and Fabian J. Theis. An integrated cell atlas of the lung in health and disease. Nature Medicine, 29(6):1563–1577, June 2023. ISSN 1546-170X. doi: 10.1038/s41591-023-02327-2.

[5] Xinyue Chen, Yin Huang, Liangfeng Huang, Ziliang Huang, Zhao-Zhe Hao, Lahong Xu, Nana Xu, Zhi Li, Yonggao Mou, Mingli Ye, Renke You, Xuegong Zhang, Sheng Liu, and Zhichao Miao. A brain cell atlas integrating single-cell transcriptomes across human brain regions. Nature Medicine, pages 1–13, August 2024. ISSN 1546-170X. doi: 10.1038/s41591-024-03150-z.

[6] Tim Stuart, Andrew Butler, Paul Hoffman, Christoph Hafemeister, Efthymia Papalexi, William M. Mauck, Yuhan Hao, Marlon Stoeckius, Peter Smibert, and Rahul Satija. Comprehensive Integration of Single-Cell Data. Cell, 177(7):1888–1902.e21, June 2019. ISSN 0092-8674. doi: 10.1016/j.cell.2019.05.031.

[7] Adam Gayoso, Zoë Steier, Romain Lopez, Jefirey Regier, Kristopher L. Nazor, Aaron Streets, and Nir Yosef. Joint probabilistic modeling of single-cell multi-omic data with totalVI. Nature Methods, pages 1–11, February 2021. ISSN 1548-7105. doi: 10.1038/s41592-020-01050-x.

[8] Zhi-Jie Cao and Ge Gao. Multi-omics single-cell data integration and regulatory inference with graph-linked embedding. Nature Biotechnology, pages 1–9, May 2022. ISSN 1546-1696. doi: 10.1038/s41587-022-01284-4.

[9] Tal Ashuach, Mariano I. Gabitto, Rohan V. Koodli, Giuseppe-Antonio Saldi, Michael I. Jordan, and Nir Yosef. MultiVI: Deep generative model for the integration of multimodal data. Nature Methods, pages 1–10, June 2023. ISSN 1548-7105. doi: 10.1038/s41592-023-01909-9.

[10] Chao Gao, Jialin Liu, April R. Kriebel, Sebastian Preissl, Chongyuan Luo, Rosa Castanon, Justin Sandoval, Angeline Rivkin, Joseph R. Nery, Margarita M. Behrens, Joseph R. Ecker, Bing Ren, and Joshua D. Welch. Iterative single-cell multi-omic integration using online learning. Nature Biotechnology, pages 1–8, April 2021. ISSN 1546-1696. doi: 10.1038/s41587-021-00867-x.

[11] Mohammad Lotfollahi, Mohsen Naghipourfar, Malte D. Luecken, Matin Khajavi, Maren Büttner, Marco Wagenstetter, Žiga Avsec, Adam Gayoso, Nir Yosef, Marta Interlandi, Sergei Rybakov, Alexander V. Misharin, and Fabian J. Theis. Mapping single-cell data to reference atlases by transfer learning. Nature Biotechnology, 40(1):121–130, January 2022. ISSN 1546-1696. doi: 10.1038/s41587-021-01001-7.

[12] Meng Yang, Yueyuxiao Yang, Chenxi Xie, Ming Ni, Jian Liu, Huanming Yang, Feng Mu, and Jian Wang. Contrastive learning enables rapid mapping to multimodal single-cell atlas of multimillion scale. Nature Machine Intelligence, pages 1–14, August 2022. ISSN 2522-5839. doi: 10.1038/s42256-022-00518-z.

[13] Lei Xiong, Kang Tian, Yuzhe Li, Weixi Ning, Xin Gao, and Qiangfeng Cliff Zhang. Online single-cell data integration through projecting heterogeneous datasets into a common cell-embedding space. Nature Communications, 13(1):6118, October 2022. ISSN 2041-1723. doi: 10.1038/s41467-022-33758-z.

[14] Joyce B. Kang, Aparna Nathan, Kathryn Weinand, Fan Zhang, Nghia Millard, Laurie Rumker, D. Branch Moody, Ilya Korsunsky, and Soumya Raychaudhuri. Efficient and precise single-cell reference atlas mapping with Symphony. Nature Communications, 12(1):5890, October 2021. ISSN 2041-1723. doi: 10.1038/s41467-021-25957-x.

[15] Fan Yang, Wenchuan Wang, Fang Wang, Yuan Fang, Duyu Tang, Junzhou Huang, Hui Lu, and Jianhua Yao. scBERT as a large-scale pretrained deep language model for cell type annotation of single-cell RNA-seq data. Nature Machine Intelligence, pages 1–15, September 2022. ISSN 2522-5839. doi: 10.1038/s42256-022-00534-z.

[16] Ricard Argelaguet, Anna S. E. Cuomo, Oliver Stegle, and John C. Marioni. Computational principles and challenges in single-cell data integration. Nature Biotechnology, pages 1–14, May 2021. ISSN 1546-1696. doi: 10.1038/s41587-021-00895-7.

[17] Gido M. van de Ven, Tinne Tuytelaars, and Andreas S. Tolias. Three types of incremental learning. Nature Machine Intelligence, pages 1–13, December 2022. ISSN 2522-5839. doi: 10.1038/s42256-022-00568-3.

[18] Liyuan Wang, Xingxing Zhang, Hang Su, and Jun Zhu. A Comprehensive Survey of Continual Learning: Theory, Method and Application. IEEE Transactions on Pattern Analysis and Machine Intelligence, pages 1–20, 2024. ISSN 1939-3539. doi: 10.1109/TPAMI.2024.3367329.

[19] Zhen He, Shuofeng Hu, Yaowen Chen, Sijing An, Jiahao Zhou, Runyan Liu, Junfeng Shi, Jing Wang, Guohua Dong, Jinhui Shi, Jiaxin Zhao, L. Ou-Yang, Yuan Zhu, Xiaochen Bo, and Xiaomin Ying. Mosaic integration and knowledge transfer of single-cell multimodal data with MIDAS. Nature Biotechnology, pages 1–12, January 2024. ISSN 1546-1696. doi: 10.1038/s41587-023-02040-y.

[20] Romain Lopez, Adam Gayoso, and Nir Yosef. Enhancing scientific discoveries in molecular biology with deep generative models. Molecular Systems Biology, 16(9):e9198, September 2020. ISSN 1744-4292. doi: 10.15252/msb.20199198.

[21] Sam Bond-Taylor, Adam Leach, Yang Long, and Chris G. Willcocks. Deep Generative Modelling: A Comparative Review of VAEs, GANs, Normalizing Flows, Energy-Based and Autoregressive Models. IEEE Transactions on Pattern Analysis and Machine Intelligence, pages 1–1, 2021. ISSN 1939-3539. doi: 10.1109/TPAMI.2021.3116668.

[22] Diederik P. Kingma and Max Welling. An Introduction to Variational Autoencoders. Foundations and Trends® in Machine Learning, 12(4):307–392, November 2019. ISSN 1935-8237, 1935-8245. doi: 10.1561/2200000056.

[23] Mark Chaffin, Irinna Papangeli, Bridget Simonson, Amer-Denis Akkad, Matthew C. Hill, Alessandro Arduini, Stephen J. Fleming, Michelle Melanson, Sikander Hayat, Maria Kost-Alimova, Ondine Atwa, Jiangchuan Ye, Kenneth C. Bedi, Matthias Nahrendorf, Virendar K. Kaushik, Christian M. Stegmann, Kenneth B. Margulies, Nathan R. Tucker, and Patrick T. Ellinor. Single-nucleus profiling of human dilated and hypertrophic cardiomyopathy. Nature, 608(7921):174–180, August 2022. ISSN 1476-4687. doi: 10.1038/s41586-022-04817-8.

[24] Yuhan Hao, Tim Stuart, Madeline H. Kowalski, Saket Choudhary, Paul Hoffman, Austin Hartman, Avi Srivastava, Gesmira Molla, Shaista Madad, Carlos Fernandez-Granda, and Rahul Satija. Dictionary learning for integrative, multimodal and scalable single-cell analysis. Nature Biotechnology, pages 1–12, May 2023. ISSN 1546-1696. doi: 10.1038/s41587-023-01767-y.

[25] Dongyuan Song, Nan Miles Xi, Jingyi Jessica Li, and Lin Wang. scSampler: Fast diversity-preserving subsampling of large-scale single-cell transcriptomic data. Bioinformatics, 38(11):3126–3127, May 2022. ISSN 1367-4803. doi: 10.1093/bioinformatics/btac271.

[26] Malte D. Luecken, M. Büttner, K. Chaichoompu, A. Danese, M. Interlandi, M. F. Mueller, D. C. Strobl, L. Zappia, M. Dugas, M. Colomé-Tatché, and Fabian J. Theis. Benchmarking atlas-level data integration in single-cell genomics. Nature Methods, 19(1):41–50, January 2022. ISSN 1548-7105. doi: 10.1038/s41592-021-01336-8.

[27] Christina V. Theodoris, Ling Xiao, Anant Chopra, Mark D. Chaffin, Zeina R. Al Sayed, Matthew C. Hill, Helene Mantineo, Elizabeth M. Brydon, Zexian Zeng, X. Shirley Liu, and Patrick T. Ellinor. Transfer learning enables predictions in network biology. Nature, 618(7965):616–624, June 2023. ISSN 1476-4687. doi: 10.1038/s41586-023-06139-9.

[28] Yuhan Hao, Stephanie Hao, Erica Andersen-Nissen, William M. Mauck, Shiwei Zheng, Andrew Butler, Maddie J. Lee, Aaron J. Wilk, Charlotte Darby, Michael Zager, Paul Hoffman, Marlon Stoeckius, Efthymia Papalexi, Eleni P. Mimitou, Jaison Jain, Avi Srivastava, Tim Stuart, Lamar M. Fleming, Bertrand Yeung, Angela J. Rogers, Juliana M. McElrath, Catherine A. Blish, Raphael Gottardo, Peter Smibert, and Rahul Satija. Integrated analysis of multimodal single-cell data. Cell, May 2021. ISSN 0092-8674. doi: 10.1016/j.cell.2021.04.048.

[29] Eleni P. Mimitou, Caleb A. Lareau, Kelvin Y. Chen, Andre L. Zorzetto-Fernandes, Yuhan Hao, Yusuke Takeshima, Wendy Luo, Tse-Shun Huang, Bertrand Z. Yeung, Efthymia Papalexi, Pratiksha I. Thakore, Tatsuya Kibayashi, James Badger Wing, Mayu Hata, Rahul Satija, Kristopher L. Nazor, Shimon Sakaguchi, Leif S. Ludwig, Vijay G. Sankaran, Aviv Regev, and Peter Smibert. Scalable, multimodal profiling of chromatin accessibility, gene expression and protein levels in single cells. Nature Biotechnology, pages 1–13, June 2021. ISSN 1546-1696. doi: 10.1038/s41587-021-00927-2.

[30] Elliott Swanson, Cara Lord, Julian Reading, Alexander T Heubeck, Palak C Genge, Zachary Thomson, Morgan DA Weiss, Xiao-jun Li, Adam K Savage, Richard R Green, Troy R Torgerson, Thomas F Bumol, Lucas T Graybuck, and Peter J Skene. Simultaneous trimodal single-cell measurement of transcripts, epitopes, and chromatin accessibility using TEA-seq. eLife, 10:e63632, April 2021. ISSN 2050-084X. doi: 10.7554/eLife.63632.

[31] Amy F. Chen, Benjamin Parks, Arwa S. Kathiria, Benjamin Ober-Reynolds, Jorg J. Goronzy, and William J. Greenleaf. NEAT-seq: Simultaneous profiling of intra-nuclear proteins, chromatin accessibility and gene expression in single cells. Nature Methods, 19(5):547–553, May 2022. ISSN 1548-7105. doi: 10.1038/s41592-022-01461-y.

[32] Hamish W. King, Kristen L. Wells, Zohar Shipony, Arwa S. Kathiria, Lisa E. Wagar, Caleb Lareau, Nara Orban, Robson Capasso, Mark M. Davis, Lars M. Steinmetz, Louisa K. James, and William J. Greenleaf. Integrated single-cell transcriptomics and epigenomics reveals strong germinal center–associated etiology of autoimmune risk loci. Science Immunology, 6(64):eabh3768, October 2021. doi: 10.1126/sciimmunol.abh3768.

[33] Shuai He, Lin-He Wang, Yang Liu, Yi-Qi Li, Hai-Tian Chen, Jing-Hong Xu, Wan Peng, Guo-Wang Lin, Pan-Pan Wei, Bo Li, Xiaojun Xia, Dan Wang, Jin-Xin Bei, Xiaoshun He, and Zhiyong Guo. Single-cell transcriptome profiling of an adult human cell atlas of 15 major organs. Genome Biology, 21(1):294, December 2020. ISSN 1474-760X. doi: 10.1186/s13059-020-02210-0.

[34] Justin Lakkis, Amelia Schroeder, Kenong Su, Michelle Y. Y. Lee, Alexander C. Bashore, Muredach P. Reilly, and Mingyao Li. A multi-use deep learning method for CITE-seq and single-cell RNA-seq data integration with cell surface protein prediction and imputation. Nature Machine Intelligence, pages 1–13, October 2022. ISSN 2522-5839. doi: 10.1038/s42256-022-00545-w.

[35] Jin-Hong Du, Zhanrui Cai, and Kathryn Roeder. Robust probabilistic modeling for single-cell multimodal mosaic integration and imputation via scVAEIT. Proceedings of the National Academy of Sciences, 119(49):e2214414119, December 2022. doi: 10.1073/pnas.2214414119.

[36] Abhishek Kumar, Sunabha Chatterjee, and Piyush Rai. Bayesian Structural Adaptation for Continual Learning. In Proceedings of the 38th International Conference on Machine Learning, pages 5850–5860. PMLR, July 2021.

[37] Qiang Gao, Zhipeng Luo, Diego Klabjan, and Fengli Zhang. Efficient Architecture Search for Continual Learning. IEEE Transactions on Neural Networks and Learning Systems, 34(11):8555–8565, November 2023. ISSN 2162-2388. doi: 10.1109/TNNLS.2022.3151511.

[38] Chen Zhang, Yu Xie, Hang Bai, Bin Yu, Weihong Li, and Yuan Gao. A survey on federated learning. Knowledge-Based Systems, 216:106775, March 2021. ISSN 0950-7051. doi: 10.1016/j.knosys.2021.106775.

[39] Andrea Soltoggio, Eseoghene Ben-Iwhiwhu, Vladimir Braverman, Eric Eaton, Benjamin Epstein, Yunhao Ge, Lucy Halperin, Jonathan How, Laurent Itti, Michael A. Jacobs, Pavan Kantharaju, Long Le, Steven Lee, Xinran Liu, Sildomar T. Monteiro, David Musliner, Saptarshi Nath, Priyadarshini Panda, Christos Peridis, Hamed Pirsiavash, Vishwa Parekh, Kaushik Roy, Shahaf Shperberg, Hava T. Siegelmann, Peter Stone, Kyle Vedder, Jingfeng Wu, Lin Yang, Guangyao Zheng, and Soheil Kolouri. A collective AI via lifelong learning and sharing at the edge. Nature Machine Intelligence, 6(3):251–264, March 2024. ISSN 2522-5839. doi: 10.1038/s42256-024-00800-2.

[40] William Gemmell Cochran. Sampling Techniques. Wiley Series in Probability and Mathematical Statistics. Wiley, New York, 3d ed edition, 1977. ISBN 978-0-471-16240-7.

[41] Stephen M Omohundro. Five Balltree Construction Algorithms. International Computer Science Institute Berkeley, 1989.

[42] Ron Edgar, Michael Domrachev, and Alex E. Lash. Gene Expression Omnibus: NCBI gene expression and hybridization array data repository. Nucleic Acids Research, 30(1):207–210, January 2002. ISSN 0305-1048. doi: 10.1093/nar/30.1.207.

[43] 10k Human PBMCs, Multiome v1.0, Chromium X. https://www.10xgenomics.com/resources/datasets/10-khuman-pbm-cs-multiome-v-1-0-chromium-x-1-standard-2-0-0, 2021.

[44] 10k Human PBMCs, Multiome v1.0, Chromium Controller. https://www.10xgenomics.com/resources/datasets/10-k-human-pbm-cs-multiome-v-1-0-chromium-controller-1-standard-2-0-0, 2021.

[45] PBMC from a Healthy Donor - No Cell Sorting (3k). https://www.10xgenomics.com/resources/datasets/pbmc-from-a-healthy-donor-no-cell-sorting-3-k-1-standard-2-0-0, 2021.

[46] PBMC from a Healthy Donor - No Cell Sorting (10k). https://www.10xgenomics.com/resources/datasets/pbmc-from-a-healthy-donor-no-cell-sorting-10-k-1-standard-2-0-0, 2021.

[47] Zhongli Xu, Elisa Heidrich-O’Hare, Wei Chen, and Richard H. Duerr. Comprehensive benchmarking of CITE-seq versus DOGMA-seq single cell multimodal omics. Genome Biology, 23(1):1–17, December 2022. ISSN 1474-760X. doi: 10.1186/s13059-022-02698-8.

[48] Tim Stuart, Avi Srivastava, Shaista Madad, Caleb A. Lareau, and Rahul Satija. Single-cell chromatin state analysis with Signac. Nature Methods, pages 1–9, November 2021. ISSN 1548-7105. doi: 10.1038/s41592-021-01282-5.

[49] Yong Zhang, Tao Liu, Clifford A. Meyer, Jérôme Eeckhoute, David S. Johnson, Bradley E. Bernstein, Chad Nusbaum, Richard M. Myers, Myles Brown, Wei Li, and X. Shirley Liu. Model-based Analysis of ChIP-Seq (MACS). Genome Biology, 9(9):R137, September 2008. ISSN 1474-760X. doi: 10.1186/gb-2008-9-9-r137.

[50] Dvir Aran, Agnieszka P. Looney, Leqian Liu, Esther Wu, Valerie Fong, Austin Hsu, Suzanna Chak, Ram P. Naikawadi, Paul J. Wolters, Adam R. Abate, Atul J. Butte, and Mallar Bhattacharya. Reference-based analysis of lung single-cell sequencing reveals a transitional profibrotic macrophage. Nature Immunology, 20(2):163–172, February 2019. ISSN 1529-2916. doi: 10.1038/s41590-018-0276-y.

[51] F. Alexander Wolf, Philipp Angerer, and Fabian J. Theis. SCANPY: Large-scale single-cell gene expression data analysis. Genome Biology, 19(1):15, February 2018. ISSN 1474-760X. doi: 10.1186/s13059-017-1382-0.

[52] Analysis, visualization, and integration of Visium HD spatial datasets with Seurat. https://satijalab.org/seurat/articles/seurat5_sketch_analysis.html, 2023.

[53] Mohammad Lotfollahi, Mohsen Naghipourfar, Fabian J Theis, and F Alexander Wolf. Conditional out-of-distribution generation for unpaired data using transfer VAE. Bioinformatics, 36(Supplement_2):i610–i617, December 2020. ISSN 1367-4803. doi: 10.1093/bioinformatics/btaa800.

[54] Mohammad Lotfollahi, Anastasia Litinetskaya, and Fabian J. Theis. Multigrate: Single-cell multi-omic data integration, March 2022.

[55] Fabian Pedregosa, Gaël Varoquaux, Alexandre Gramfort, Vincent Michel, Bertrand Thirion, Olivier Grisel, Mathieu Blondel, Peter Prettenhofer, Ron Weiss, Vincent Dubourg, Jake Vanderplas, Alexandre Passos, David Cournapeau, Matthieu Brucher, Matthieu Perrot, and Édouard Duchesnay. Scikit-learn: Machine Learning in Python. Journal of Machine Learning Research, 12(85):2825–2830, 2011. ISSN 1533-7928.

[56] Ilya Korsunsky, Nghia Millard, Jean Fan, Kamil Slowikowski, Fan Zhang, Kevin Wei, Yuriy Baglaenko, Michael Brenner, Po-ru Loh, and Soumya Raychaudhuri. Fast, sensitive and accurate integration of single-cell data with Harmony. Nature Methods, 16(12):1289–1296, December 2019. ISSN 1548-7105. doi: 10.1038/s41592-019-0619-0.

[57] Maren Büttner, Zhichao Miao, F. Alexander Wolf, Sarah A. Teichmann, and Fabian J. Theis. A test metric for assessing single-cell RNA-seq batch correction. Nature Methods, 16(1):43–49, January 2019. ISSN 1548-7105. doi: 10.1038/s41592-018-0254-1.

[58] Peter J. Rousseeuw. Silhouettes: A graphical aid to the interpretation and validation of cluster analysis. Journal of Computational and Applied Mathematics, 20:53–65, November 1987. ISSN 0377-0427. doi: 10.1016/0377-0427(87)90125-7.

[59] Ritambhara Singh, Pinar Demetci, Giancarlo Bonora, Vijay Ramani, Choli Lee, He Fang, Zhijun Duan, Xinxian Deng, Jay Shendure, Christine Disteche, and William Stafford Noble. Unsupervised manifold alignment for single-cell multi-omics data. In Proceedings of the 11th ACM International Conference on Bioinformatics, Computational Biology and Health Informatics, BCB ‘20, pages 1–10, New York, NY, USA, September 2020. Association for Computing Machinery. ISBN 978-1-4503-7964-9. doi: 10.1145/3388440.3412410.

[60] Kevin E. Wu, Kathryn E. Yost, Howard Y. Chang, and James Zou. BABEL enables cross-modality translation between multiomic profiles at single-cell resolution. Proceedings of the National Academy of Sciences, 118(15), April 2021. ISSN 0027-8424, 1091-6490. doi: 10.1073/pnas.2023070118.

[61] William M. Rand. Objective Criteria for the Evaluation of Clustering Methods. Journal of the American Statistical Association, 66(336):846–850, 1971. ISSN 0162-1459. doi: 10.2307/2284239.

